# Deterministic evolution and stringent selection during pre-neoplasia

**DOI:** 10.1101/2022.04.09.487529

**Authors:** Kasper Karlsson, Moritz J. Przybilla, Eran Kotler, Aziz Khan, Hang Xu, Kremena Karagyozova, Alexandra Sockell, Wing H. Wong, Katherine Liu, Amanda Mah, Yuan-Hung Lo, Bingxin Lu, Kathleen E. Houlahan, Zhicheng Ma, Carlos J. Suarez, Chris P. Barnes, Calvin J. Kuo, Christina Curtis

## Abstract

The earliest events during human tumor initiation, while poorly characterized, may hold clues to malignancy detection and prevention^1^. Here we model occult pre-neoplasia by bi-allelically inactivating *TP53*, a common early event in gastric cancer, in human gastric organoids. Causal relationships between this initiating genetic lesion and resulting phenotypes were established using experimental evolution in multiple clonally derived cultures over two years. *TP53* loss elicited progressive aneuploidy, including copy number alterations and structural variants prevalent in gastric cancers, with evident preferred orders. Longitudinal single cell sequencing of *TP53* deficient gastric organoids similarly indicates progression towards malignant transcriptional programs. Moreover, high-throughput lineage tracing with expressed cellular barcodes demonstrates reproducible dynamics whereby initially rare subclones with shared transcriptional programs repeatedly attain clonal dominance. This powerful platform for experimental evolution exposes stringent selection, clonal interference, and a striking degree of phenotypic convergence in pre-malignant epithelial organoids. These data imply predictability in the earliest stages of tumorigenesis and reveal evolutionary constraints and barriers to malignant transformation with implications for earlier detection and interception of aggressive, genome instable tumors.

## Main

In rapidly adapting asexual populations, including microbes tumors, multiple mutant lineages often compete for dominance^2^. These complex dynamics determine the outcomes of evolutionary adaptation but are difficult to observe *in vivo*. Experimental evolution has yielded fundamental insights into clonal dynamics in microbes, enabling characterization of mutant clones and their fitness benefits^3, 4^. The same forces of mutation and selection fuel clonal expansions in somatic cells during aging, contributing to malignancy, but their dynamics are poorly understood^5–7^.

Cancers arise from a mutated cell that undergoes premalignant clonal expansion while accruing additional mutations. These mutations can spread in phenotypically normal tissues prior to apparent morphological changes, with aneuploidy and driver mutations preceding cancer diagnosis by years^5, 8, 9^. Identifying the causes of and barriers to malignant transformation requires characterization of the molecular phenotypes that precede this event in a tissue- specific manner. However, repeated sampling of healthy or pre-neoplastic tissue is impractical, and thus evolutionary dynamics have been inferred from sequencing data^5, 6, 10^. For example, we inferred stringent subclonal selection in pre-malignant Barrett’s esophagus (BE), whereas matched adenocarcinomas largely exhibited neutral evolution^6^, presumably due to rapid growth after transformation and diminishing returns epistasis^11^. Despite these insights, the order of somatic alterations and patterns of clonal expansion that precede transformation are obscured in established cancers^5, 12^, necessitating new approaches to empirically measure pre-malignant evolution.

Gastric cancer (GC), the fourth-leading cause of cancer mortality worldwide, lacks routine screening albeit its long lead times contributing to late diagnoses, poor prognosis and limited treatment options^13, 14^. Therefore, it is crucial to identify the molecular determinants of GC and its non-obligate precursor, intestinal metaplasia (IM), which is poorly characterized compared to precursor lesions in the adjacent esophagus (BE)^15–17^. The utility of forward- genetic and GC organoids as pre-clinical models has been established^18–21^, however, to bypass nascent progression and accelerate transformation, combinatorial hits were engineered^19, 21^.

Here, we model tumorigenesis from the “bottom up”, using CRISPR/Cas9-engineered human gastric organoids (HGOs) to identify causal relationships between initiating genetic insults and resultant genotypes and phenotypes. Since *TP53* inactivation is a common early event preceding numerical and structural chromosomal abnormalities (aneuploidy) in chromosomal instable (CIN) GC^19, 22, 23^, we use non-malignant HGO as a *tabula rasa* to study pre-neoplasia induced by *TP53* deficiency over a two-year time span. HGOs are ideal for this task as they recapitulate cellular attributes of *in vivo* models, including three-dimensional tissue structure, multi-lineage differentiation and disease pathology^21^.

While *TP53* is altered in >70% of CIN GCs^22, 23^, its ability to elicit aneuploidy, a hallmark of most solid cancers, has been controversial and appears tissue dependent^24–27^. Moreover, the extent to which specific copy number alterations (CNAs) are selectively advantageous, and their tumorigenic impact is largely unknown^28, 29^. We chart genotype to phenotype maps of gastric pre-neoplasia upon *TP53* inactivation in multiple HGO cultures and demonstrate that these models recapitulate genomic hallmarks of gastro-esophageal tumorigenesis, including the multi- hit, temporal and repeated acquisition of CNAs and structural variants (SVs), accompanied by progression towards malignant transcriptional states. Prospective lineage tracing with linked single cell expression profiles delineates early clonal dynamics, revealing extensive clonal interference, stringent selection, and rapid adaptation, underpinned by temporal genomic contingencies and phenotypic convergence. Our findings highlight the power of experimental evolution in human organoids to investigate occult pre-neoplastic processes and the repeatability of somatic evolution.

### *TP53*^-/-^ induces CNAs in defined orders

To model tumor initiation in CIN GC, we established HGOs from non-malignant tissue from three human donors undergoing gastrectomy and introduced bi-allelic *TP53* frameshift mutations via CRISPR/Cas9, resulting in an inactive gene product (**Fig. 1a, Extended Data Fig. 1, Supplementary Fig. 1-3, Supplementary Table 1-2, Methods**). From each donor (D1-3), 3 independent clonally-derived *TP53^-/-^* cultures (C1-3) were established, yielding 9 cultures for long-term propagation, 5 of which were each split into 3 replicates (R1-3) for cellular barcoding studies (n=24 cultures). Another ‘hit’ in the *APC* tumor suppressor, a Wnt pathway negative regulator altered in 20% of CIN GC (**Extended Data Fig. 2a**), was concurrently engineered in cultures 2 and 3 from donor 3 (referred to as D3C2 and D3C3, **Supplementary Fig. 1,3**) to examine the evolutionary consequence of dual tumor suppressor inactivation. Clonal status of CRISPR-edited sites was verified via Sanger sequencing and confirmed by WGS at multiple time points (**Supplementary Fig. 1-3**). Throughout, we refer to time as days after *TP53* deficiency was engineered and we group *TP53*^-/-^ and *TP53^-/-^, APC^-/-^*cultures unless otherwise specified.

**Fig. 1.**
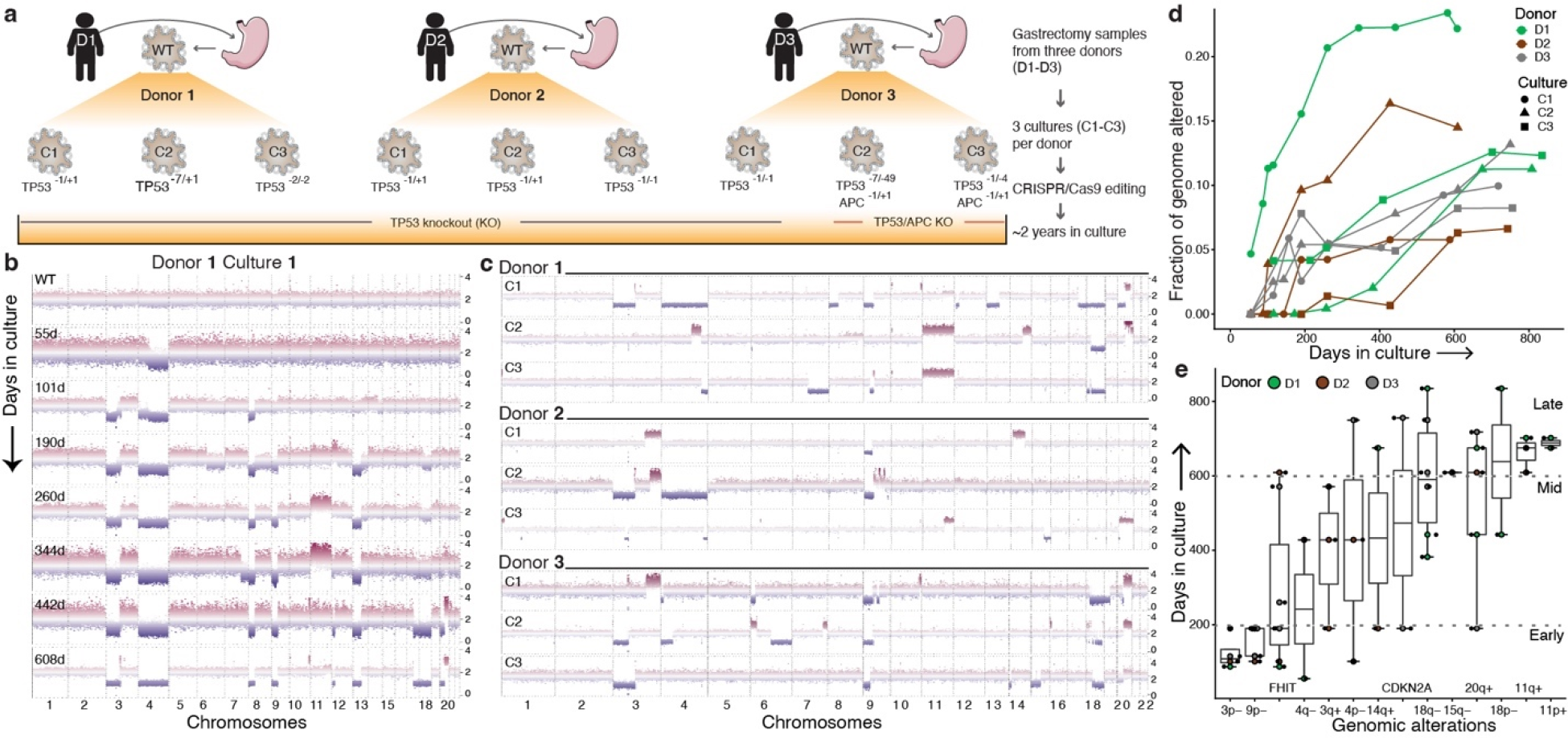
*TP53* deficiency in gastric organoids (HGOs) induces aneuploidy and gastric cancer associated copy number alterations (CNAs) along a defined temporal order. a, Schematic overview of HGO establishment and generation of *TP53^-/-^* and *TP53^-/-^,APC^-/-^*cultures via CRISPR/Cas9 editing (Methods). b, Genome-wide CNA profiles in D1C1 at multiple time points assessed through sWGS. Normalized read counts across 50kb genomic windows for each timepoint. c, CNA profiles for the 9 organoid cultures sampled between days 588-835. d, Fraction Genome Altered (FGA) over time for each culture. e, Time of appearance (days in culture) of persistent arm-level CNAs (or alterations in *FHIT* and *CDKN2A*) in *TP53/APC*- deficient organoids (alterations that become extinct are not considered). The prevalence of these alterations in GC is summarized in Extended Data Fig. 2a,b. Boxes show inter-quartile range (IQR), center lines represent the median, whiskers extend by 1.5 × IQR.

We first asked whether *TP53* deficiency elicits aneuploidy, measured as numeric and/or structural chromosomal abnormalities (genome instability). To investigate CNAs, we sequenced the 9 cultures (shallow whole-genome sequencing, sWGS; median 0.2x coverage) at up to 11 timepoints, spanning *Early* (0-200 days), *Mid* (∼200-400 days), *Late* (∼400-900 days) intervals (**Extended Data Fig. 1, Supplementary Table 3**). *TP53* deficient organoids progressively acquired CNAs, first accruing chromosome-arm level losses, followed by copy number gains (**Fig. 1b**). In contrast, wild-type (WT) gastric organoids remained genomically stable long-term (13-26 passages, **Supplementary Fig. 4-6**). Cultures from the same donor exhibited variable CNA patterns, suggesting that genetic background does not wholly constrain subsequent alterations (**Fig. 1c, Supplementary Fig. 4-6**). Despite this variability, CNAs prevalent in the TCGA GC cohort were recurrently altered in *TP53^-/-^*cultures, including loss of chromosome (chr) 3p, 9p, 18q and gain of 20q (**Extended Data Fig. 2**) ^22^. Additionally, arm-level CNAs present in two or more *TP53*^-/-^ HGOs were enriched in gastric and esophageal cancers, but not other tumor types (p<0.05, two-sided Wilcoxon rank-sum test, **Extended Data Fig. 2c**). Thus, recurrent tissue-specific CNAs accrue in *TP53^-/-^* HGOs. During the experiment, mycoplasma was detected in early passage WT cultures and some derivative samples, and an antibiotic (normocin) was used to eliminate infections (**Methods, Supplementary Fig. 7a-f**). Accordingly, replicate experiments were performed in mycoplasma-free conditions and demonstrate that mycoplasma infection was not associated with CNAs or other molecular features (**Extended Data Fig. 1, Methods, Supplementary Fig. 7g, 8**).

Across all cultures the fraction of genome altered (FGA), a measure of aneuploidy, increased over time at varying rates and plateaued around day 600 (**Fig. 1d**, **Methods**). For example, D1C1, which accrued early arm-level alterations, exhibited >20% FGA by day 260, compared to a ∼5% median FGA across all cultures at similar timepoints. *TP53^-/-^* and *TP53^-/-^*/*APC^-/-^*cultures exhibited comparable FGA at final time points (average 11.3% and 10.7%, respectively), consistent with expectation that *APC* loss does not fuel gastric cell aneuploidy. In several cultures, FGA decreased over an interval due to clonal extinction (D3C3 d190 vs 442; D2C2 d190 vs 260) (**Fig. 1d, Supplementary Fig. 5,6**). As expected, the FGA was lower in *TP53^-/-^* HGOs than CIN GCs (median FGA 34.5% in TCGA, per cBioPortal).

Investigation of the temporal onset of arm-level and focal CNAs in *TP53^-/-^* HGOs revealed preferred orders (**Fig. 1e, Supplementary Table 3**). Specifically, loss of chr9p and chr3p repeatedly occurred (across donors and cultures) within 200 days but seldom later, suggesting a period during which these alterations were particularly advantageous. Chr9p deletion spans the *CDKN2A* tumor suppressor commonly altered in the CIN subgroup of gastric (∼41%, **Extended Data Fig. 2a**) and esophageal (∼74%) adenocarcinomas, and co-occurs with *TP53* alterations^19, 22^. Indeed, *CDKN2A* loss signals the initiation of BE progression to dysplasia and esophageal adenocarcinoma^30^ and GC pre-malignancy^19^. Deeper sequencing confirmed bi- allelic loss of *CDKN2A* via focal deletion (D3C1, D3C3) or truncating mutations in p16 (*INK4A*) along with heterozygous loss (D1C3) (**Supplementary Fig. 9, Supplementary Table 3**). Similarly, deletion of the FHIT/FRA3B protein encoded on chr3p commonly occurred early in *TP53^-/-^* HGOs (median 190 days) (**Fig. 1e, Supplementary Fig. 10**, **Supplementary Table 3**). A genome caretaker, *FHIT* is lost early during tumor progression, leading to deoxythymidine triphosphate depletion, replication stress and DNA breaks^31^. Notably, 12% of CIN GCs harbor *FHIT* alterations (**Extended Data Fig. 2a**). Although CDKN2A and FHIT deletions are insufficient for malignant transformation^15, 32^, their recurrent early loss during *in vitro* evolution and in GCs implies a role in tumor initiation. Additional GC-associated CNAs include loss of chr18q and gain of chr20q, which consistently occurred late (∼600 days). Such late alterations may reflect dynamic selective pressures from increased fitness or new evolutionary paths enabled by earlier alterations^4^. These data demonstrate that *TP53* loss facilitates aneuploidy in gastric cells and accrual of tissue-specific CNAs in a defined order.

### Selection and clonal interference

We next sequenced (WGS, mean coverage 26x) five *TP53^-/-^* cultures at multiple timepoints (**Fig. 2a, Extended Data Fig. 3a, Supplementary Table 4**). This confirmed bi-allelic *TP53* and *APC* inactivation at CRISPR target sites (**Supplementary Fig. 2**) and revealed an increase in the weighted genome instability index (wGII), the fraction of genome with loss of heterozygosity (LOH), as well as focal deletions and amplifications during prolonged culture (**Fig. 2a**, **Extended Data Fig. 3b, Supplementary Table 4**). Single nucleotide variants (SNVs) and SVs also increased over time (**Fig. 2a, Extended Data Fig. 3b, Supplementary Table 4**). At late timepoints, SNV burden was higher in *TP53^-/-^/AP*C*^-/-^*(D3C2, D3C3) than in *TP53^-/-^* HGOs. Few GC associated genes were mutated across donors (**Extended Data Fig. 2e**).

**Fig. 2.**
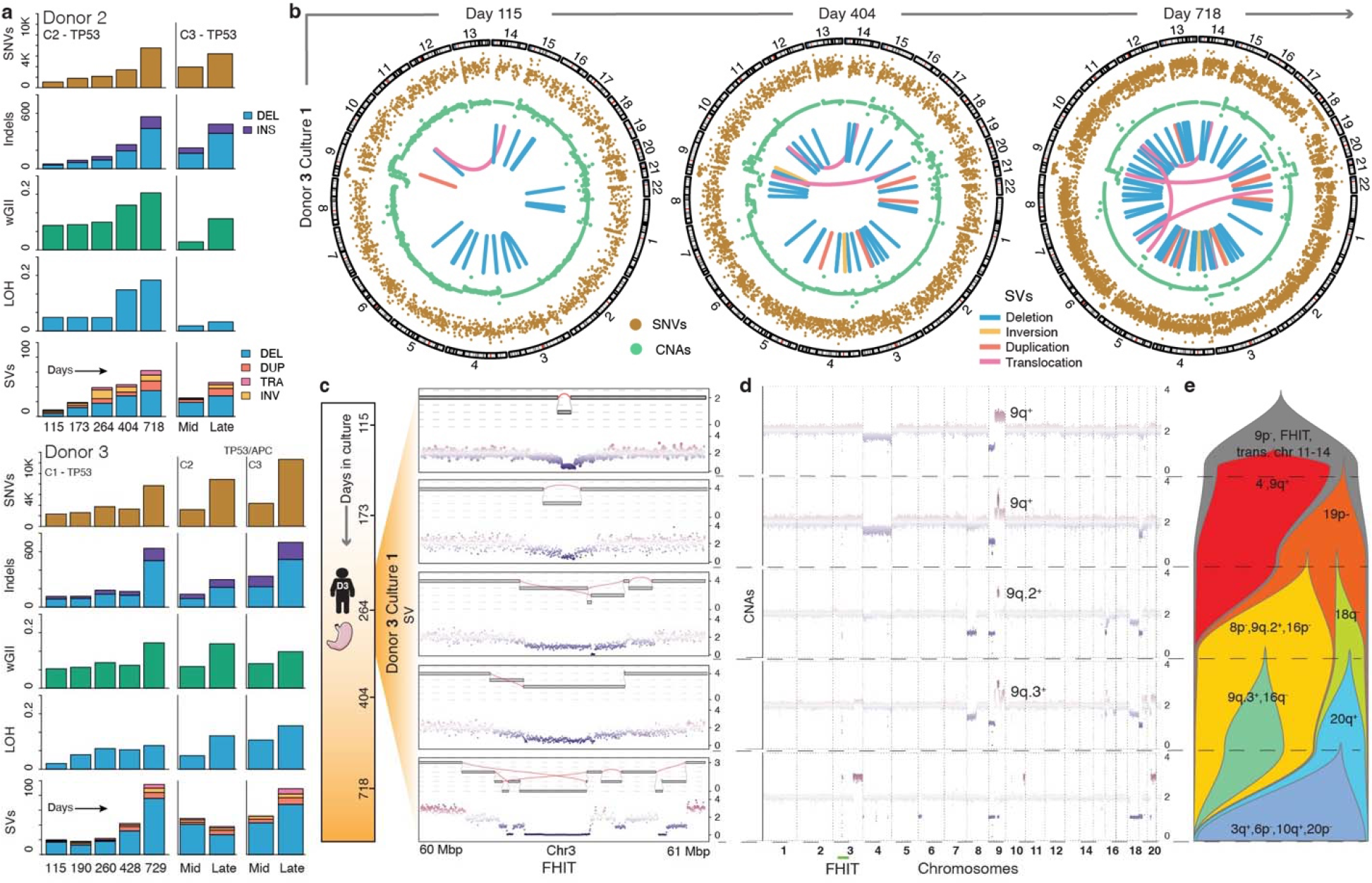
*TP53* deficiency elicits subclonal copy number evolution, SVs, and clonal interference. **a,** Burden of different classes of somatic genomic alterations in *TP53^-/-^* and *TP53^-/-^, APC^-/-^*HGOs (relative to wild-type, WT) over time, as assessed through longitudinal WGS of individual cultures at the specified time point (*Mid* – d296, *Late* d743-756). **b,** Circos plots for D3C1 illustrates increasing genomic instability and complexity with time. Classes of alterations shown include: SNVs (adjusted variant allele frequencies), copy number alterations (logR) and SV consensus calls (Methods). **c,** Evolution of rigma-like SVs at the *FHIT* fragile site on chr3p. Zoomed in view of a 1Mb region in the *FHIT* locus. Reconstructed SVs are shown in the upper panel, with corresponding CNA profiles in the lower panel. **d,** Longitudinal CNA profiles for D3C1. **e,** Fishplot schematic for D3C1 illustrates subclonal CNA evolution, clonal interference and extinction. Subclone frequencies (x-axis) are determined based on the CNAs visualized in d (Methods).

Several regions of densely clustered mutations (hypermutation) were noted, including the *FHIT* fragile site in all sequenced cultures (**Supplementary Fig. 11-13**, **Supplementary Table 3**). WT cultures exhibited simple focal *FHIT* deletions at late passages, likely due to clonal expansion of an initially rare event, and suggestive of somatic mosaicism (**Supplementary Fig. 11a-b, 12a-b**). Single base substitution (SBS) 1, 5 and 40 which are ubiquitous and implicated in aging and cancer were the most prevalent mutational signatures^33^. However, by the *Late* time point, D3C2 developed SBS 17a/b (**Extended Data Fig. 3b, Supplementary Table 4**) which is prevalent in gastro-esophageal carcinomas and progressing BE lesions^32^.

All classes of alterations accumulated in evolved *TP53^-/-^* HGOs, but SVs were particularly notable with non-clustered and simple clustered rearrangements dominating at early timepoints, followed by complex clusters (≥10 rearrangements) involving deletions, inversions and translocations over time (**Fig. 2a, Extended Data Fig. 3c,d**). This is exemplified in D3C1 which accrued multiple inter-chromosomal rearrangements (**Fig. 2b**). While such complex SVs are seldom reported in normal tissues, they are prevalent in progressing BE^16^. SV burden increased dramatically in *TP53^-/-^*HGOs between *early* and *late* timepoints (median change of 148%), exceeding by >3-fold the change in SV burden (45%) between endoscopies in BE patients harboring bi-allelic *TP53* inactivation who subsequently *progressed* to esophageal adenocarcinoma (average 2.2 years, range 0.65-6.16)^16^. In contrast, BE *non-progressors* (lacking *TP53* bi-allelic inactivation) had low and stable SV burden between endoscopies (**Extended Data Fig. 3e**).

The *FHIT* locus frequently harbored complex SVs, including deletion chasms at fragile sites (rigma), as reported in GC and BE^34^ (**Fig. 2c, Extended Data Fig. 4a, Methods**). We traced the genesis of rearrangements at the *FHIT* in D3C1, starting from a small deletion at day 115, culminating in rigma by day 264. The subclone harboring this rearrangement was lost (**Fig. 2d-e**, yellow subclone) and a separate subclone (blue) with a distinct *FHIT* rigma emerged and persisted, suggesting convergent evolution. Thus, rearrangements with multiple junctions evolve over several generations, not as a single event as previously proposed^35^. Similar events evolved in other cultures, including a chr3 and chr9 translocation (**Extended Data Fig. 4b-f, Supplementary Fig. 14a-b**). Despite these rearrangements, overall genomic content remained diploid as confirmed by flow cytometry (**Supplementary Fig. 14c, Supplementary Table 4**).

Clonal competition and extinction were investigated by determining subclonal populations from CNA profiles (via bulk WGS) across five timepoints for D3C1 and D2C2. By day 115, D3C1 had acquired numerous deletions (9p, *FHIT*) and several SVs, including a persistent chr11-chr14 translocation. Over 600 days, multiple CNA-defined subclones increased in frequency prior to extinction (**Fig. 2d-e**, **Supplementary Table 5**). For example, a chr4^-^, 9q^+^ subclone arose early but disappeared by day 264, outcompeted by a chr19p^-^ subclone that later acquired chr8p^-^, 9q.2^+^, 16p^-^ alterations and remained dominant until day 404. This subclone was ultimately outcompeted by one with chr18q loss, which acquired gain of chr20q, a recurrent late event in multiple cultures. Thus, some clones fix and achieve dominance while others reach substantial frequencies before going extinct, presumably due to clonal interference. Distinct CNA subclones coexisted for extended durations (∼140 days), suggesting comparable fitness (*e.g.,* chr8p^-^, 9q.2^+^,16p^-^ and 18q^-^ subclones) and intermittent periods of clonal competition and stasis as seen in other cultures (**Extended Data Fig. 4c,d**). These data expose stringent selection and pervasive clonal interference in premalignant epithelial populations.

### Transcriptional changes following *TP53*^-/-^

Phenotypic and transcriptional changes during *in vitro* evolution were evaluated based on growth dynamics of *TP53^-/-^* HGO cultures and single-cell RNA sequencing (scRNA-seq) at *Early*, *Mid* and *Late* time points (**Fig. 3a, Extended Data Fig. 1, 3a, Supplementary Table 2**; **Methods**). We investigated changes in cell proliferation by fitting a LOESS regression model to cell numbers at each passage, using growth derivative and fold change as a surrogate for fitness. Higher growth derivatives were observed at *Late* or *Mid* versus *Early* time points (**Fig. 3b, Extended Data Fig. 5a**); using raw cell numbers yielded similar results (p=0.003, two-way repeated-measures ANOVA; **Extended Data Fig. 5b**). scRNA-seq of the 12 cultures (7 *TP53*^-/-^ 2, *APC/TP53*, 3 WT), yielded 31,606 cells for analysis following quality control (**Fig. 3c**, **Extended Data Fig. 6, 7a,b**, **Supplementary Table 6; Methods**). Normal gastric tissue markers were expressed in WT HGOs from the three donors, including pit mucosal cell markers (PMCs: *MUC5AC*, *TFF1*, *TFF2*, *GKN2*) in D1, enterocyte markers (*FABP1*, *FABP2*, *ANPEP*, *PHGR1*, *KRT20*) in D2 and gland mucosal cell (GMCs) markers (*MUC6*, *PGC*, *TFF2*, *LYZ*) in D3, but were heterogeneous in *TP53*^-/-^ HGOs (**Fig. 3c-e, Extended Data Fig. 6, 7, Supplementary Table 6**).

**Fig. 3.**
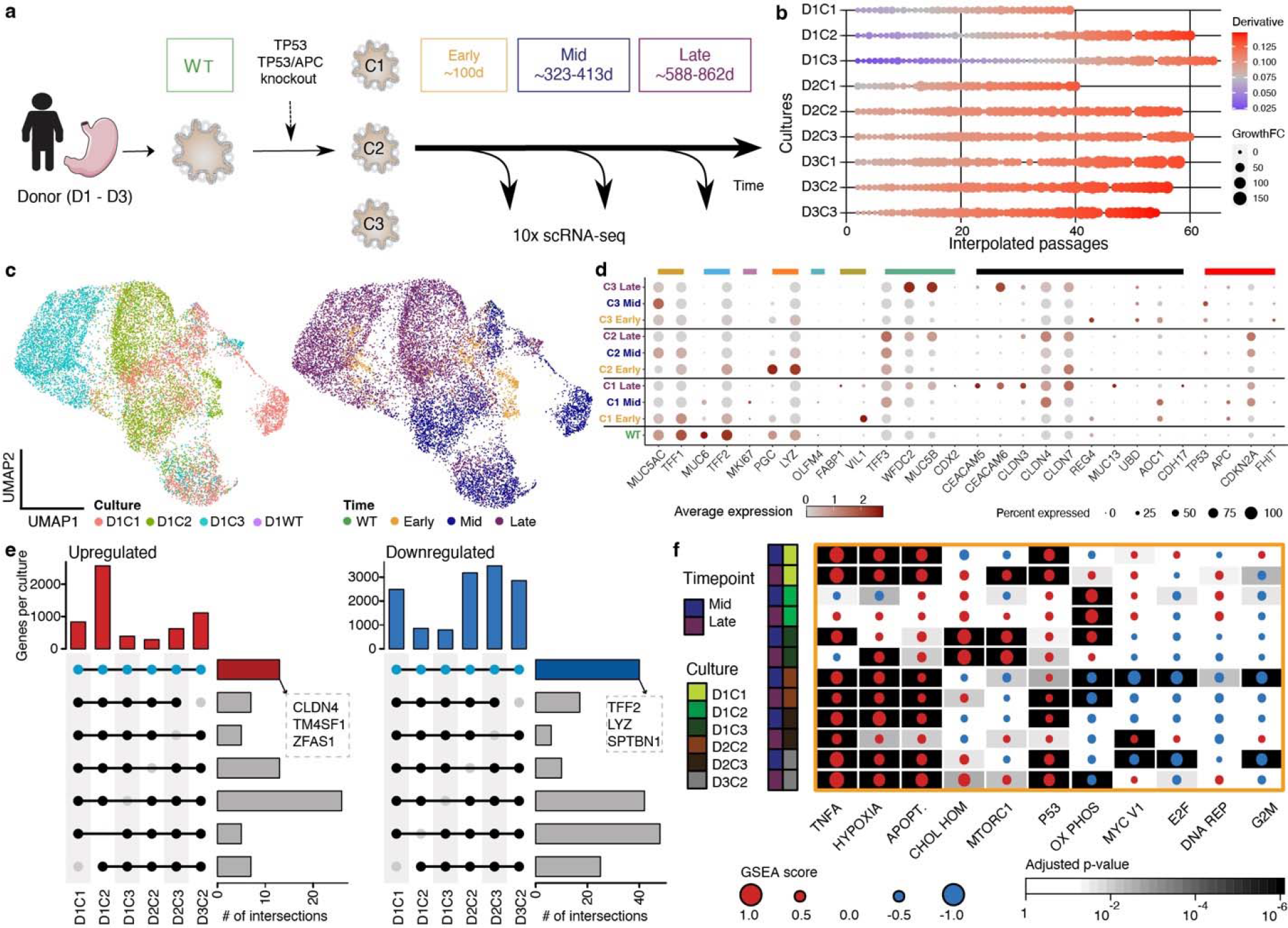
Transcriptional deregulation in *TP53* deficient gastric organoids. **a,** Experimental overview of longitudinal single-cell RNA (scRNA-seq) profiling of gastric organoid cultures. Wild- type and replicate *TP53^-/-^* HGOs were sampled at multiple time points (*Early*, ∼100 days, orange; *Mid*, ∼320 days, blue; *Late*, ∼770 days, purple) and subject to scRNA-seq. **b,** Dotplot depicting estimated growth curve derivatives and growth fold-change (FC) from previous time point for each culture over time (interpolated passage number). **c**, UMAP visualizations colored according to culture (left) and time point (right) for Donor 1 depicting 13,984 cells. **d**, Dotplot depicting the expression of selected marker genes for individual cultures and time points. Colored bars highlight marker genes associated with normal gastric and intestinal cell types, genes up-regulated in the gene expression profiling interactive analysis (GEPIA) of GC, and others of functional relevance. Pit mucosal cells: *MUC5AC*, *TFF1* – dark yellow; Gland mucosal cells: *MUC6*, *TFF2* – light blue; Proliferative cells: *MKI67* – purple; Neck-like cells: *PGC*, *LYZ* – orange; Mucosal stem cells: *OLFM4* – turquoise; Enterocytes: *FABP1*, *VIL1* – olive; Goblet cells: *TFF3*, *WFDC2*, *MUC5B*, *CDX2* – green; GEPIA top 12 genes: *CEACAM5*, *CEACAM6*, *CLDN3*, *CLDN4*, *CLDN7*, *REG4*, *MUC3A*, *MUC13*, *PI3*, *UBD*, *AOC1*, *CDH17* – black; Other: *TP53*, *APC*, *CDKN2A*, *FHIT* – red. **e,** Upset plot representing shared differentially up- (left) and down-regulated genes (right) across donors and cultures. **f,** GSEA heatmap for MsigDB Hallmark gene sets showing the most significantly altered pathways for each culture (Kolmogorov-Smirnov statistic, Benjamini-Hochberg adjusted). GSEA score is indicated (dot size) and colored according to the directionality of expression profiles (up, red; down, blue).

The mucosal-like phenotype in WT cultures, defined by mucin and TFF gene expression, was lost following *TP53^-/-^* in D1 and D3. Additionally, in D1, intestinal goblet cell-specific markers, including *TFF3*, *WFDC2* and *MUC5B*, were upregulated at the *Late* time point, as commonly seen in IM^36^. GC associated genes, including claudins (*CLDN3*, *CLDN4*, *CLDN7)* and carcinoembryonic antigens (CEA) family (*CEACAM5*, *CEACAM6)* increased in expression over time in D1 and D3. The inverse was observed in D2 cultures, plausibly due to an inflamed biopsy and the predominance of enterocytes in WT culture^14^ (**Extended Data Fig. 7a,c**). The absence of *MUC5AC* after *TP53^-/-^* and increase in *CEACAM6* expression was verified by immunofluorescence staining in D3C2 (**Extended Data Fig. 5c**).

We investigated the overlap in transcriptional features across *TP53^-/-^* HGO by intersecting significantly differentially expressed genes (DEGs) from *Early* to *Late* time points across the six cultures with scRNA-seq data. In total, 13 consistently up-regulated and 40 down- regulated genes were identified (Bonferroni corrected p<0.05, Wilcoxon Rank-Sum test, **Fig. 3e**, **Supplementary Table 6**). Upregulated genes included *CLDN4, TM4SF1*, *ZFAS1* which are implicated in GC^22, 37^, whereas those downregulated included *SPTBN1*, a cytoskeletal protein involved in TGF-β signaling^38^, and mucin production modulators *LYZ* and *TFF2*.

Lastly, we assessed pathway-level changes by gene set enrichment analysis (GSEA) of DEGs in *Late* versus *Early* as well as *Mid* versus *Early* time points (**Fig. 3f**, **Supplementary Table 6**). Several pathways were enriched across multiple cultures and donors, including upregulation of TNF-α signaling via NF-κβ as reported in CIN tumors^39^ and comparisons of GC versus normal tissue^40^ (4/6 cultures), apoptosis (5/6 cultures), and hypoxia (5/6). Downregulated pathways included MYC, E2F targets and G2M checkpoints, although these were more variable and likely reflect survival programs. Thus, despite heterogenous single gene trajectories, pathways implicated in malignancy were shared across cultures and donors.

### Emergence of malignant expression states

To identify pathologic features, we projecting HGO longitudinal scRNA-seq data onto a reference atlas comprised of normal and GC scRNA-seq^41^ (**Methods**). Restricting the reference to epithelial cells yielded 6,001 cells (1,354 normal; 4,647 tumor) assigned to distinct cell type clusters using literature-derived marker genes^40–42^. Two tumor cell clusters emerged, comprised of mucosal-like malignant and non-mucosal-like malignant cells^41^ where the latter included malignant markers *(KRT17*, *KRT7*, *LY6D*) but lacked mucosal markers (*MUC5AC*, *TFF2*, *TFF1*) (**Fig. 4a**). PMCs, GMCs, chief cells, parietal cells, enterocytes, enteroendocrine cells, goblet cells and proliferative cells were also assigned to clusters (**Fig. 4b**).

**Fig. 4.**
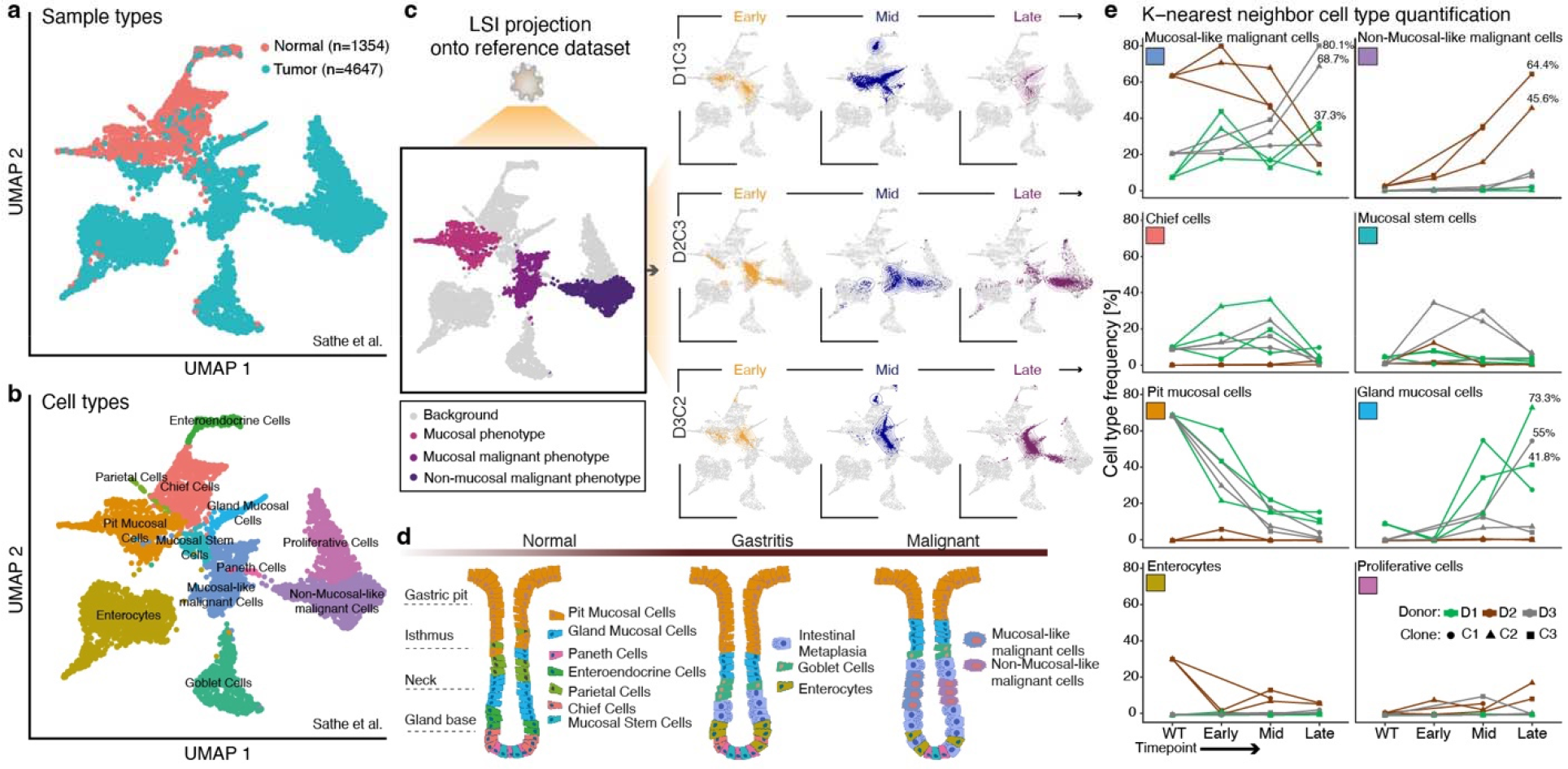
Unsupervised assessment indicates progression towards malignant transcriptional states. **a**, **b** UMAP embedding of 6,080 epithelial cells from the Sathe et al. gastric tumor-normal scRNA-seq dataset, colored according to histology (a) and assigned cell type (b). Detected cell types included pit mucosal cells (PMCs), gland mucosal cells (GMCs), chief cells, parietal cells, enterocytes, enteroendocrine cells, goblet cells and proliferative cells, as well as two types of malignant cells (mucosal-like and non-mucosal-like). **c,** Latent Semantic Indexing (LSI) projection of *TP53^-/-^* HGOs sampled at *Early* (orange), *Mid* (blue) and *Late* (purple) timepoints onto the reference dataset (left), colored by cellular phenotypes of interest, providing orientation for the LSI projection of the three HGO cultures at the specified timepoints (right). The density of projected cells is highlighted using a 2D density distribution. **d,** Schematic representation of shifts in cell populations proposed to accompany the transition from normal tissue to gastritis which can lead to IM and malignancy, adapted from ^40^. **e,** Projected cell type frequencies based on the 25 nearest neighbors in HGOs over time.

We next projected batch corrected scRNA-seq data from *Early*, *Mid* and *Late* time points individually onto the reference embedding to identify the gastric cell types that were most similar (**Fig. 4c, Extended Data Fig. 7e-h, Methods**). As the reference atlas lacked pre-neoplastic populations, cells were projected onto either normal or tumor cells states, where the majority of HGO cells mapped onto the latter. Shifts in cell states over time were evident for all cultures, some of which have been implicated in the normal-to-gastritis transition that can lead to IM and ultimately malignancy (shown schematically in **Fig. 4d**). Changes in cell type frequencies were quantified for each HGO culture by identifying the 25 nearest neighbors (NNs) in the reference population (**Fig. 4e**). An increase in mucosal-like malignant cells was observed in 3/7 cultures at the *Late* time point, with 68.7%, 80.1% and 37.3% of NNs being mucosal-like malignant cells for D3C2, D3C3 and D1C1, respectively. In contrast, for D2, mucosal-like malignant cells decreased while non-mucosal-like malignant cells increased from WT to the *Late* timepoint (D2C2 – 45.6%, D2C3 – 64.4%, NNs) (**Fig. 4e**), explaining transcriptional differences relative to D1 and D3 (**Fig. 3**). Notably, ∼30% of cells in D2WT projected near enterocytes, potentially contributing to gastritis-like features, and underlining the transcriptional similarity between enterocytes and malignant cells ^42^. WT cultures from D1 and D3 exhibited predominantly mucosal phenotypes. The decrease in mucosal gene expression suggests that the evolved *TP53* deficient HGOs are *en route* towards IM and malignancy, albeit at different rates, corroborating the supervised analyses based on specific marker genes. While our HGO cultures harbor hallmarks of CIN GC, they do not exhibit evidence of histologic transformation (**Extended Data Fig. 5d**).

### Deterministic growth of rare subclones

We next leveraged our HGO models to characterize pre-neoplastic subclonal dynamics at cellular resolution via prospective lineage tracing with high-complexity cellular barcodes. To jointly recover lineage and transcriptional states, we developed expressed cellular barcodes (ECB), which uniquely label each cell (**Supplementary Fig. 15a,b, Methods**). Five *TP53^-/-^* (D1C1, D1C3, D2C1, D2C2, D2C3) and one *TP53^-/-^*, *APC^-/-^* (D3C2) culture were transduced with ECB lentivirus between days 101-115 and evolved in parallel to the non-barcoded cultures for over one year. All cultures were Sanger sequenced at multiple time points to verify *TP53/APC* deletion clonality, resulting in exclusion of D1C3 (**Supplementary Fig. 15c,d**). Each ECB parental line was split into three replicates to evaluate the reproducibility of clonal dynamics, where outgrowth of the same subclone is assumed to reflect an *intrinsic* fitness advantage and divergent subclone dominance suggests *acquired* fitness differences (**Fig. 5a**).

**Fig. 5.**
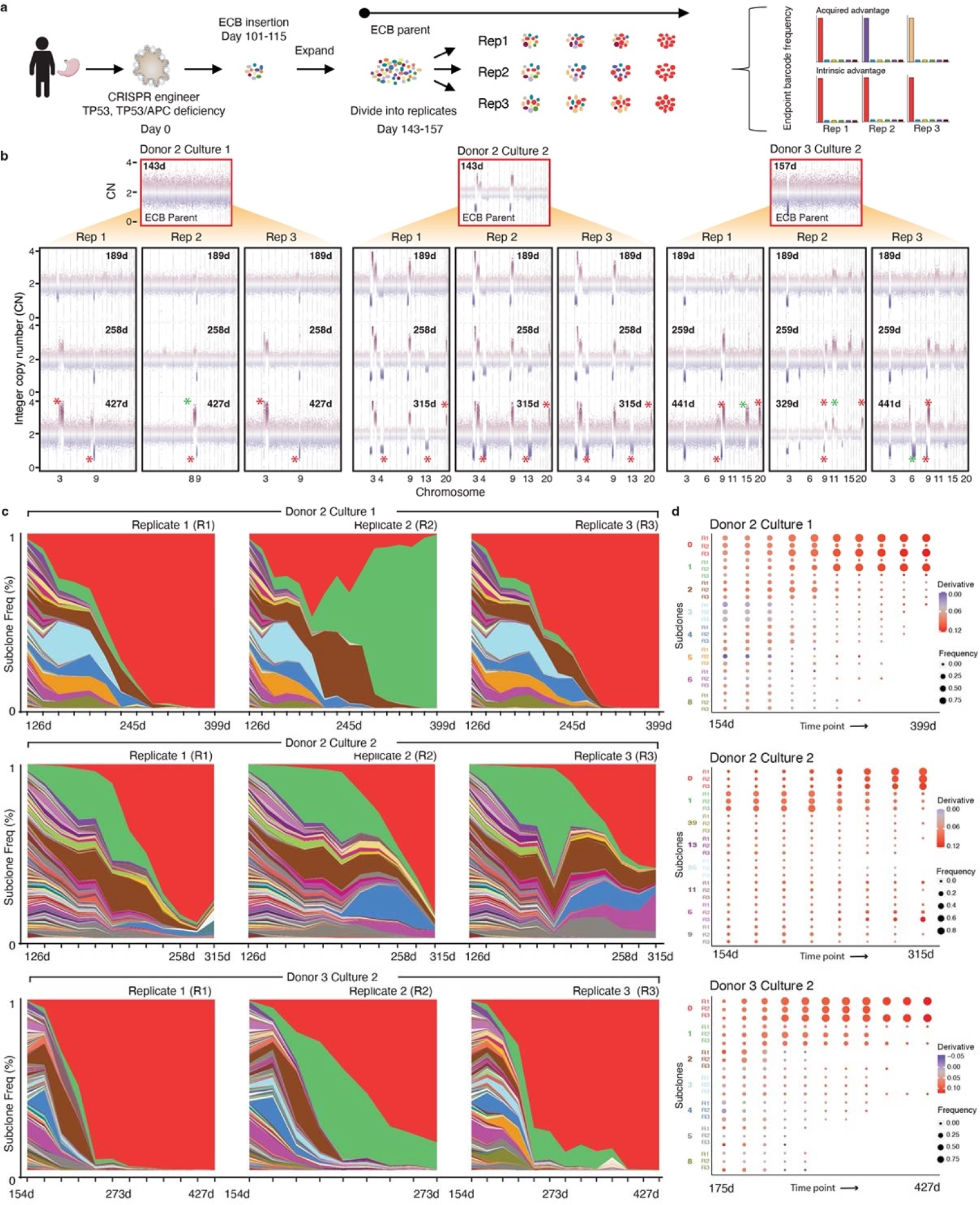
Lineage tracing reveals subclonal dynamics and deterministic outgrowth. **a,** Overview of prospective lineage tracing studies in *TP53*^-/-^ HGOs using ECBs. The ECB parental population was split into replicates, and individual cultures evolved in parallel and subject to longitudinal barcode sequencing, revealing subclonal dynamics and assessment of intrinsic or acquired fitness advantages amongst replicate cultures. **b,** CNA profiles were assessed by sWGS prior to introduction of the ECB in the parental line and across replicate ECB cultures at multiple time points. Red stars denote CNAs present in at least two replicates but not in the parental population; green stars denote CNAs unique to one replicate. Only chromosomes that harbor newly arising CNAs (not present in the parental population) are numbered for simplicity. **c,** Mueller plots depict ECB frequencies (assessed by barcode sequencing) over time where each color represents a distinct subclone in each replicate. Note that for D3C2-R2 the barcode was lost ∼day 273. **d,** Dotplots indicate ECB subclone frequency (size) and estimated growth curve derivative per subclone (color).

Longitudinal sWGS of these long-term ECB cultures demonstrated striking reproducibility at the genomic level with recurrent CNAs shared across replicate cultures (**Fig. 5b, Extended Data Fig. 8, Supplementary Fig. 16**). For example, in D2C2, new CNAs emerged around day 258 (loss of chr4q and chr13; gain of chr20q) across all three replicates. In D2C1, replicate 2 (D2C1R2) gain of chr8q was detected by day 258 and persisted (**Fig. 5b**) but was mutually exclusive with gains of chr3q in R1 and R3. In contrast, CNAs in different cultures from the same donor were more variable (**Fig. 1c**).

Through DNA sequencing of ECBs at regular intervals, we estimated the relative abundances of subclones over time and constructed Mueller plots to visualize clonal dynamics (**Fig 5c**). Colors were assigned to barcodes based on subclone frequencies across replicates within a culture and the highest frequency subclone was colored red. For example, the red band in D2C1R1 represents the same barcoded subclone as in D2C1R2 and D2C1R3. For each culture (except D2C1R2) the same (red) subclone became dominant across all replicates (**Fig. 5c**), consistent with an intrinsic fitness advantage and deterministic outgrowth (**Fig. 5a**). For D2C1 replicates R1 and R3 the red subclone became dominant, in-line with their shared CNA profiles, whereas in R2, the green subclone which acquired a chr8q gain (spanning the *MYC* oncogene), overtook the population. Intriguingly, the brown and green clone expanded concomitantly before going extinct, suggesting their mutual dependence.

Of note, subclone frequency correlations over time across replicates was generally high, reflecting similar subclonal dynamics within a culture and similar patterns across cultures (**Extended Data Fig. 8**). Especially striking was the remergence of the blue and purple subclones in D2C2 R2 and R3 at ∼200 days (**Fig. 5c**). By constructing subclone specific growth curves and estimating their derivatives, we found that winning subclones had high initial fitness and increased in proliferative capacity over time (**Fig. 5d, Methods**). Thus, lineage tracing reveals reproducible dynamics across replicate cultures with adaptive lineages sweeping rapidly to fixation and dominant clones comprising 75% (median across cultures) of the population by day 144 post ECB transduction (**Supplementary Table 7**). These patterns are reminiscent of rapid adaptation in isogenic microbial populations attributable to standing variation in the initial population^43, 44^.

### Molecular features of winning subclones

To investigate the targets of selection and how they change through time and across populations, we leveraged the ECBs which jointly capture lineage and transcriptional states in individual cells. Specifically, we sought to characterize the molecular features of ‘winning’ subclones that dominated the population after prolonged evolution, by performing scRNA-seq for several donors and replicates at selected timepoints when the population was heterogeneous. For D2C2R2, which was sampled at day 173, 1,284 cells passed QC and we identified 20 subclones with at least 10 cells, all of which were among the top 38 most frequent ECBs based on barcode sequencing. Arm-level CNAs were inferred from the scRNA-seq data using *inferCNV* (**Methods**), revealing numerous subclone-specific CNAs (**Fig. 5b, 6a, Methods**). Reassuringly, aggregate CNA landscapes were concordant with WGS data, and single cell DNA-sequencing showed similar profiles and frequencies to subclone-specific CNAs inferred from scRNA-seq (**Supplementary Fig. 17, Methods**). A detailed examination of this replicate (D2C2R2) revealed complex evolutionary dynamics amongst coexisting subclones. Most cells comprising the winning subclone (ECB-0, red) acquired chromosome 3p^-^, 3q^+^, 9p^-^ and 9q^+^ alterations early, as these events were clonal or nearly clonal in the parent population at day 143 (**Fig. 6a,b**). A subpopulation within ECB-0 (termed “0a”) additionally acquired chr4q^-^ and chr20q^+^ and ultimately became dominant with these alterations present in ∼90% of the population at day 315 (**Fig. 5b, Supplementary Fig. 18**). Similar dynamics were seen across all replicate cultures where winning subclones contained a nested CNA-defined subclone (**Extended Data Fig. 9-11**). These patterns may reflect a ‘rich-get-richer’ effect where fitness advantages acquired early drive clonal expansions, thereby increasing the chance of additional alterations that fuel growth^45^.

**Fig. 6.**
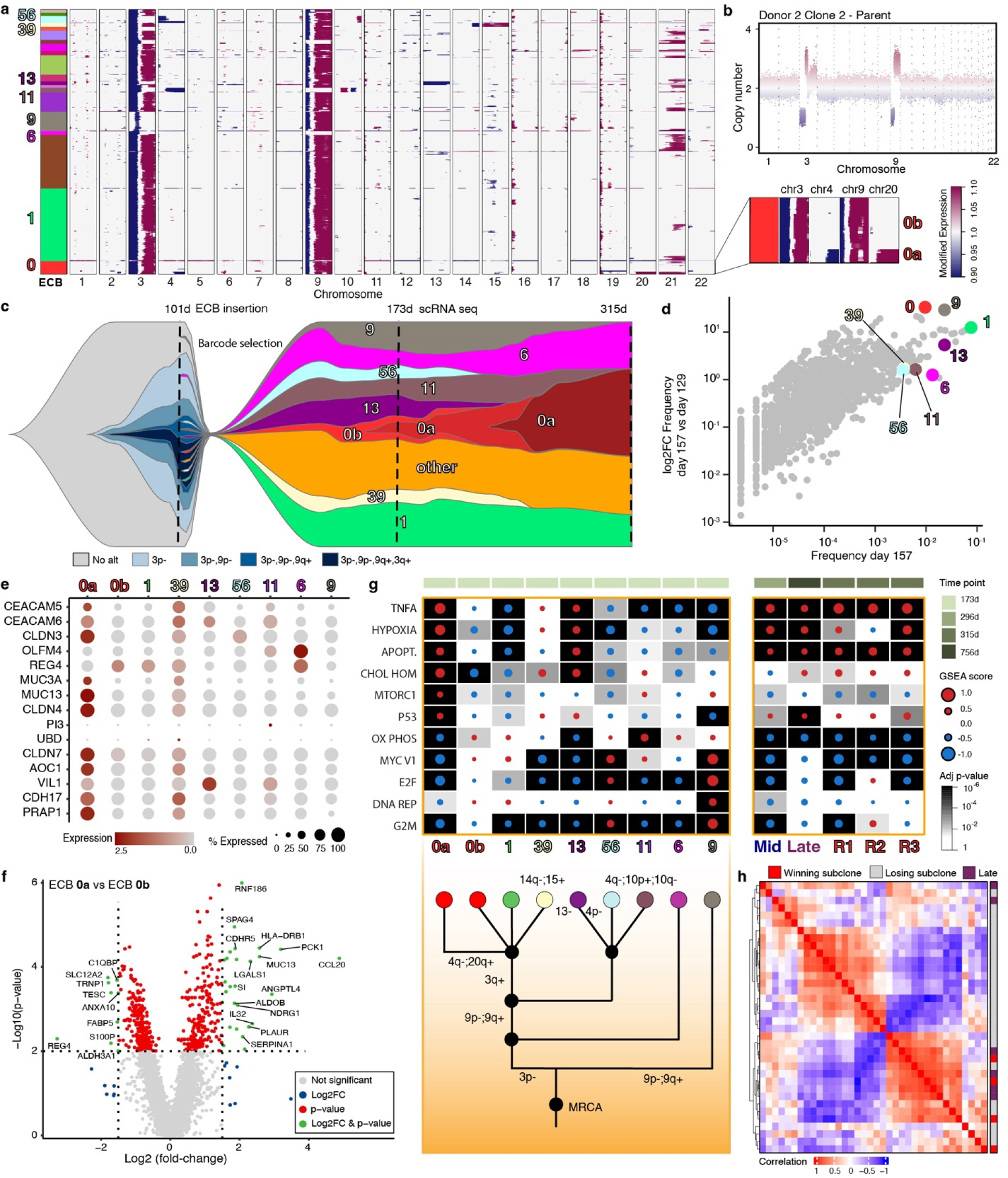
Genotype to phenotype mapping defines molecular determinants of winning subclones. **a,** Inferred CNA heatmap from scRNA-seq data for D2C2, Replicate 2 (D2C2R2) at day 173, where each row is a cell. The color bar at the left indicates the ECB each cell maps to. Numbered barcodes were selected for further investigation. Inset shows a subpopulation within ECB-0 with additional CNAs, termed “0a”, while ECB-0 parent subclone is termed “0b”. **b,** CNA profile for the D2C2 parental population (also shown in Fig 5b). **c,** Fishplot schematic illustrating the link between lineage (ECBs) and CNA subclones. To facilitate visualization, subclones of interest (denoted in a) are shown, while the remainder grouped as “*other*”, all values are log transformed. **d,** Scatterplot comparing subclone frequency at day 157 and the log2 fold change between days 129 and 157. All subclones are shown, those of interest are highlighted as in a. **e,** Dotplot showing the expression of top DEGs based on gene expression profiling interactive analysis (GEPIA) of gastric cancers. **f,** Volcano plot illustrating DEGs from the comparison of the winning subclone 0a and its parental subclone 0b. Vertical and horizontal lines correspond to absolute log2 fold change values of 1.5 and p < 0.01, respectively. **g,** GSEA heatmap from MsigDB Hallmark gene sets showing the most significantly altered pathways (Kolmogorov- Smirnov statistic, Benjamini-Hochberg adjusted) for specific subclones at day 173 (left) and later time points for the same culture (right). A manually reconstructed phylogeny is shown below. **h,** Pairwise spearman correlation between samples based on GSEA score for the top 10 most altered pathways for *Late* relative to *Early* timepoints and for D2C2 subclones.

Since successful subclones consistently acquired additional genetic diversity, we sought to investigate the functional relevance of these events, focusing on a subset of subclones with divergent CNAs. As an example, D2C2R2 consisted of at least five different CNA clones at the time of barcode insertion (**Fig. 6c, Supplementary Table 8**). Multiple instances of convergent evolution were evident *within* this culture, where subclones acquired the same CNA independently, implying stringent selection. For example, ECB-0a, ECB-11 and ECB-56 each lost variable sized regions of chr4q. ECB-9 lacked common early alterations including chr3p^-^, but subsequently acquired chr9p/q alterations. Despite the incomplete set of CNAs, ECB-9’s growth closely trailed that of the winning subclone (ECB-0, **Fig. 6d**). Convergent CNA evolution was also evident *across* cultures, where chr15 and chr20 amplifications were present in the majority of cells in D3C2 R1 at day 441, while these events plus chr11 amplification were present in R2 and R3 subclones by day 259.

Although highly fit subclones differed in genomic landscapes, we reasoned they would share transcriptional programs. Indeed D2C2R2, the winning subclone 0a (but not its parent 0b), exhibited high expression of several GC genes, including *CEACAM5*, *CEACAM6*, *CLDN3*, *CLDN4* and *CLDN7* (**Fig. 6e**). These genes were also highly expressed in winning subclones of all other replicate cultures, except for ECB-1a (green) in D2C1R2 which acquired 8q gain (**Extended Data Fig. 9e, 10d, 11d**). The winning subclone 0a (versus 0b) also upregulated GC genes, including *RNF186* which regulates intestinal homeostasis and is associated with ulcerative colitis^46^; *MUC13* that encodes a transmembrane mucin glycoprotein^47^; *CCL20,* a chemokine and candidate biomarker^48^, and *LGALS1* (galectin-1) which promotes epithelial-mesenchymal transition, invasion and vascular mimicry^49^ (**Fig. 6f, Supplementary Table 9**). GSEA analysis, comparing the winning subclone in D2C2 to all other cells, revealed up- regulation of several pathways, including TNF-α signaling via NF-κβ, as well as hypoxia, apoptosis and p53 (**Fig. 6g, Extended Data Fig. 12a, Supplementary Table 9**). These same pathways were upregulated in three barcoded replicates for D2C2 at the final time point (day 315), as well as in the non-barcoded D2C2 culture at *Mid* and *Late* time points (relative to *Early*) (**Fig. 6f**, right) and other donors/cultures (D1C1, D1C2, D1C3, D2C3, D3C2) (**Fig. 3f**). Moreover, these pathways were upregulated in winning subclones from independent barcoded donors/cultures (**Extended Data Fig. 9-11**), including the divergent subclone (ECB-1a, green) in D2C1R2 (**Fig. 5c**, **Extended Data Fig. 10f**), emphasizing their reproducibility. More generally, strong concordance between winning subclone and non-barcoded *Late* subclones was observed across the top 10 altered gene sets, irrespective of mycoplasma levels, antibiotic treatment, and other sources of biological and technical variation (**Supplementary Fig. 19, Methods**). Similarly, winning subclones clustered with *Late* cultures, which exhibited malignant transcriptional states based on the unsupervised LSI projection (D1C1, D2C2, D2C3 and D3C2) (**Fig. 6h, Extended Data Fig. 12a-b**). Notably, there was a significant difference in the activation of p53, apoptosis and TNF-a signaling via NF-κβ pathways (Fisher exact test, Bonferroni corrected p<0.05) between Late (relative to Early) and winning subclones, compared with all other subclones (**Extended Data Fig. 12c**). These data highlight convergent phenotypic evolution in which the early activation of specific pathways is selectively advantageous, canalizing cells towards malignancy.

## Discussion

Through multi-year experimental evolution of *TP53*-deficient HGO cultures, we model pre- neoplastic evolution and genotype-phenotype relationships following this common initiating insult. Remarkably, *TP53* deficiency was sufficient to recapitulate multiple hallmarks of CIN GC including aneuploidy, specific CNAs, SVs, and transcriptional programs, emphasizing the importance of cell intrinsic processes during pre-malignant evolution. Although aneuploidy propagates heterogenous evolution, our data reveal preferred orders in the acquisition of CNAs, with early loss of chr3p and 9p frequently followed by bi-allelic inactivation of *CDKN2A* and/or *FHIT* and relatively late gain of 20q. Such preferred mutational orders have been described during tumorigenesis, most notably in the colon, but the resolution of inferences from cross-sectional data or established tumors is inherently limited^12, 50^. Evolutionary phases in which deletions preceded whole-genome doubling (WGD) and subsequent amplifications were recently reported in a murine model of *KrasG12D, Trp53*-deficient pancreatic cancer, but gene or chromosome level orderings were not seen in this system^51^.

Our *TP53*^-/-^ HGOs exhibited transcriptional and genomic hallmarks of pre-malignant gastro-esophageal lesions, despite remaining histologically normal. This is consistent with significant genomic perturbation being required for even the earliest stages of gastro- esophageal carcinogenesis and the accrual of complex rearrangements years before cancer diagnosis^15, 16, 23^.

*TP53*^-/-^ HGOs appear to be on a trajectory similar to *TP53*-null BE for which the presumed cell of origin is gastric cardia^17^ and proposed biomarkers of progression to esophageal cancer include CNA acquisition, and SV burden^16, 52^. These *in vitro* models thus recapitulate occult pre- neoplasia and mirror the latency of human tumorigenesis, with additional time or *in vivo* selective pressures evidently required for malignant transformation and further features of invasive disease such as WGD or *ERBB2* amplification^22^.

The finding that *TP53* deficiency elicits a temporally defined order of genomic aberrations raises the possibility that these features may similarly predict progression to CIN GC. Future evaluation of this hypothesis will require annotated IM tissue collection with long-term follow-up. While *TP53* deficiency elicits tissue-specific alterations that may aid the detection of high-risk lesions, this constrained evolutionary state is unlikely to persist indefinitely given ensuing genome instability, emphasizing the need for earlier detection.

By jointly measuring lineage, CNAs, and transcriptional states in individual cells, we investigated the molecular basis of clonal expansions and fitness. This revealed stringent selection and reproducible subclonal dynamics across replicate cultures in which the same, initially rare, subclone fixed in the population. Pervasive clonal interference was evident amongst subclones, accompanied by intermittent periods of relative stasis, suggesting that an optimal karyotype has yet to be achieved, as reported in colorectal adenomas^53^. Further, we observed a striking degree of phenotypic convergence on common dominant pathways across cultures and donors, irrespective of mycoplasma infection and antibiotic treatment. This evolutionary reproducibility is particularly notable given these and other potential sources of technical and biological variation and implies that any such effects are evidently modest relative to the overwhelmingly dominant effect of *TP53* inactivation.

These first in-kind measurements address open questions concerning selection and determinism in clonal evolution extendable to other tissues. In the vast space of initiating insults, recurrent tissue-specific alterations can be prioritized to identify selectively advantageous alterations, temporal order constraints and convergent phenotypes. Such constraints, due to epistasis, can reveal barriers to malignant transformation and potential therapeutic targets. We anticipate that our results will advance empirical and theoretical investigations of mutation, selection and genome instability in human cells, much as the long-term evolution experiments pioneered by Lenski and colleagues decades ago continue to yield fundamental insights into microbial adaptation^3, 4^.

## Acknowledgments

The authors thank Zheng Hu, Susanne Tilk, Laura Attardi, and Ami Bhatt for helpful discussions, the Stanford University Hospital Tissue Procurement Shared Resource facility for specimen procurement and the Stanford Functional Genomics Core for assistance with sequencing. This work was supported by the National Institutes of Health (NIH) Director’s Pioneer Award: DP1CA238296 to C.C. and a National Cancer Institute (NCI) Cancer Target Discovery and Development Center (U01CA217851) to C.K. and C.C. K.K. was supported in part by a Swedish Research Council (Ventenskapsradet) International postdoc grant (2018- 00454).

## Author contributions

Conceptualization: CC; Study design: KK,CC; Organoid culture: KK, KrK, AS, AM, CK; Sequencing and library preparation: KK, KrK, AS, ZM; Histology: KK, KrK, WW, AM, YHL, CJS; Development of expressed cellular barcodes: KK, CC; Growth curve derivate estimation: EK; Analysis of sWGS and WGS data: KK, MJP, AK, HX, BL,CB, CC; Analysis of single cell RNA sequencing data: KK, MJP, CC; Analysis of ECB barcode data: KK, MJP, EK, KL, CC; Visualization: KK, MJP, HX, EK, AK, KEH, CC; Funding acquisition: CK, CC; Project administration: CC; Supervision: CK, CC; Writing – original draft: KK, MJP, CC. Writing – review and editing: all authors.

## Reagent availability

Requests for reagents should be directed to the corresponding author.

## Data availability

Metadata, cellranger outputs are available via Zenodo: https://doi.org/10.5281/zenodo.6401895 scRNA-seq data are available via bioProject ID PRJNA838456.

## Code availability

The computational methods, procedures and analyses summarized above are implemented in custom R, python and bash scripts will be available via the Curtis Lab Github: https://github.com/cancersysbio/gastric_organoid_evolution.

## Additional information

Supplementary Information is available for this paper.

Correspondence and requests for materials should be addressed to C.Curtis.

Reprints and permissions information is available at www.nature.com/reprints.

## Methods

A detailed description of the Materials and Methods is available in the Supplementary Information.

---

**Extended Data Fig. 1.**
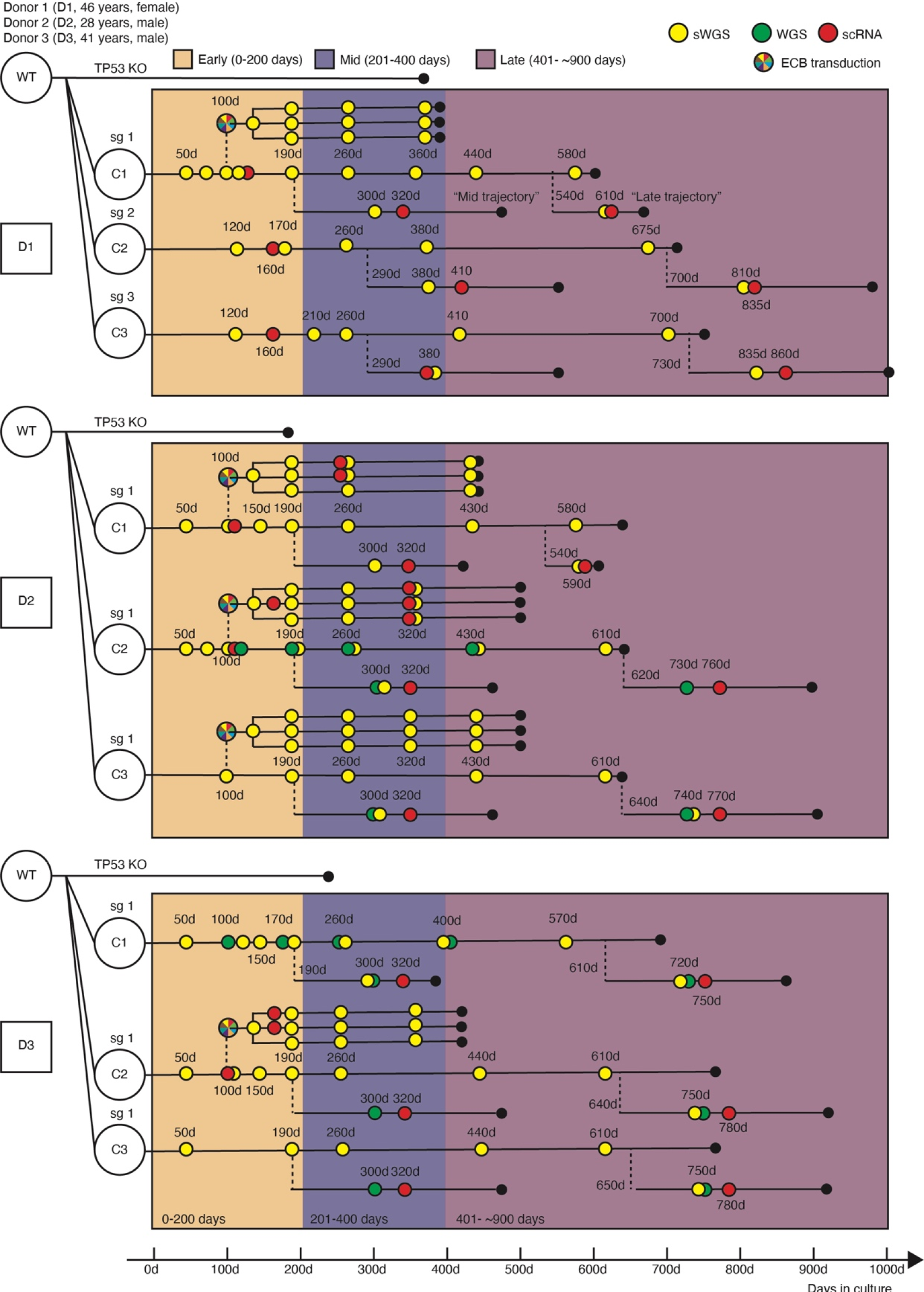
Schematic overview of gastric organoid (HGO) cultures, assays and sequencing time points. Organoids were established from three donors (abbreviated D) as wild-type (WT) cultures. For each donor (D1-D3), three independent CRISPR/Cas9 edited *TP53^-/-^* or *TP53^-/-^, APC^-/-^* cultures (abbreviated C) were established (indicated by sg 1-3) and referred to as C1-C3 (Methods). The WT and genome edited cultures were evolved under defined conditions for over two years. Sequencing was performed across the experimental time course at defined intervals: *Early* (∼0-200 days), *Mid* (∼200-400 days), *Late* (∼400-900 days). Each original culture was thawed (indicated by dashed lines) at an *Early/Mid* (190-290 days) and *Late* (540-730 days) time point for additional replication and comparisons. The thawed samples were treated with normocin to eliminate mycoplasma (Methods). All cultures were subject to shallow WGS (sWGS). A subset of cultures underwent deeper WGS and/or single cell RNA (scRNA)-sequencing at select time points. In addition to these non-barcoded cultures, representative *TP53^-/-^*HGO cultures from each donor were selected for prospective lineage tracing via transduction of a lentiviral expressed cellular barcode (ECB), as indicated by the multi-colored circle in the legend. These ECB cultures were similarly subject to sWGS and scRNA-seq. Broad time intervals are indicated as in the legend, while days in culture are provided for individual cultures. Note that scRNA for D1C3 “Mid trajectory” was sampled at day 413.

**Extended Data Fig. 2.**
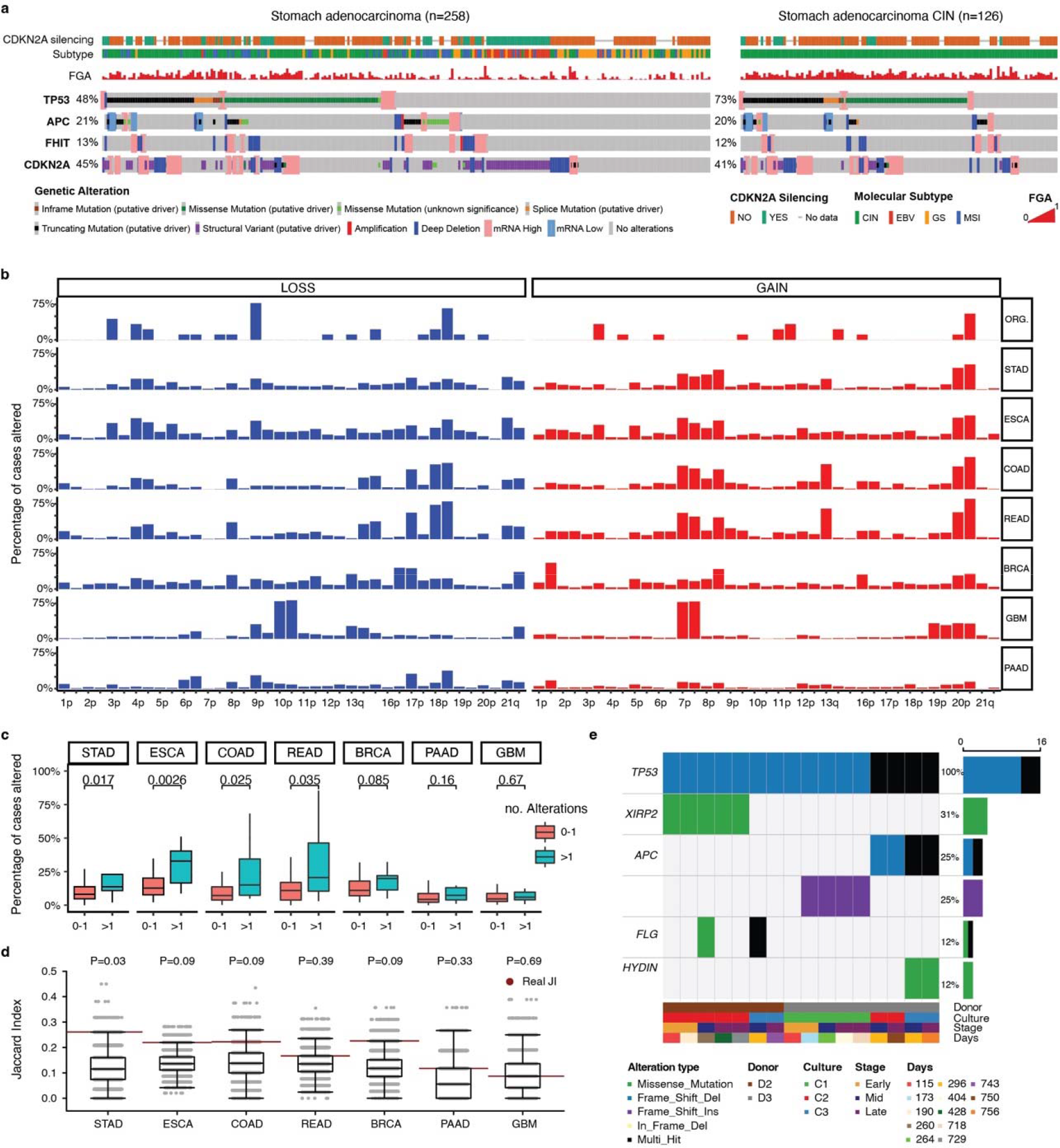
Recurrent copy number aberrations in *TP53* deficient gastric organoids (HGOs) are enriched in gastric and esophageal cancers. a, Prevalence of somatic alterations, including *TP53, APC, CDKN2A* and *FHIT* in gastric cancer (stomach adenocarcinoma, STAD) from TCGA and their association with molecular subgroups and fraction of genome altered. Data derive from the cBioPortal. b, Frequency of chromosome arm alterations in *TP53*^-/-^ HGOs at late time points (days 588 to 835, same time point as in Fig. 1c) relative to different tumor types profiled in TCGA. TCGA data were obtained from Firehose (http://gdac.broadinstitute.org/#). c, Enrichment of chromosome arm level alterations in *TP53*^-/-^ gastric organoid cultures across cancer types. Boxes show inter-quartile range (IQR), center lines represent the median, whiskers extend by 1.5 × IQR. Arm-level CNAs altered in two or more *TP53*^-/-^ gastric organoid cultures (n=11) were significantly more frequently altered than alterations present in 1 or fewer cultures (n=71) in both STAD and ESCA (p-value shown, two- sided Wilcoxon rank sum test). Tumor types assessed include Stomach (gastric) Adenocarcinoma (STAD), Esophageal carcinoma (ESCA), Colorectal Adenocarcinoma (COAD), Rectum Adenocarcinoma (READ), Breast invasive carcinoma (BRCA), Glioblastoma Multiforme (GBM), Pancreatic Adenocarcinoma (PAAD). d, The Jaccard Index (JI) was calculated by comparing CNAs that occurred in more than 1 chromosome arm in organoid cultures with CNAs occurring in 15% or more cases in a given tumor type. Then, for each tumor type, organoid CNA labels were randomly sampled (n=10,000) from all possible chromosome arm level events in a given TCGA tumor types, and the JI was calculated. The p-value was calculated as: P = 1- (sum(real JI > null Jis)/number of null Jis). e, Oncoplot showing alterations that occurred in two or more WGS samples for genes that are commonly (>10% of cases) mutated in esophageal adenocarcinoma or gastric cancer.

**Extended Data Fig. 3.**
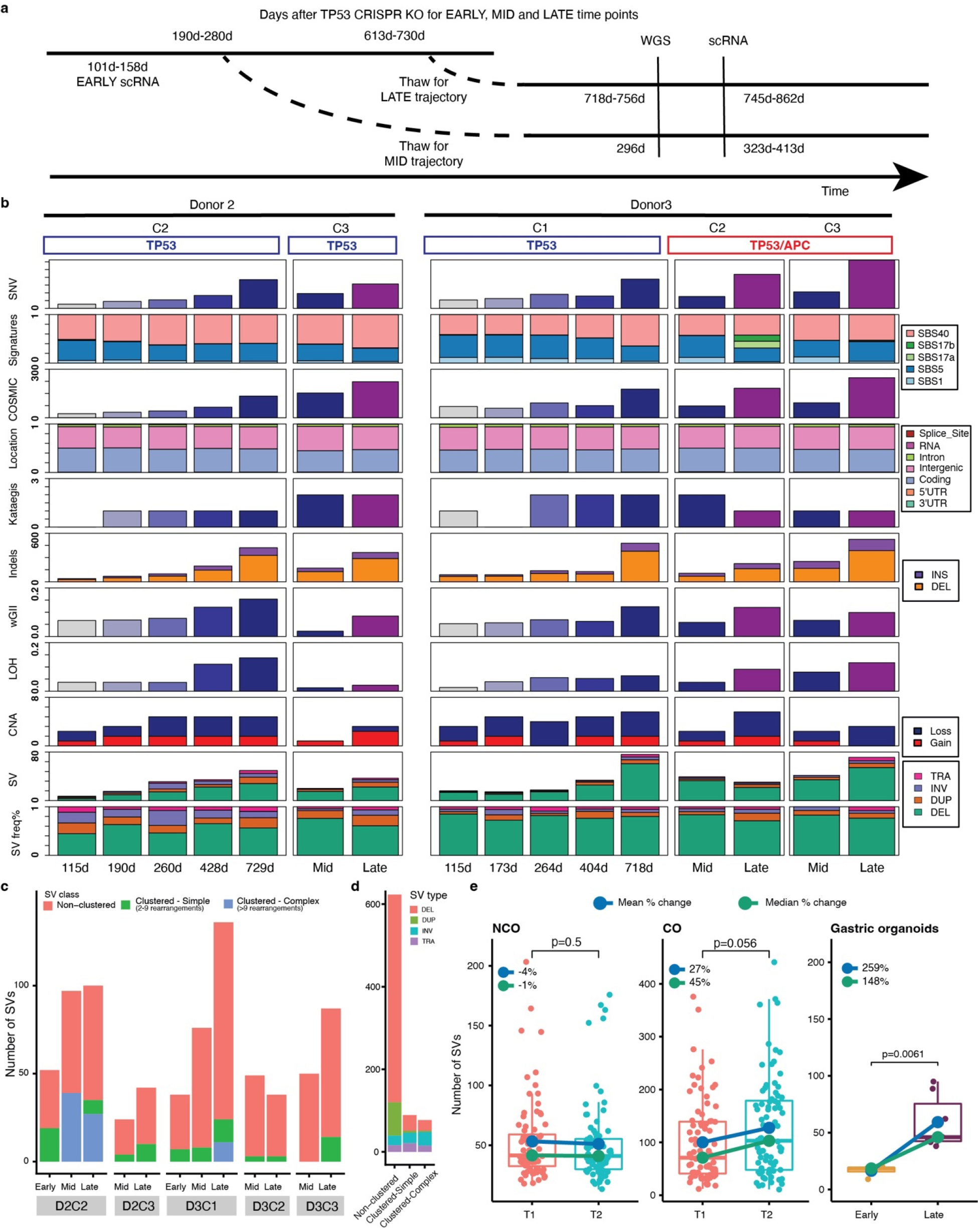
Longitudinal whole genome sequencing (WGS) of *TP53^-/-^* gastric organoids (HGOs). **a**, Overview of WGS and scRNA-seq time points for Early, Mid and Late cultures. Time is indicated in days (d). **b**, Summary of genomic features as assessed by WGS of multiple time points for Donors 2 and 3, expanding upon Fig. 2a. *Mid* and *Late* time points correspond to day 296 and 705-754, respectively. **c**, Distribution of non-clustered SVs, simple SVs (2-9 rearrangements) and complex SVs (10 or more rearrangements) defined using ClusterSV (Methods). **d**, Distribution of SV types across the three classes of SVs (non- clustered, clustered-simple and clustered-complex) **e**, Boxplot comparing total SV burden at the time of endoscopy (T1 and T2) for four Barrett’s esophagus biopsies per patient with cancer outcome (CO, n=160) or noncancer outcome (NCO; n=160) from the Paulson et al. cohort relative to the total SV burden between early (n=4) and late (n=9) timepoints in *TP53^-/-^*and *TP53^-/-^, APC^-/-^*HGOs (right). P-values were calculated using Wilcoxon rank sum test (two sided, unpaired). Boxes show IQR, center lines represent the median, whiskers extend by 1.5 × IQR.

**Extended Data Fig. 4.**
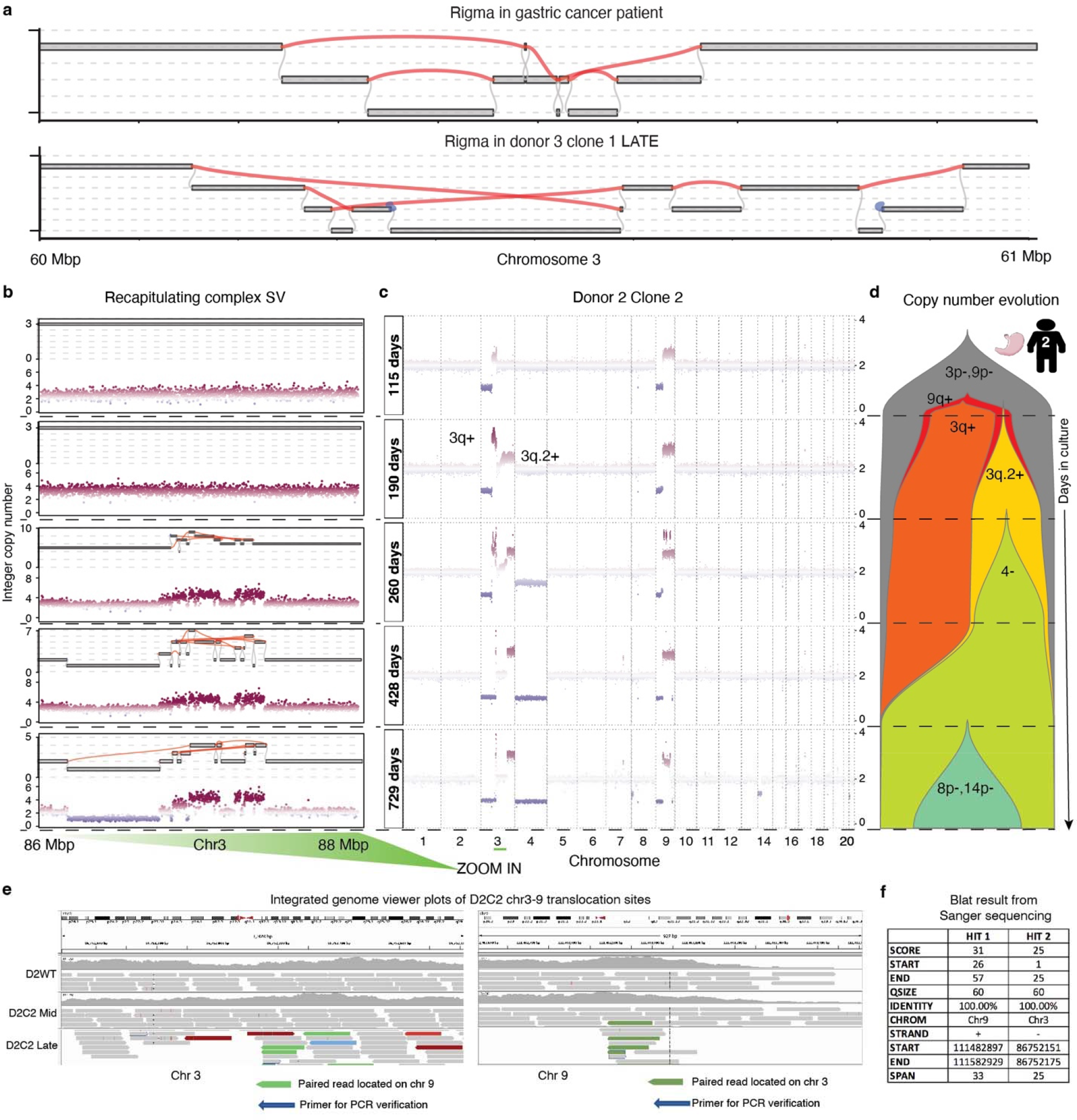
*TP53* deficient gastric organoids (HGOs) recapitulate complex structural variants (SVs) observed in gastric cancers. **a**, Complex rigma-like SVs seen in gastric cancer (GC) patients such as pfg008 from the Wang et al. cohort (Methods) are similar to those in *TP53^-/-^* HGOs including D3C1 (also shown in Fig. 2e). **b**, Zoomed-in view of a region on chromosome 3 which evolved complex SVs during *in vitro* culture. **c**, Corresponding CNA profiles based on longitudinal WGS of D2C2 at 5 timepoints spanning days 115 to 729 in culture. **d**, Fishplot for D2C2 depicting CNA evolution inferred from WGS. **e**, IGV plots indicate the translocation between chromosomes 3 and 9 for D2C2, corresponding to the complex SV in panel a. Primers used for PCR amplification, gel electrophoresis purification and Sanger sequencing as marked in the plots **f**, BLAT results from Sanger sequencing of the PCR primers demonstrating partial alignment to chr3 and chr9.

**Extended Data Fig. 5.**
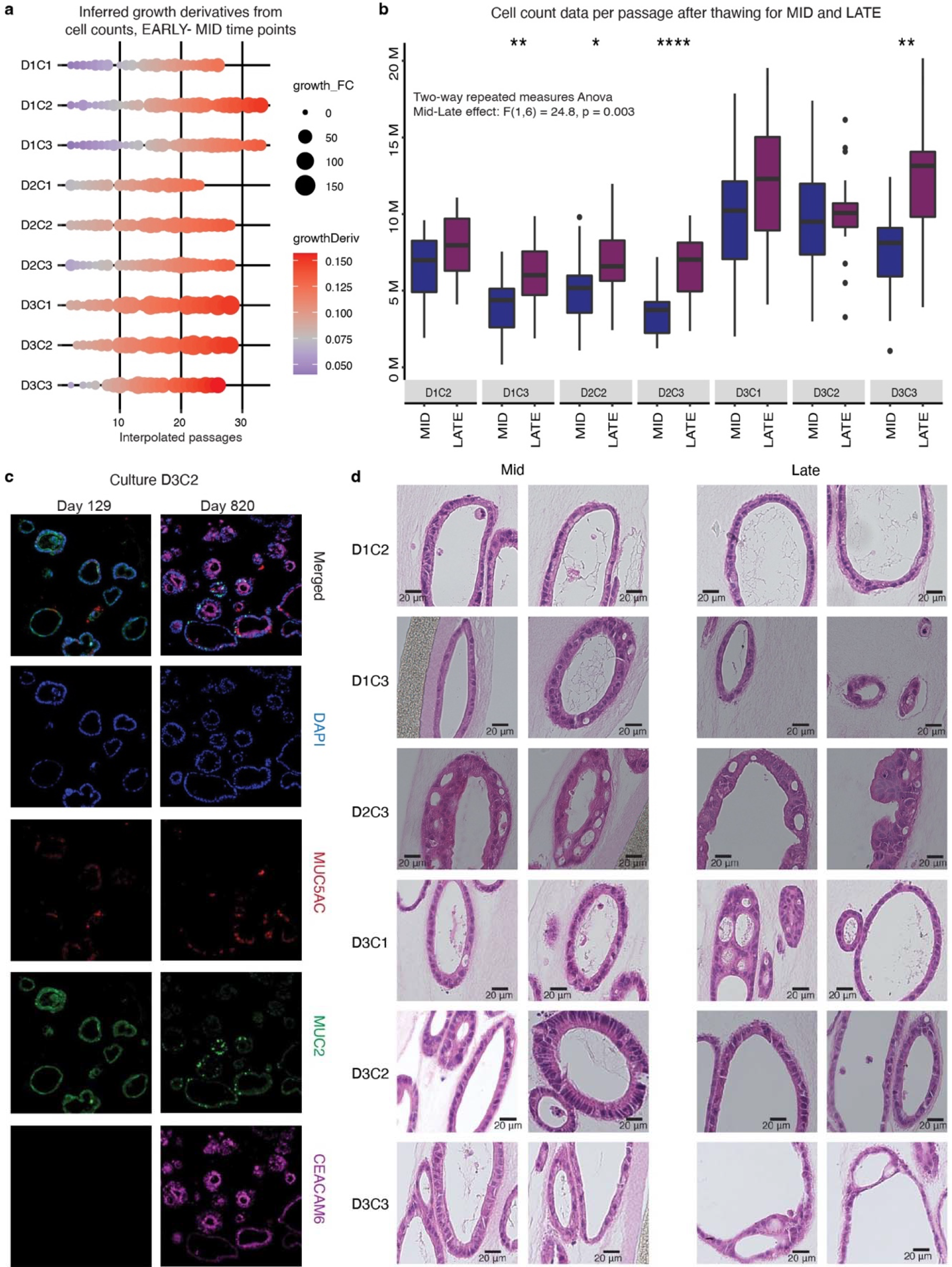
Prolonged culturing leads to increased growth rates and modest histologic changes. **a,** Dotplot of loess regression inferred growth derivative per culture over the Early-Mid trajectory, equivalent to that shown in Fig. 3b. **b,** Boxplots of raw cell numbers during passaging (y-axis) for seven cultures thawed at the Mid (∼320 days) and Late time points (∼770 days). Boxes show IQR, center lines represent the median, whiskers extend by 1.5 × IQR. Students t-test was used to compare time points for individual cultures (n=18 for each group, two-sided, not paired p-values are reported and normality assumed). To compare the Mid-Late effect across all cultures, two-way repeated measures ANOVA was used (Mid-Late effect and individual time points). The Mid-Late effect was significant (p-value 0.003). **c,** Representative immunofluorescent staining in culture D3C2 indicates that *TP53*-deficient, early passage organoids (passage 8) expressed *MUC2* but not *CEACAM6*. After prolonged *in vitro* evolution (passage 57), the HGOs expressed both *MUC2* and high level of *CEACAM6*. **d,** Hematoxylin and eosin (H&E) staining of *TP53* deficient HGOs at Mid and Late time points indicates the absence of mitoses and lack of apparent metaplasia/dysplasia per expert pathology review.

**Extended Data Fig. 6.**
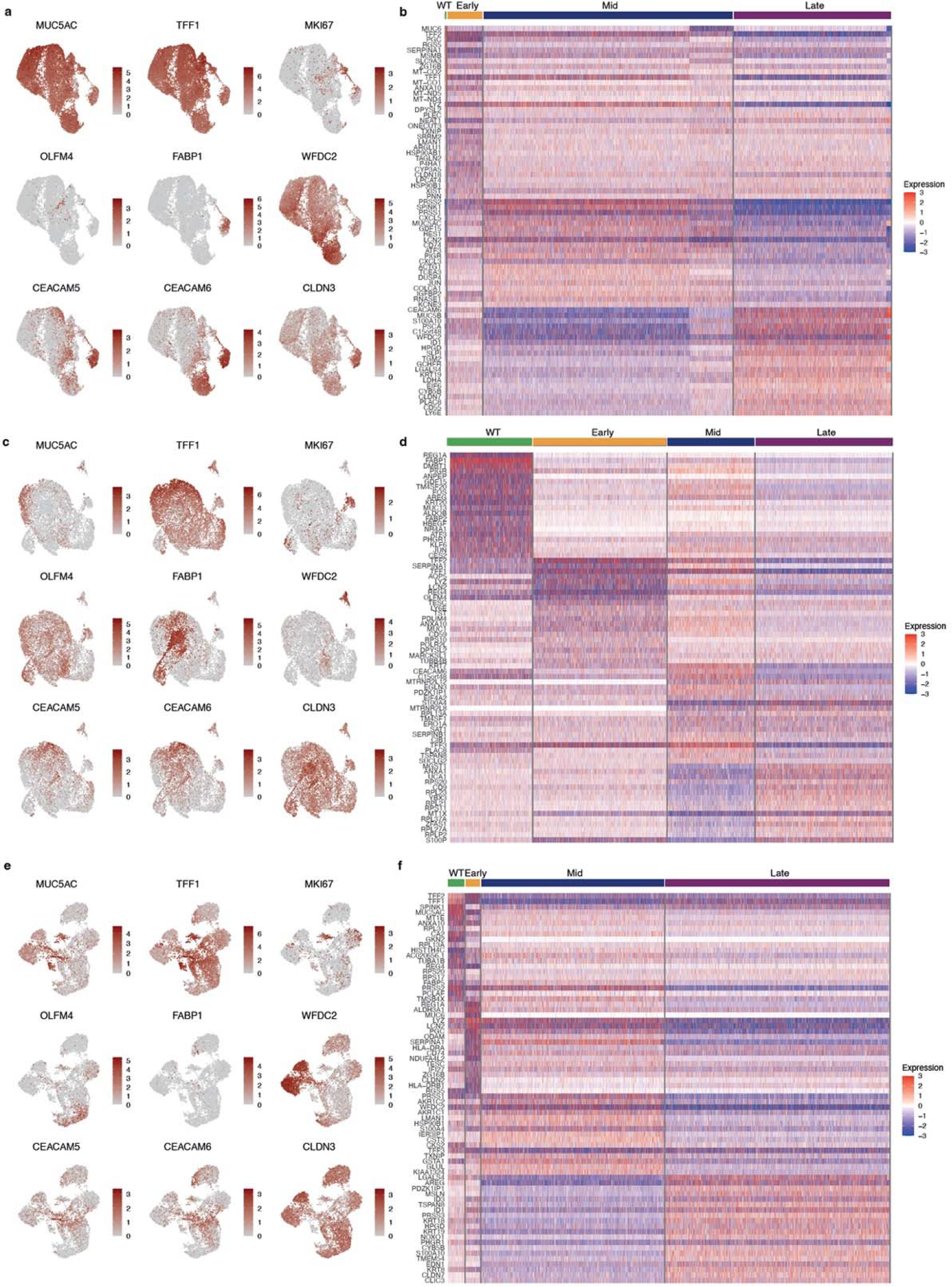
Differential gene expression highlights transcriptional heterogeneity across donors and time. **a,** UMAP embedding for D1 highlighting the expression of 9 literature-derived gastric marker genes. Each marker represents a specific cell identity: pit mucosal cells (PMCs; *MUC5AC*, *TFF1*), proliferating cells (PCs; *MKI67*), mucosal stem cells (MSCs; *OLFM4*), enterocytes (*FABP1*), goblet cells (*WFDC2*) and malignant cells (*CEACAM5*, *CEACAM6*, *CLDN3*). **b,** Heatmap showing the top 20 differentially expressed genes per timepoint (WT, Early, Mid, Late). **c-d,** Equivalent UMAP embedding and heatmap from panels a and b for D2. **E-f,** Equivalent UMAP embedding and heatmap as in panels a and b but for D3.

**Extended Data Fig. 7.**
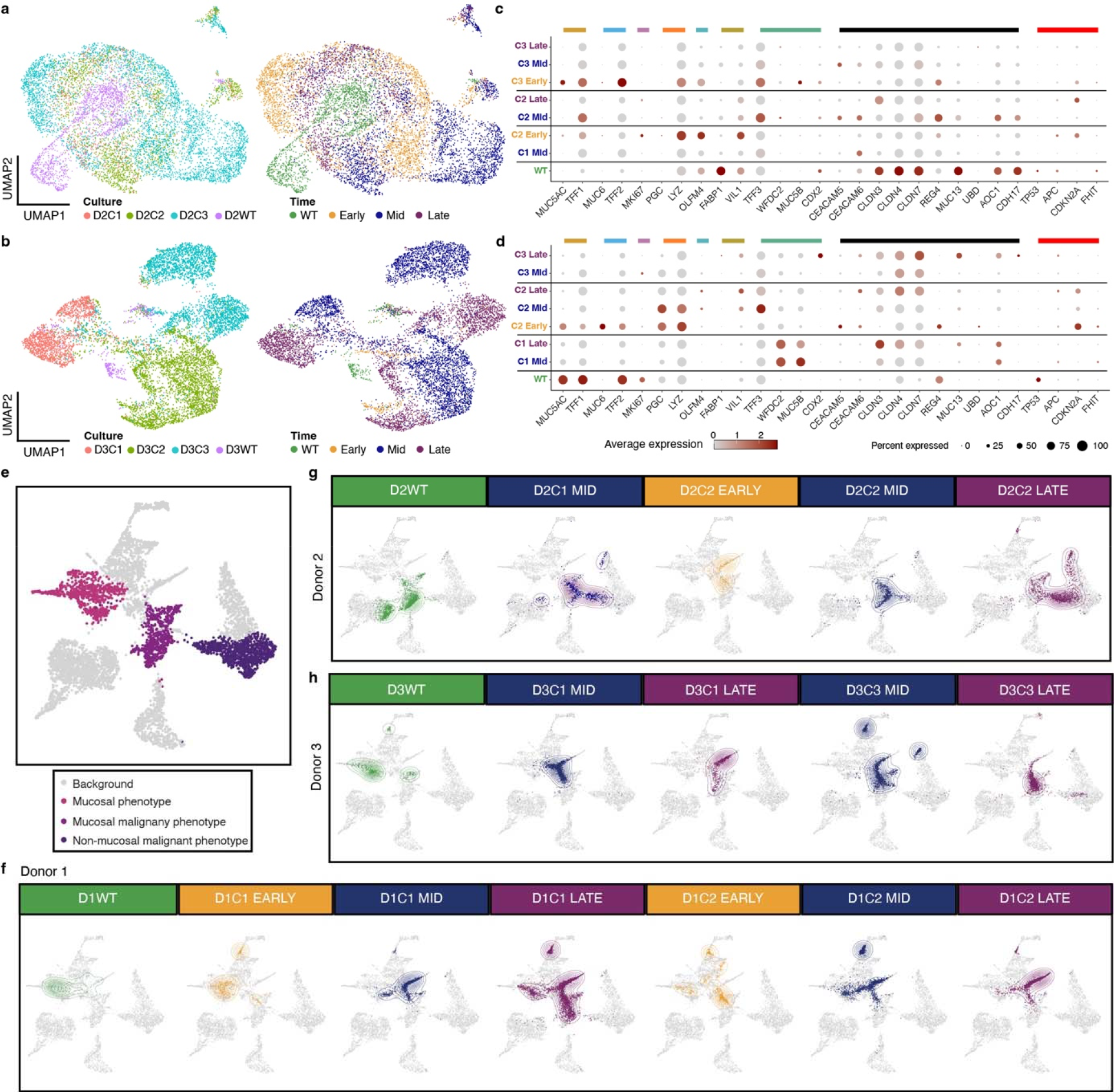
Latent Semantic Index (LSI) projection of gastric organoid (HGO) onto gastric tissue dataset. **a-b**, UMAP visualizations colored according to culture (left) and time point (right) for Donors 2 and 3 depicting 9,031 and 8,591 cells, respectively. **c-d**, Dotplot depicting the expression of selected marker genes for individual cultures and time points. Colored bars highlight marker genes associated with normal gastric and intestinal cell types, genes up-regulated in the gene expression profiling interactive analysis (GEPIA) of GC, and others of functional relevance. Pit mucosal cells: *MUC5AC*, *TFF1* – dark yellow; Gland mucosal cells: *MUC6*, *TFF2* – light blue; Proliferative cells: *MKI67* – purple; Neck-like cells: *PGC*, *LYZ* – orange; Mucosal stem cells: *OLFM4* – turquoise; Enterocytes: *FABP1*, *VIL1* – olive; Goblet cells: *TFF3*, *WFDC2*, *MUC5B*, *CDX2* – green; GEPIA top 12 genes: *CEACAM5*, *CEACAM6*, *CLDN3*, *CLDN4*, *CLDN7*, *REG4*, *MUC3A*, *MUC13*, *PI3*, *UBD*, *AOC1*, *CDH17* – black; Other: *TP53*, *APC*, *CDKN2A*, *FHIT* – red. **e,** The reference gastric tumor-normal dataset (Sathe et al.) for the LSI projection is shown with key cellular populations colored according to their expression phenotype, as denoted in the legend. This panel is identical to that shown in Fig. 4c. **f-h,** Showing the LSI projection of individual cultures for donors 1, 2 and 3.

**Extended Data Fig. 8.**
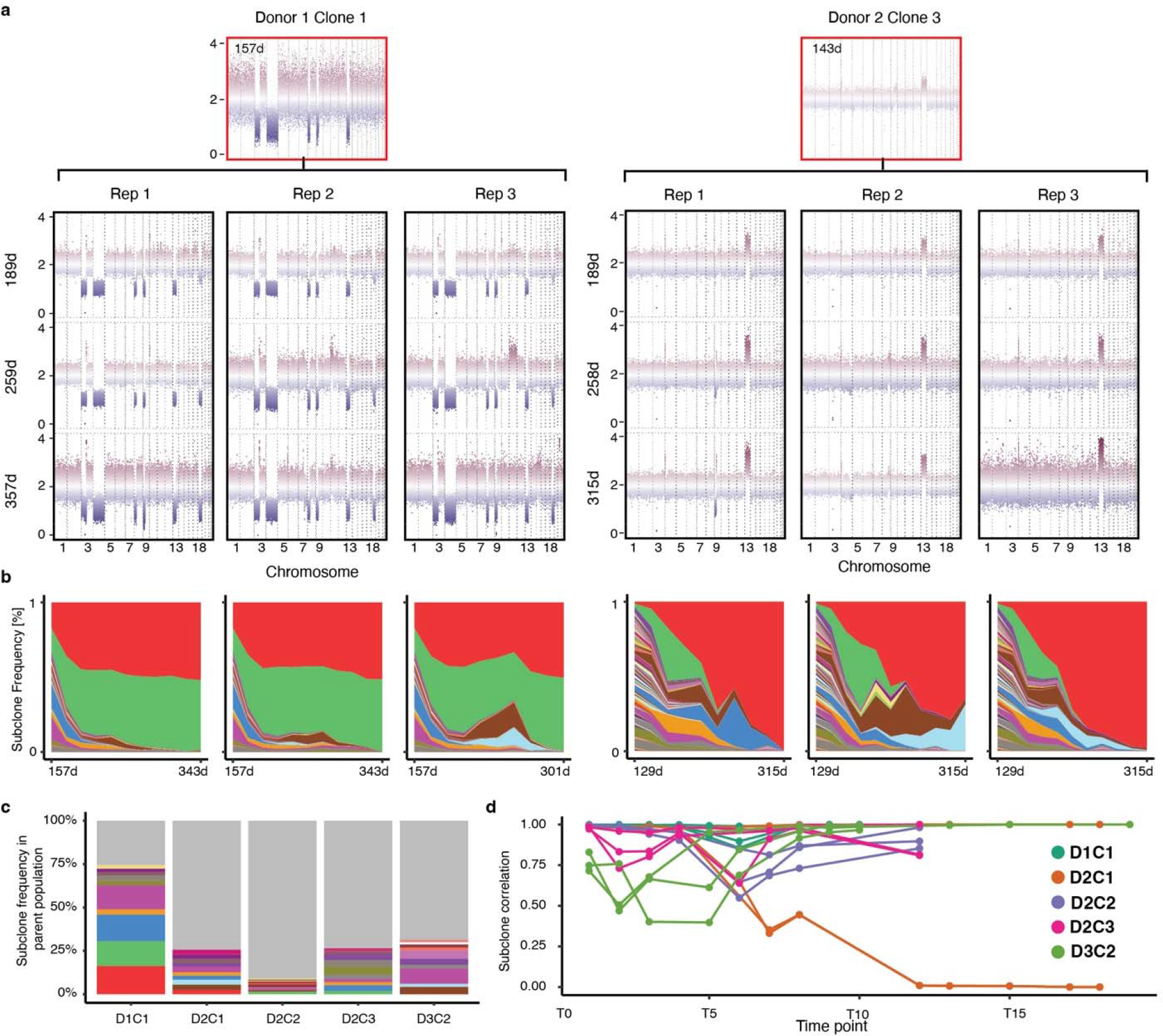
Subclone growth and frequency comparison and barcode trajectories for D1C1 and D2C3. **a**, sWGS of the parental population and three time points for D1C1 and D2C3, replicates 1-3. **b,** Mueller plots of the ECB subclone frequency over time for the cultures shown in panel a reveals similar lineage dynamics across replicates and deterministic outgrowth. **c**, Barplot depicting frequencies of the top 10 subclones in the parental population and the winning subclone (which was not one of the top subclones in the parental population). All other subclones colored gray. **d,** Pairwise Pearson correlation over time between replicate cultures for D1C1, D2C1, D2C2, D2C3 and D3C2. Time points correspond to passage times, where intervals between passages are approximately two weeks.

**Extended Data Fig. 9.**
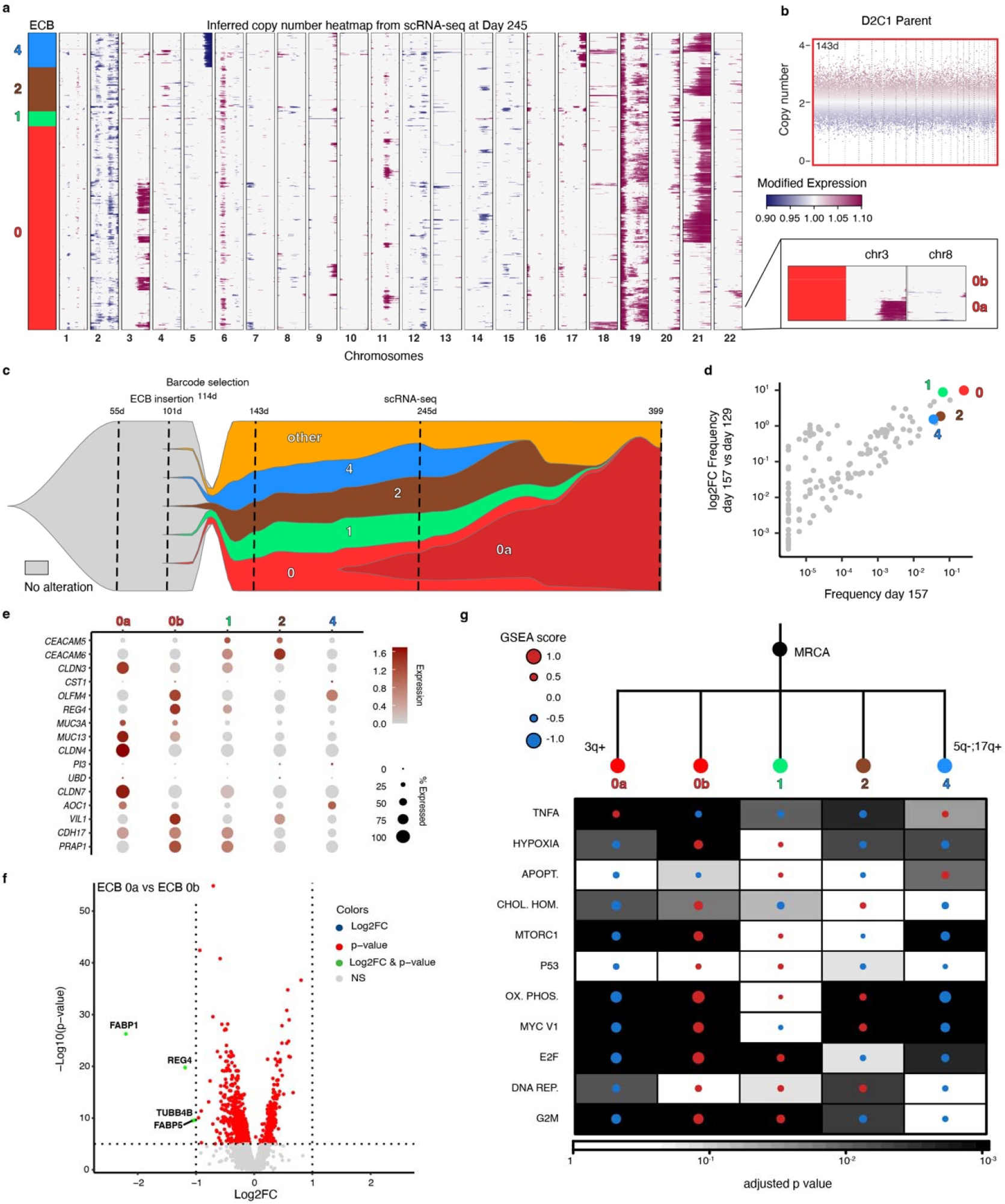
Linking single-cell genotypes to their transcriptional phenotypes in D2C1R1. **a,** Inferred copy number heatmap from scRNA-seq data at day 245, analogous to Fig. 6a. **b,** Copy number plot of parent population for D2C1. **c,** Fishplot schematic illustrating the link between lineage (ECBs) and copy number based subclones, similar to Fig. 6b. The actual subclone frequencies are shown in Fig. 5c. **d**, Scatterplot comparing subclone frequencies at day 157 and the fold change between days 129 and 157 for all subclones. Subclones of interest are highlighted as in panel a. **e**, Dotplot showing the expression of top differentially expressed genes (DEGs) based on GEPIA (gene expression profiling interactive analysis) of gastric cancer (GC). **f,** Volcano plot illustrating DEGs from the comparison of the winning subclone 0a and its parent subclone 0b. Vertical and horizontal lines correspond to absolute log2FC values of 1 and p < 0.00001 respectively. **g,** GSEA heatmap from MSigDB Hallmark gene sets showing the most significantly altered pathways for the highlighted subclones (right; Kolmogorov-Smirnov statistic, Benjamini-Hochberg adjusted). A manually reconstructed copy number phylogeny is shown above. This visualization is equivalent to Fig. 6f.

**Extended Data Fig. 10.**
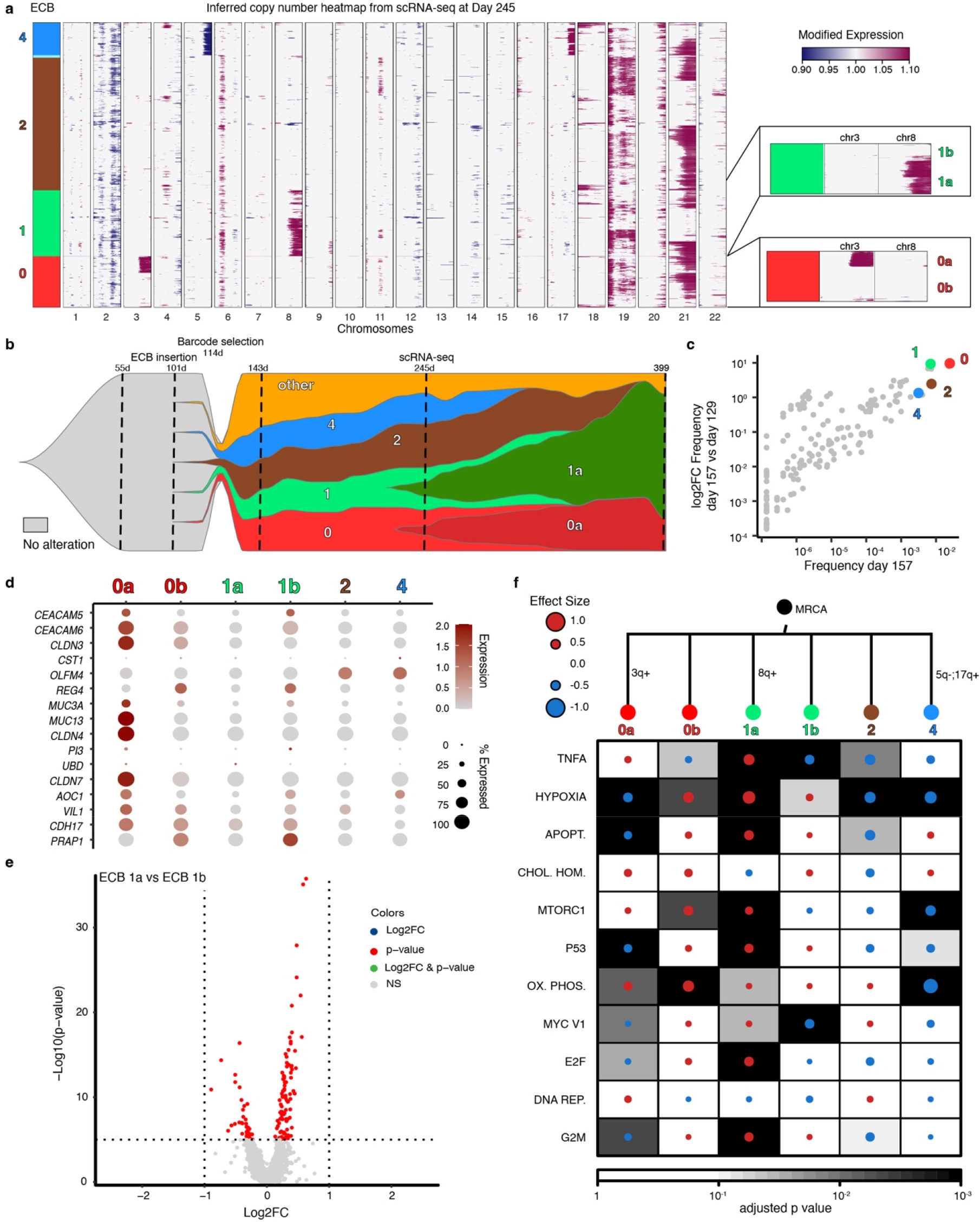
Linking single-cell genotypes to their transcriptional phenotypes in in D2C1R2. **a**, Inferred copy number heatmap from scRNA-seq data at day (d) 245, analogous to Fig. 6a. **b,** Fishplot schematic illustrating the link between lineage (ECBs) and copy number based subclones, similar to Fig. 6b. The actual subclone frequencies are shown in Fig. 5c. **c**, Scatterplot comparing subclone frequencies at day 157 and the fold change between days 129 and 157 for all subclones. Subclones of interest are highlighted as in panel a. **d,** Dotplot showing the expression of top differentially expressed genes (DEGs) based on GEPIA (gene expression profiling interactive analysis) of gastric cancer (GC). **e,** Volcano plot illustrating DEGs from the comparison of the winning subclone 0a and its parent subclone 0b. Vertical and horizontal lines correspond to absolute log2FC values of 1 and p < 0.00001 respectively. **f,** GSEA heatmap from MSigDB Hallmark gene sets showing the most significantly altered pathways for the highlighted subclones (right; Kolmogorov-Smirnov statistic, Benjamini- Hochberg adjusted). A manually reconstructed copy number phylogeny is shown above. This visualization is equivalent to Fig. 6f.

**Extended Data Fig. 11.**
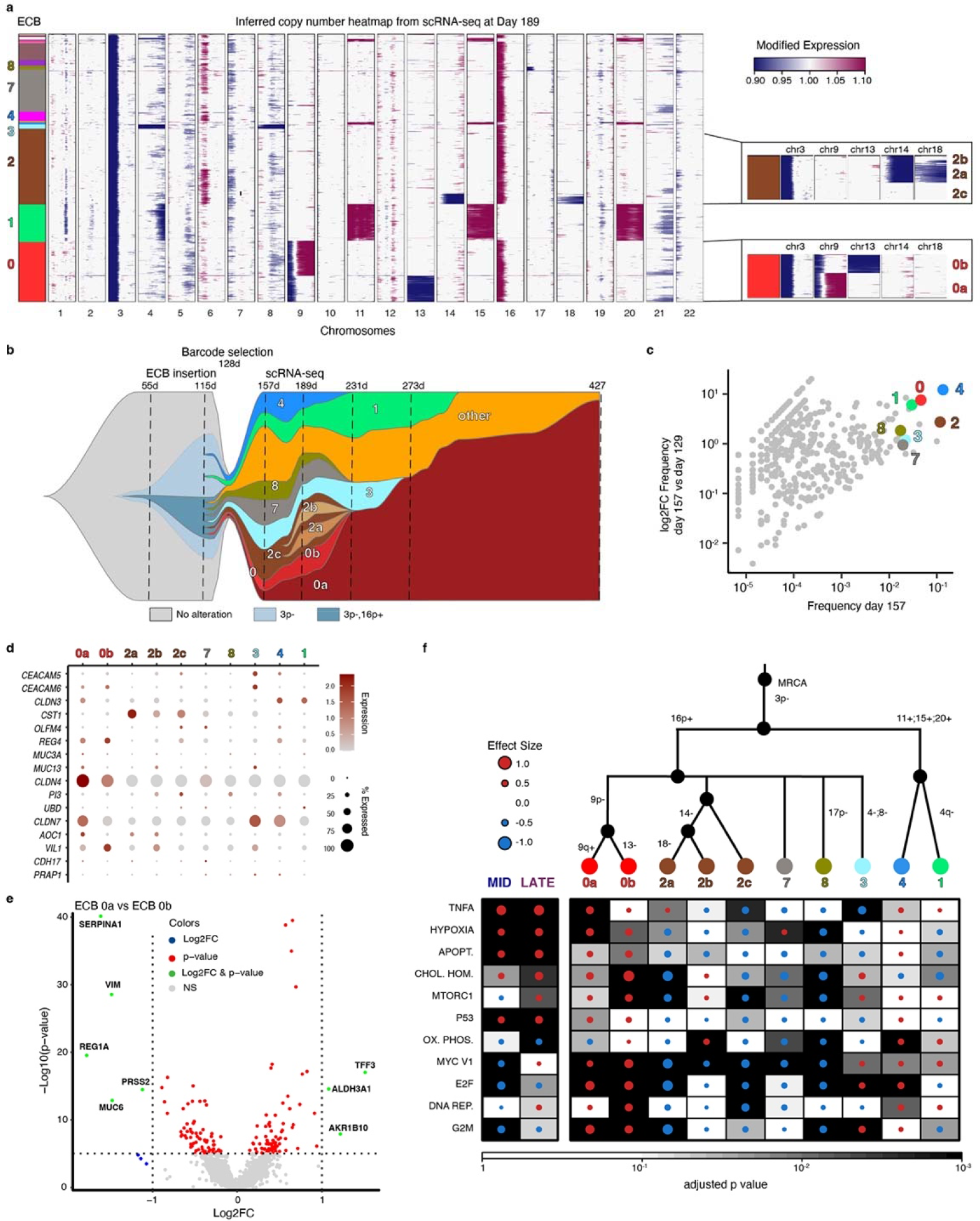
Linking single-cell genotypes to their transcriptional phenotypes in in D3C2R1. **a,** Inferred copy number heatmap from scRNA-seq data at day (d) 189, analogous to Fig. 6a. **b,** Fishplot schematic illustrating the link between lineage (ECBs) and copy number based subclones, similar to Fig. 6b. The actual subclone frequencies are shown in Fig. 5c. **c**, Scatterplot comparing subclone frequencies at day 157 and the fold change between days 129 and 157 for all subclones. Subclones of interest are highlighted as in panel A. **d,** Dotplot showing the expression of top differentially expressed genes (DEGs) based on GEPIA (gene expression profiling interactive analysis) of gastric cancer (GC). **e,** Volcano plot illustrating DEGs from the comparison of the winning subclone 0a and its parent subclone 0b. Vertical and horizontal lines correspond to absolute log2FC values of 1 and p < 0.00001 respectively. **f,** GSEA heatmap from MSigDB Hallmark gene sets showing the most significantly altered pathways for the highlighted subclones (right; Kolmogorov-Smirnov statistic, Benjamini- Hochberg adjusted). A manually reconstructed copy number phylogeny is shown above. The visualization is equivalent to Fig. 6f.

**Extended Data Fig. 12.**
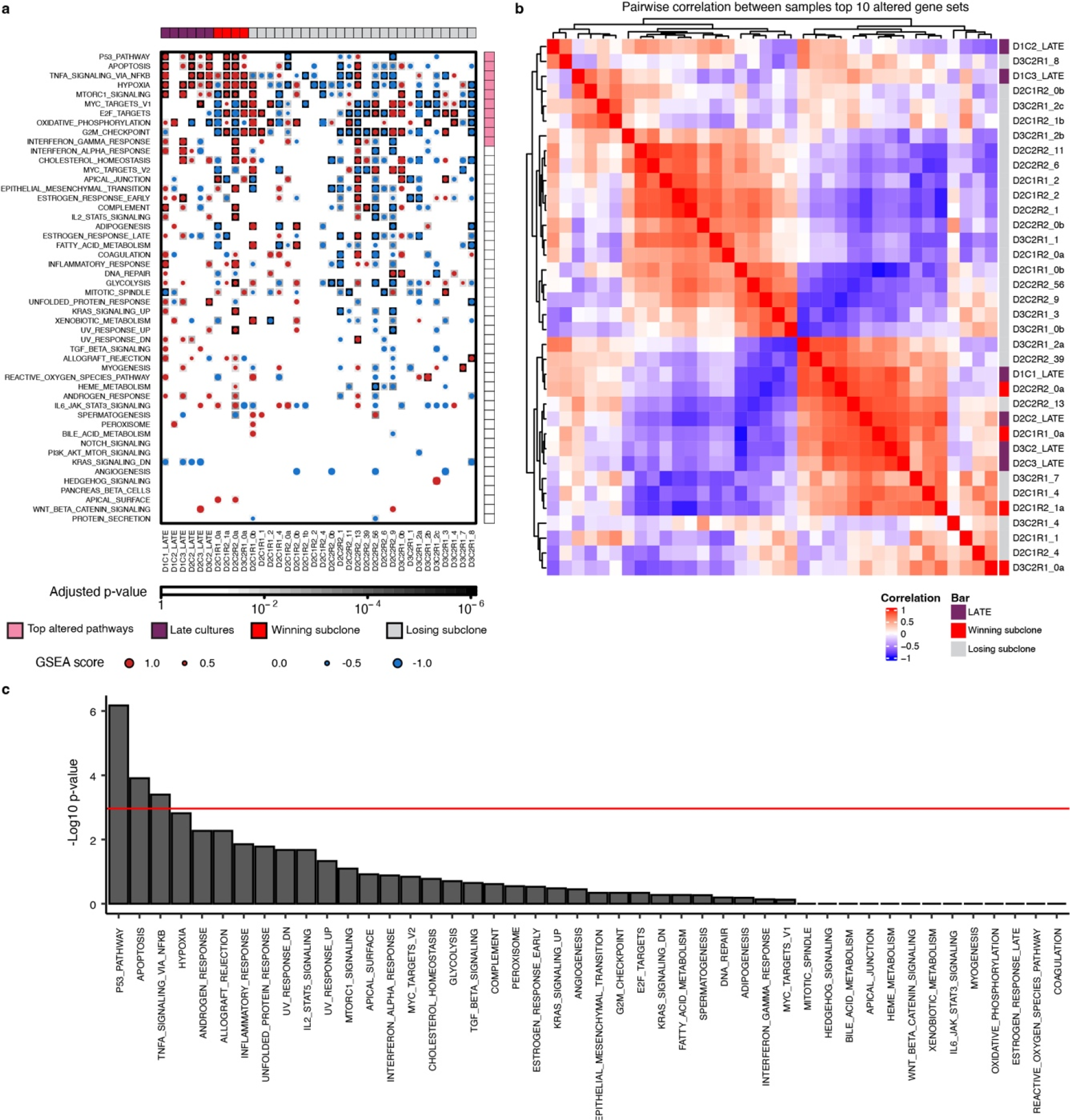
Enrichment of similar gene sets between winning subclones and *Late* cultures. **a,** GSEA heatmap from MSigDB Hallmark gene sets for each barcoded culture, including replicates: D2C2-R2 (day 173), D2C1-R1 (day 245), D2C1-R2 (day 245), D3C2-R1 (day 189), as well as non-barcoded *Late* cultures for D1C1, D1C2, D1C3, D2C2, D2C3 and D3C2. The GSEA score is indicated by dot size and colored according to the directionality of expression profiles (up, red; down, blue). Background shading indicates statistical significance (Kolmogorov-Smirnov statistic, Benjamini-Hochberg adjusted) **b,** Winning subclones and *Late* samples cluster together based on GSEA score. Pairwise spearman correlation between samples based on GSEA scores for the top 10 most altered pathways for all *Late* and winnings subclone samples. The most altered pathways are shown in a. This plot is identical to Fig. 6g but with sample annotations included. Note that the winning subclones (marked in red) cluster with the *Late* samples for D1C1, D2C2, D2C3 and D3C2 which exhibit a more malignant phenotype compared to *Late* samples D1C2 and D1C3 based on the LSI projection (Fig. 4c,d, Extended Data Fig. 7). **c,** Fisher exact test of independence (two-sided), comparing significance for each pathway, and status for each sample - *Late* and winning subclones (n=16) relative to all other subclones analyzed (n=29). The red line indicates the significance threshold (0.05) with Bonferroni correction.

## References

1. Crosby, D. et al. Early detection of cancer. Science 375, eaay9040 (2022).

2. Vázquez-García, I. et al. Clonal Heterogeneity Influences the Fate of New Adaptive Mutations. Cell Rep. 21, 732–744 (2017).

3. Lenski, R. E., Rose, M. R., Simpson, S. C. & Tadler, S. C. Long-Term Experimental Evolution in Escherichia coli. I. Adaptation and Divergence During 2,000 Generations. Am. Nat. 138, 1315–1341 (1991).

4. Good, B. H., McDonald, M. J., Barrick, J. E., Lenski, R. E. & Desai, M. M. The dynamics of molecular evolution over 60,000 generations. Nature 551, 45–50 (2017).

5. Sottoriva, A. et al. A Big Bang model of human colorectal tumor growth. Nat. Genet. 47, 209–216 (2015).

6. Sun, R. et al. Between-region genetic divergence reflects the mode and tempo of tumor evolution. Nat. Genet. 49, 1015–1024 (2017).

7. Laconi, E., Marongiu, F. & DeGregori, J. Cancer as a disease of old age: changing mutational and microenvironmental landscapes. Br. J. Cancer 122, 943–952 (2020).

8. Tsao, J. L. et al. Genetic reconstruction of individual colorectal tumor histories. Proc. Natl. Acad. Sci. U. S. A. 97, 1236–1241 (2000).

9. Baker, A.-M., Graham, T. A. & Wright, N. A. Pre-tumour clones, periodic selection and clonal interference in the origin and progression of gastrointestinal cancer: potential for biomarker development. J. Pathol. 229, 502–514 (2013).

10. Williams, M. J. et al. Quantification of subclonal selection in cancer from bulk sequencing data. Nat. Genet. 50, 895–903 (2018).

11. Rogers, Z. N. et al. Mapping the in vivo fitness landscape of lung adenocarcinoma tumor suppression in mice. Nat. Genet. 50, 483–486 (2018).

12. Gerstung, M. et al. The evolutionary history of 2,658 cancers. Nature 578, 122–128 (2020).

13. Waddingham, W. et al. Recent advances in the detection and management of early gastric cancer and its precursors. Frontline Gastroenterol. 12, 322–331 (2021).

14. Mysuru Shivanna, L. & Urooj, A. A Review on Dietary and Non-Dietary Risk Factors Associated with Gastrointestinal Cancer. J. Gastrointest. Cancer 47, 247–254 (2016).

15. Li, X. et al. Temporal and spatial evolution of somatic chromosomal alterations: a case- cohort study of Barrett’s esophagus. Cancer Prev. Res. 7, 114–127 (2014).

16. Paulson, T. G. et al. Somatic whole genome dynamics of precancer in Barrett’s esophagus reveals features associated with disease progression. Nat. Commun. 13, 2300 (2022).

17. Nowicki-Osuch, K. et al. Molecular phenotyping reveals the identity of Barrett’s esophagus and its malignant transition. Science 373, 760–767 (2021).

18. Yan, H. H. N. et al. A Comprehensive Human Gastric Cancer Organoid Biobank Captures Tumor Subtype Heterogeneity and Enables Therapeutic Screening. Cell Stem Cell 23, 882–897.e11 (2018).

19. Sethi, N. S. et al. Early TP53 alterations engage environmental exposures to promote gastric premalignancy in an integrative mouse model. Nat. Genet. 52, 219–230 (2020).

20. Seidlitz, T., Koo, B.-K. & Stange, D. E. Gastric organoids-an in vitro model system for the study of gastric development and road to personalized medicine. Cell Death Differ. 28, 68–83 (2021).

21. Lo, Y.-H. et al. A CRISPR/Cas9-Engineered ARID1A-Deficient Human Gastric Cancer Organoid Model Reveals Essential and Nonessential Modes of Oncogenic Transformation. Cancer Discov. 11, 1562–1581 (2021).

22. Cancer Genome Atlas Research Network. Comprehensive molecular characterization of gastric adenocarcinoma. Nature 513, 202–209 (2014).

23. Wang, K. et al. Whole-genome sequencing and comprehensive molecular profiling identify new driver mutations in gastric cancer. Nat. Genet. 46, 573–582 (2014).

24. Ben-David, U. & Amon, A. Context is everything: aneuploidy in cancer. Nat. Rev. Genet. 21, 44–62 (2020).

25. Narkar, A. et al. On the role of p53 in the cellular response to aneuploidy. Cell Rep. 34, 108892 (2021).

26. Weiss, M. B. et al. Deletion of p53 in human mammary epithelial cells causes chromosomal instability and altered therapeutic response. Oncogene 29, 4715–4724 (2010).

27. Drost, J. et al. Sequential cancer mutations in cultured human intestinal stem cells. Nature 521, 43–47 (2015).

28. Taylor, A. M. et al. Genomic and Functional Approaches to Understanding Cancer Aneuploidy. Cancer Cell 33, 676–689.e3 (2018).

29. Salehi, S. et al. Clonal fitness inferred from time-series modelling of single-cell cancer genomes. Nature 595, 585–590 (2021).

30. Barrett, M. T. et al. Evolution of neoplastic cell lineages in Barrett oesophagus. Nat. Genet. 22, 106–109 (1999).

31. Saldivar, J. C. & Park, D. Mechanisms shaping the mutational landscape of the FRA3B/FHIT-deficient cancer genome. Genes Chromosomes Cancer 58, 317–323 (2019).

32. Newell, F. et al. Complex structural rearrangements are present in high-grade dysplastic Barrett’s oesophagus samples. BMC Med. Genomics 12, 31 (2019).

33. Alexandrov, L. B. et al. Signatures of mutational processes in human cancer. Nature 500, 415–421 (2013).

34. Hadi, K. et al. Distinct Classes of Complex Structural Variation Uncovered across Thousands of Cancer Genome Graphs. Cell 183, 197–210.e32 (2020).

35. Glover, T. W., Wilson, T. E. & Arlt, M. F. Fragile sites in cancer: more than meets the eye. Nat. Rev. Cancer 17, 489–501 (2017).

36. Birchenough, G. M. H., EV Johansson, M., Gustafsson, J. K., Bergström, J. H. & Hansson, G. C. New developments in goblet cell mucus secretion and function. Mucosal Immunology vol. 8 712–719 Preprint at https://doi.org/10.1038/mi.2015.32 (2015).

37. Dong, D., Mu, Z., Zhao, C. & Sun, M. ZFAS1: a novel tumor-related long non-coding RNA. Cancer Cell Int. 18, 125 (2018).

38. Rao, S. et al. β2-spectrin (SPTBN1) as a therapeutic target for diet-induced liver disease and preventing cancer development. Sci. Transl. Med. 13, eabk2267 (2021).

39. Paludan, S. R., Reinert, L. S. & Hornung, V. DNA-stimulated cell death: implications for host defence, inflammatory diseases and cancer. Nat. Rev. Immunol. 19, 141–153 (2019).

40. Zhang, M. et al. Dissecting transcriptional heterogeneity in primary gastric adenocarcinoma by single cell RNA sequencing. Gut 70, 464–475 (2021).

41. Sathe, A. et al. Single-Cell Genomic Characterization Reveals the Cellular Reprogramming of the Gastric Tumor Microenvironment. Clin. Cancer Res. 26, 2640–2653 (2020).

42. Zhang, P. et al. Dissecting the Single-Cell Transcriptome Network Underlying Gastric Premalignant Lesions and Early Gastric Cancer. Cell Rep. 30, 4317 (2020).

43. Lang, G. I. et al. Pervasive genetic hitchhiking and clonal interference in forty evolving yeast populations. Nature 500, 571–574 (2013).

44. Levy, S. F. et al. Quantitative evolutionary dynamics using high-resolution lineage tracking. Nature 519, 181–186 (2015).

45. Nguyen Ba, A. N., et al. High-resolution lineage tracking reveals travelling wave of adaptation in laboratory yeast. Nature 575, 494–499 (2019).

46. Fujimoto, K. et al. Regulation of intestinal homeostasis by the ulcerative colitis-associated gene RNF186. Mucosal Immunol. 10, 446–459 (2017).

47. Takeno, A. et al. Gene expression profile prospectively predicts peritoneal relapse after curative surgery of gastric cancer. Ann. Surg. Oncol. 17, 1033–1042 (2010).

48. Maity, A. K. et al. Novel epigenetic network biomarkers for early detection of esophageal cancer. Clin. Epigenetics 14, 23 (2022).

49. You, X. et al. Galectin-1 promotes vasculogenic mimicry in gastric adenocarcinoma via the Hedgehog/GLI signaling pathway. Aging 12, 21837–21853 (2020).

50. Fearon, E. R. & Vogelstein, B. A genetic model for colorectal tumorigenesis. Cell 61, 759–767 (1990).

51. Baslan, T. et al. Ordered and deterministic cancer genome evolution after p53 loss. Nature (2022) doi:10.1038/s41586-022-05082-5.

52. Killcoyne, S. et al. Genomic copy number predicts esophageal cancer years before transformation. Nat. Med. 26, 1726–1732 (2020).

53. Cross, W. et al. The evolutionary landscape of colorectal tumorigenesis. Nat Ecol Evol 2, 1661–1672 (2018).

## References

54. Sato, T. et al. Long-term expansion of epithelial organoids from human colon, adenoma, adenocarcinoma, and Barrett’s epithelium. Gastroenterology 141, 1762–1772 (2011).

55. Bartfeld, S. et al. In vitro expansion of human gastric epithelial stem cells and their responses to bacterial infection. Gastroenterology 148, 126–136.e6 (2015).

56. Schwank, G. et al. Functional repair of CFTR by CRISPR/Cas9 in intestinal stem cell organoids of cystic fibrosis patients. Cell Stem Cell 13, 653–658 (2013).

57. Roerink, S. F. et al. Intra-tumour diversification in colorectal cancer at the single-cell level. Nature 556, 457–462 (2018).

58. Brinkman, E. K., Chen, T., Amendola, M. & van Steensel, B. Easy quantitative assessment of genome editing by sequence trace decomposition. Nucleic Acids Res. 42, e168 (2014).

59. Reber, S. et al. CRISPR-Trap: a clean approach for the generation of gene knockouts and gene replacements in human cells. Mol. Biol. Cell 29, 75–83 (2018).

60. Neal, J. T. et al. Organoid Modeling of the Tumor Immune Microenvironment. Cell 175, 1972–1988.e16 (2018).

61. Bhang, H.-E. C. et al. Studying clonal dynamics in response to cancer therapy using high- complexity barcoding. Nat. Med. 21, 440–448 (2015).

62. Stoeckius, M. et al. Cell Hashing with barcoded antibodies enables multiplexing and doublet detection for single cell genomics. Genome Biol. 19, 224 (2018).

63. Pleguezuelos-Manzano, C. et al. Establishment and culture of human intestinal organoids derived from adult stem cells. Curr. Protoc. Immunol. 130, e106 (2020).

64. Olarerin-George, A. O. & Hogenesch, J. B. Assessing the prevalence of mycoplasma contamination in cell culture via a survey of NCBI’s RNA-seq archive. Nucleic Acids Res. 43, 2535–2542 (2015).

65. Uemura, N. et al. Helicobacter pylori infection and the development of gastric cancer. N. Engl. J. Med. 345, 784–789 (2001).

66. Wang, J., Wang, Y., Li, Z., Gao, X. & Huang, D. Global Analysis of Microbiota Signatures in Four Major Types of Gastrointestinal Cancer. Front. Oncol. 11, 685641 (2021).

67. Nascimento Araujo, C. do et al. Evaluating the presence of Mycoplasma hyorhinis, Fusobacterium nucleatum, and Helicobacter pylori in biopsies of patients with gastric cancer. Infect. Agent. Cancer 16, 70 (2021).

68. Zella, D. et al. Mycoplasma promotes malignant transformation in vivo, and its DnaK, a bacterial chaperone protein, has broad oncogenic properties. Proc. Natl. Acad. Sci. U. S. A. 115, E12005–E12014 (2018).

69. Sethi, N., et al. Mutant p53 induces a hypoxia transcriptional program in gastric and esophageal adenocarcinoma. JCI Insight 4, (2019).

70. Gabridge, M. G. & Lundin, D. J. Cell Culture User’s Guide to Mycoplasma Detection and Control. Bionique Testing Laboratories, Saranac Lake, NY.

71. Smart, D. J. & Lynch, A. M. Evaluating the genotoxicity of topoisomerase-targeted antibiotics. Mutagenesis 27, 359–365 (2012).

72. Bhattacharya, P., Mukherjee, S. & Mandal, S. M. Fluoroquinolone antibiotics show genotoxic effect through DNA-binding and oxidative damage. Spectrochim. Acta A Mol. Biomol. Spectrosc. 227, 117634 (2020).

73. Garcia, M. et al. Sarek: A portable workflow for whole-genome sequencing analysis of germline and somatic variants. F1000Res. 9, 63 (2020).

74. Li, H. & Durbin, R. Fast and accurate short read alignment with Burrows-Wheeler transform. Bioinformatics 25, 1754–1760 (2009).

75. McKenna, A. et al. The Genome Analysis Toolkit: a MapReduce framework for analyzing next-generation DNA sequencing data. Genome Res. 20, 1297–1303 (2010).

76. Scheinin, I. et al. DNA copy number analysis of fresh and formalin-fixed specimens by shallow whole-genome sequencing with identification and exclusion of problematic regions in the genome assembly. Genome Res. 24, 2022–2032 (2014).

77. Kim, S. et al. Strelka2: fast and accurate calling of germline and somatic variants. Nat. Methods 15, 591–594 (2018).

78. Cameron, D. L. et al. GRIDSS, PURPLE, LINX: Unscrambling the tumor genome via integrated analysis of structural variation and copy number. bioRxiv 781013 (2019) doi:10.1101/781013.

79. Priestley, P. et al. Pan-cancer whole-genome analyses of metastatic solid tumours. Nature 575, 210–216 (2019).

80. Endesfelder, D. et al. Chromosomal instability selects gene copy-number variants encoding core regulators of proliferation in ER+ breast cancer. Cancer Res. 74, 4853–4863 (2014).

81. Ha, G. et al. TITAN: inference of copy number architectures in clonal cell populations from tumor whole-genome sequence data. Genome Res. 24, 1881–1893 (2014).

82. Shen, R. & Seshan, V. E. FACETS: allele-specific copy number and clonal heterogeneity analysis tool for high-throughput DNA sequencing. Nucleic Acids Res. 44, e131 (2016).

83. Zaccaria, S. & Raphael, B. J. Accurate quantification of copy-number aberrations and whole-genome duplications in multi-sample tumor sequencing data. Nat. Commun. 11, 1–13 (2020).

84. Das, S. et al. Next-generation genotype imputation service and methods. Nat. Genet. 48, 1284–1287 (2016).

85. Chen, X. et al. Manta: rapid detection of structural variants and indels for germline and cancer sequencing applications. Bioinformatics 32, 1220–1222 (2015).

86. Rausch, T. et al. DELLY: structural variant discovery by integrated paired-end and split- read analysis. Bioinformatics 28, i333–i339 (2012).

87. Wala, J. A. et al. SvABA: genome-wide detection of structural variants and indels by local assembly. Genome Res. 28, 581–591 (2018).

88. Li, Y. et al. Patterns of somatic structural variation in human cancer genomes. Nature 578, 112–121 (2020).

89. Obenchain, V. et al. VariantAnnotation : a Bioconductor package for exploration and annotation of genetic variants. Bioinformatics 30, 2076–2078 (2014).

90. Tate, J. G. et al. COSMIC: the Catalogue Of Somatic Mutations In Cancer. Nucleic Acids Res. 47, D941–D947 (2019).

91. Alexandrov, L. B. et al. The repertoire of mutational signatures in human cancer. Nature 578, 94–101 (2020).

92. Gori, K. & Baez-Ortega, A. sigfit: flexible Bayesian inference of mutational signatures. bioRxiv 372896 (2020) doi:10.1101/372896.

93. Miller, C. A. et al. Visualizing tumor evolution with the fishplot package for R. BMC Genomics 17, 880 (2016).

94. Smith, T., Heger, A. & Sudbery, I. UMI-tools: modeling sequencing errors in Unique Molecular Identifiers to improve quantification accuracy. Genome Res. 27, 491–499 (2017).

95. Tirosh, I. et al. Single-cell RNA-seq supports a developmental hierarchy in human oligodendroglioma. Nature 539, 309–313 (2016).

96. Hafemeister, C. & Satija, R. Normalization and variance stabilization of single-cell RNA-seq data using regularized negative binomial regression. Genome Biol. 20, 296 (2019).

97. McGinnis, C. S., Murrow, L. M. & Gartner, Z. J. DoubletFinder: Doublet Detection in Single- Cell RNA Sequencing Data Using Artificial Nearest Neighbors. Cell Systems 8, 329–337.e4 (2019).

98. Tang, Z. et al. GEPIA: a web server for cancer and normal gene expression profiling and interactive analyses. Nucleic Acids Res. 45, W98–W102 (2017).

99. Granja, J. M. et al. Single-cell multiomic analysis identifies regulatory programs in mixed- phenotype acute leukemia. Nat. Biotechnol. 37, 1458–1465 (2019).

100. Andreatta, M. et al. Interpretation of T cell states from single-cell transcriptomics data using reference atlases. Nat. Commun. 12, 2965 (2021).

101. Stuart, T. et al. Comprehensive Integration of Single-Cell Data. Cell (2019) doi:10.1016/j.cell.2019.05.031.

102. Tickle, T., Tirosh, I., Georgescu, C., Brown, M. & Haas, B. inferCNV of the Trinity CTAT Project. Klarman Cell Observatory, Broad Institute of MIT and Harvard (2019).

103. Patel, A. P. et al. Single-cell RNA-seq highlights intratumoral heterogeneity in primary glioblastoma. Science vol. 344 1396–1401 doi.org/10.1126/science.1254257 (2014).

104. Fan, J. et al. Linking transcriptional and genetic tumor heterogeneity through allele analysis of single-cell RNA-seq data. Genome Res. 28, 1217–1227 (2018).

105. Liberzon, A. et al. The Molecular Signatures Database (MSigDB) hallmark gene set collection. Cell Syst 1, 417–425 (2015).

