## Supplementary Materials and Methods for "Deterministic evolution and stringent selection during pre-neoplasia"

[1]Department of Medicine, Division of Oncology, Stanford University School of Medicine, Stanford, CA, USA. [2]Department of Genetics, Stanford University School of Medicine, Stanford, CA, USA. [3]Stanford Cancer Institute, Stanford University School of Medicine, Stanford, CA, USA. [4]Science for Life Laboratory and Department of Oncology-Pathology, Karolinska Institutet, Stockholm, Sweden. [5]Department of Biology, Stanford University School of Medicine, Stanford, CA, USA. [6]Department of Medicine, Division of Hematology, Stanford University School of Medicine, Stanford, CA USA. [7]Department of Cell & Developmental Biology, University College London, London, UK. [8]Department of Pathology, Stanford University School of Medicine, Stanford, CA, USA. [9]Department of Biomedical Data Science, Stanford University School of Medicine, Stanford, CA, USA. [10]Chan Zuckerberg Biohub

[%]Current address: Wellcome Sanger Institute & University of Cambridge, Hinxton Cambridgeshire, UK
[†]Contributed equally

### Materials and Methods

#### Generation of genome-edited human gastric organoids

##### Establishment of human gastric organoid cultures

Clinical samples (gastric corpus tissue) were obtained with informed consent from three patients undergoing sleeve gastrectomy under an IRB approved protocol (# 11977) through the Stanford University Hospital Tissue Procurement Shared Resource facility. Specimens were confirmed to be non-malignant and were used to generate wild-type (WT) human gastric organoid (HGO) cultures and subsequently CRISPR/Cas9 edited cultures based on an established protocol [54]. HGOs were generated as previously described with minor adaptations [21,55]. Briefly, tissue was placed in 10 ml PBS plus PSQ, dissected into smaller pieces and placed in ice cold chelation buffer containing 5.6 mmol/L $Na_2HPO_4$, 8.0 mmol/L $KH_2PO_4$, 96.2 mmol/L NaCl, 1.6 mmol/L KCl, 43.4 mmol/L sucrose, 54.9 mmol/L D-sorbitol and 0.5 mmol/L DTT. Next, 10 mmol/L EDTA pH 8.0 was added to the sample which was placed in a 4ºC rocking chamber for 3-5 hours. After incubation, the chelation buffer plus EDTA was removed and the sample was resuspended in 5 ml chelation buffer alone. Samples were vigorously shaken by hand (100 times) and supernatant containing stomach crypts were stored in a 15 ml tube. This step was repeated 8 times and each fraction was examined under a microscope. The fractions with the highest number of clean stomach crypts were combined and plated in Matrigel (R&D systems, Basement Membrane Extract type 2) in a 24 well plate. After the Matrigel had solidified, WENR media was added with the addition of 10 µM Y-27632 (Peprotech, #1293823) and 3 µM CHIR-99021 (R&D Systems, #4223) compounds as well as 200 ng/mL fibroblast growth factor (FGF) (Peprotech, #100-26).

##### Organoid culturing

The organoid culture media contained Advanced DMEM/F-12 (Thermo Fisher Scientific, #12634028) with 1X Penicillin/Streptomycin/Glutamine (Thermo Fisher Scientific, #10378016), 5% FBS, 1 mM HEPES (Thermo Fisher Scientific, #15630080), 1 mM N-Acetylcysteine (Sigma, A9165), 1X B-27 Supplement (Thermo Fisher Scientific, #12587001), 500 nM A83-01 (Tocris Bioscience, #2939), 1X GlutaMax Supplement (Thermo Fisher Scientific, #35050061), 10 µM SB-202190 (Biogems, #1523072), 50 ng/mL EGF (PeproTech, AF-100-15), 10 mM Gastrin (Sigma), and 50% Wnt-3A/R-spondin/Noggin conditioned media. Every two weeks organoids were passaged. First media was removed and 500 µl TrypLE (Invitrogen, #12604-012) was added to each well. Organoids in TrypLE were incubated for 10 min and the Matrigel plug was scraped with a pipette tip and all wells were combined by pipetting the cells into a 15 ml tube. Organoids in TrypLE were incubated at 37C in a water bath for 30 min to dissolve the Matrigel and break up the organoids to single cells, before 1 ml FBS was added to neutralize the TrypLE, and the mix was spun down at 600g for 5 min. Supernatant was removed and the cell pellet was resuspended in 1-2 mL Base media for washing and cell counting. After counting 10-20,000 cells per well (**Supplementary Table 2**) were transferred to another 15 mL tube, spun down and resuspended in Matrigel (R&D systems, Basement Membrane Extract type 2) for plating. Either 12 or 24 wells of a 24 well plate were used well (**Supplementary Table 2**). After the Matrigel solidified, 500 µl WENR medium was added with 10 µM Y-27632 (Peprotech, #1293823) and 3 µM CHIR-99021 (R&D Systems, #4223) for *TP53* deficient organoids. For WT organoids FGF10 (Peprotech, #100-26) was added. Media was changed after 7 and 10 days, and organoids were passaged again after 14 days, with few exceptions. Consistent with their diploid karyotype, WT organoid cultures had more limited growth potential following culture freeze thaw than *TP53* deficient lines.

##### Tumor suppressor gene editing in organoids

To simplify downstream phenotypic interpretation, we model *TP53* deficiency rather than specific *TP53* mutations that could induce gain-of-function effects*.* *TP53* and *APC* knockouts

(KO) were established using spinoculation and the pX330 plasmid (https://www.addgene.org/42230/) [56]. Briefly, TrypLE was used to break down Matrigel and dissociate organoids to single cells as in the organoid passaging protocol. Organoids were counted and 500,000-1,000,000 cells were resuspended in 500 µl WENR + Y + CHIR media and plated in one well of a 24 well plate. Separately, 10µl Lipofectamine 2000 (Invitrogen, #11668-019), 90 µl Opti-MEM media (Gibco) and a total of 5 µg pX330 plasmid DNA with either a guide for *TP53* or *APC* (**Supplementary Table 1**) were mixed and incubated 5 min at RT before 100 µl of the mixture was added to the previously plated cells. Cells were spun at 600g at 32 °C for 60 min and then the plate was incubated in 37 °C for 4 hours before they were transferred to a 1.5 ml Eppendorf tube, spun down at 600g for 5 min and resuspended into 250-500 µl Matrigel and plated into 6-12 wells of a 24 well plate. After Matrigel polymerization WENR media + Y + CHIR was added. Single CRISPR/Cas9-edited organoids were picked following selection via addition of 10µM Nutlin-3 (for *TP53* loss) and removal of *WNT* and *RSPO* (for *APC* loss) and expanded [27]. Selection was initiated 7 days after spinoculation. Nutlin-3 did not exert an anti-proliferative effect on *TP53* mutants, indicating that the subclones are functionally p53 deficient. To ensure clonal organoid cultures, single clones were picked ~21 days after spinoculation, before the culture had been passaged [57]. Clonality was verified by Sanger sequencing and Tracking of Indels by Decomposition (TIDE) analysis [58] at multiple timepoints during the course of the experiment. For D1C2 and D1C3 the *TP53* KO used different guides and was performed at an earlier time point compared to the seven other cultures. As expected, CRISPR/Cas9 induced KO of *TP53* and *APC* resulted in reduced expression of these genes (**Fig. 3d, Extended Data Fig. 7c,d**), although residual expression may occur [59].

#### Hematoxylin and eosin staining

For the purposes of hematoxylin and eosin (H&E) staining only, organoids were grown using the air liquid interface (ALI) 3D culturing method [60] for 2 weeks and were fixed using 2% paraformaldehyde/PBS (Electron Microscopy Sciences; #15714) for 30 minutes at room temperature. Following fixation, organoids were washed three times with 1x PBS for 5 minutes each, embedded in HistoGel (Thermo Scientific, HG-4000-012), and transferred to 70% ethanol. Organoids were paraffin embedded and sectioned at 5 µm thickness. Paraffin-embedded sections were deparaffinized and rehydrated prior to H&E staining.

#### Genomic sequencing

Genomic DNA was extracted from cellular pellets with DNeasy Blood and Tissue Kit according to the manufacturer's recommendation. The NEBNext Ultra II FS DNA Library Prep Kit was used to prepare libraries for shallow whole genome sequencing (sWGS). For WGS, the NEBNext Ultra II FS DNA Library Prep Kit was adapted for use with IDT duo indexes. Genomic DNA (500 ng per sample), was fragmented, end-repaired, and poly-A tail was added in a single thermocycler reaction using 1 µl NEBNext Ultra II FS enzyme mix and 3.5 µl buffer for 7.5 min at 37 °C, 30 min 65 °C to achieve a fragment length of 300-400bp. IDT indexes, 1.25 µl, were directly added to the resulting FS product along with 15 µl ligation master mix and 0.5 µl enhancer and ran on the thermocycler for 15 min at 20 °C. The resulting product was cleaned up using Zymo clean up kit followed by size selection with Zymo size selection kit to retrieve 300bp fragments and above. The entire eluted product, 15 µl, was then PCR amplified with 25 µl NEBNext Ultra II Q5 Master Mix and 10 µl PPC from Illumina TrueSeq. The protocol included: 98 °C for 30 sec; 7 cycles of: 98 °C for 10 sec, 65 °C for 75 sec; 65 °C for 5 min; 4 °C for 5 min. The PCR product was then purified with Zymo clean up kit and the resulting product was size selected with Zymo size selection kit to retrieve fragments 300bp and above. The target depth was 0.2X for sWGS and 25X for WGS. Given the clonal and epithelial origin of the HGOs, 25X depth was adequate for calling somatic alterations at individual timepoints, as detailed below. While deeper sequencing may reveal additional subclonal SNVs and CNAs, it is unlikely to alter the overall landscape of events. Instead, we investigate single cell copy number states through inference on single cell RNA-sequencing (scRNA-seq) data and linkage to lineage using expressed cellular barcodes (ECB), as described below.

In addition to sWGS which was performed on all HGO cultures throughout the multi-year experimental time course, several samples were profiled via whole exome sequencing (WES, using the IDT xGen Exome Hub Panel v2) during initial QC assessment. These data were used to confirm the CRISPR/Cas9 edit sites (**Supplementary Fig. 2-3**) and a truncating *CDKN2A* (P16) mutation (D1C3, **Supplementary Fig. 9a**).

#### FACS analysis of genomic content

First media was removed and 500 µl TrypLE (Invitrogen, #12604-012) was added to each well. Organoids in TrypLE were incubated for 10 min and the Matrigel plug was scraped with a pipette tip and all wells were combined by pipetting the cells into a 15 ml tube. Organoids in TrypLE were incubated at 37C in a water bath for 30 min to dissolve the Matrigel and break up the organoids to single cells, before 1 ml FBS was added to neutralize the TrypLE, and the mix was spun down at 600g for 5 min. Supernatant was removed and the cell pellet was resuspended in staining buffer (2% inactivated FBS in PBS with 0.09% sodium azide filtered with 0.2 micron filter). After cell counting, 1 million cells in 100 μl staining buffer were aliquoted to a well in a 96 well plate, and 5 μl of DAPI staining solution (Fisher, #D1306). Samples were analyzed on the Aurora Flow Cytometry System (Cytek Biosciences), and resulting data analyzed by FlowJo.

##### Generation and transduction of expressed cellular barcodes (ECBs)

##### Production of the expressed cellular barcode (ECB) library

The expressed cellular barcode (ECB) consists of a stretch of 30 semirandom DNA bases, alternating between the weak bases "A" or "T" and the strong bases "C" or "G", similar to Clontracer [61]. Differences include 1) use of a different plasmid backbone: pCDH-EF1-MCS-BGH-PGK-GFP-T2A-Puro cDNA Cloning and Expression Vector (System Biosciences, #CD550A-1), and 2) the Clontracer sequence was inserted downstream of a strong promotor (EF1-alpha) and upstream of a poly-A tag, and 3) the barcode insert was annealed instead of created by extension reaction 4) to increase barcode complexity bacteria were grown on large agar plates instead of in suspension. The pCDH-EF1-MCS-BGH-PGK-GFP-T2A-Puro was modified by first destroying the ClaI restriction site, by ClaI-HF restriction enzyme (New England Biolabs), according to manufacturer's instructions (**Supplemental Fig. 15a**). The edges of the linearized plasmid were then dephosphorylated by Antarctic Phosphatase (New England Biolabs), according to manufacturer's instructions. The plasmid was linearized again using oligos Cla1_dest_O1 and Cla1_dest_O2 (**Supplementary Table 1**) and Quick Ligase (New England Biolabs) according to manufacturer's instructions. This process was then repeated to adjust the multiple cloning site to insert ClaI and XhoI restriction sites, using restriction enzymes BamHI and NheI (New England Biolabs). Dephosphorylation was performed with Antarctic Phosphatase (New England Biolabs) and ligation performed using oligos MCS_dest_O1 and MCS_dest_O2 (**Supplementary Table 1**) and Quick Ligase (New England Biolabs). Next a molecular stuffer (1.3 kb) was inserted using ClaI and XhoI restriction enzymes (New England Biolabs), with the dephosphorylation and ligation reactions performed as above. Each restriction enzyme step can be found in the file "ECB_library.dna" on https://github.com/cancersysbio/gastric_organoid_evolution. ECB barcode insertion was performed as follows: the two oligonucleotides ECB_forward and ECB_reverse (**Supplementary Table 1**) were combined with 10 ml of each (100 mM), with 80 ml $H_20$, and annealed by heating to 95 °C in a thermocycler, and then cooled 1 °C per minute to 25 °C. Meanwhile, the plasmid "pCDHcopGFPPuroClonoseq" was mixed with restriction enzymes ClaI (New England Biolabs) and XhoI (New England Biolabs) in 1X Cutsmart buffer (New England Biolabs), to linearize it. The reaction was incubated at 37 °C for 60 min, followed by heat inactivation of the restriction enzymes at 65 °C for 20 min. 5' phosphate was removed using Antarctic Phosphatase (New England Biolabs) according to the manufacturer's instructions. The resulting linearized plasmid was gel purified and eluted in 20 µl $H_20$. Next, 50 ng of the linearized plasmid was ligated with 1 ml of the annealed ECB oligonucleotides using Quick Ligase (New England Biolabs, #M2200S). The reaction mix was incubated at room temperature for 5 mins and then purified with AmPure XP (Beckman Coulter, # A63881) according to the manufacturer's instructions. The purified plasmid, the ECB plasmid, was quantified using NanoDrop. The efficacy of the ligation reaction was tested by transforming One Shot™ Stbl3™ Chemically Competent E. coli (Invitrogen) according to manufacturer's recommendations. 10 colonies were picked and the correct plasmid product was verified by sanger sequencing using the ECB_plasmid_sanger primer (**Supplementary Table 1**). Ideally 80% or more of the colonies should have the correct plasmid product.

For plasmid expansion, 4 large plates (Molecular Devices, # X6023), were covered with 1L Luria Agar and Carbenicillin (concentration: 100 µg/ml). The plates were cooled down at 4 °C overnight and the following day, the plates were pre-warmed at 37 °C. Following this, two vials of Endura Electro Competent Cells (Lucigen, #60242-1) were thawed on ice and 2 µl of the ECB plasmid was added to each vial. Bacteria and plasmids were mixed and incubated on ice for 30 min prior to transfer to four prechilled 0.1 cm width (Bacteria) Electroporation Cuvette with White Cap (USA Scientific, # 9104-1050), 25 µl per cuvette, for electroporation using the settings 1.8 kV, 600 ohms, and 10uF (Gene Pulser Xcell, BioRad). Immediately after electroporation 1 ml Lucigen recovery media was added to each cuvette and bacteria from all 4 cuvettes were transferred to a 14 mL round bottom culture tube. Cells were then shaken at 250 rpm at 37 °C for 2 hours. Following this, 1 ml of bacterial cells were then plated on each plate, and plastic beads were used to spread out the bacteria. Plates were incubated overnight (>18 hours) at 37 °C. Next day for each plate 20 mL Luria Broth was added to the plates and bacteria was scraped and collected into two 50 ml tubes. The bacteria were then centrifuged at 4000 rpm for 20 min, and the plasmid purified using Qiagen Maxiprep.

**Production of ECB lentivirus**

The ECB plasmid were co-transfected with lentiviral packaging plasmids pLP1 and pLP2 and viral envelope plasmid VSV-G into 293T cells by polyethylenimine (PEI) into HEK293 cells. Briefly, the day before transfection 3,500,000 cells were plated and 10 mL media (DMEM/10% FBS) was added. The next day a mixture of 2 mL OptiMem (Thermo Fisher Scientific, #31985062), 25 µl PEI (dissolved at a concentration of 50g/L), 4 µg ECB plasmid and 1.3 µg each of plasmids pLP1, pLP2 and vsv-g was incubated 20 min at RT, before being added dropwise to the cells. Lentiviral supernatants were collected at 48 hours and 72 hours post-transfection. The collected media was spun down at 1000g for 10 min at 4 °C, then supernatant was transferred to a new tube and spun down again at 1000g for 5 min at 4 °C. The supernatant was collected, and PEG was added at a final concentration of 8% and NaCl at a final concentration of 0.4M. The mixture was centrifuged at 1500g for 30 min at 4 °C. After the supernatant was aspirated, the lentiviral particles were washed with ice-cold PBS, resuspended in PBS and stored at -80 °C. Sequencing of the plasmid barcode showed no sign of skewing (**Supplementary Fig. 15b**).

**Lentiviral transduction of ECBs in organoids**

Organoids were transduced with the ECB lentivirus, with a multiplicity of infection (MOI) of 0.1, ~115 days after TP53-KO when sufficient biomass was available. Approximately, five million cells were dissociated into smaller clusters with TrypLE (Invitrogen, #12604-012) for 15 min at 37 °C. Following this, clusters were resuspended into 500 µL transduction solution containing 8 µg/mL polybrene (Sigma, #107689), 3 µM CHIR-99021 (R&D Systems, #4223), 10 µM Y-27632 (Peprotech, #1293823) and concentrated lentivirus in organoid culture media. Spinoculation of resuspended organoids was performed at 600 g for 1 hour at 32 °C. After spinoculation, organoids were incubated for 12-14 hours at 37 °C and replated onto a new 24-well plate.

##### Culturing ECB transduced organoids

Following lentiviral transduction of organoids with the ECB library, cells were expanded for 28 days and then FACS sorted for GFP expression aiming for at least 100,000 cells. D1C1 yielded 37,000 GFP+ cells, D2C1, D2C3 and D3C2 yielded 100,000, and D2C2 300,000 cells. Cells were re-expanded for 28 days and divided into three replicates that were monitored for up to 40 weeks. Every two weeks (with few exceptions), organoids were dissociated to single cells, counted, imaged, stored, barcode sequenced and passaged with 10,000 or 20,000 cells per well. Details for each culture are provided in **Supplementary Table 2**.

##### ECB DNA sequencing

Barcodes were isolated by amplification of the region from genomic DNA in a PCR reaction using 2X KAPA Hifi PCR Master Mix (Roche Sequencing Solutions, #KK2601) with a minimum of 200 ng of DNA input per sample. The PCR product was then purified using Ampure magnetic beads (Beckman Coulter, #A63881) and eluted in 12 µl, 2 µl of which was used in a second PCR reaction to amplify the isolated barcode sequence and add adaptor sequences. KAPA HiFi Hot start was used in this PCR reaction and 6bp identification indexes were integrated in the reverse primer (**Supplementary Table 1**). The PCR product of all samples was then combined and purified using Qiagen PCR purification kit. To remove primer dimers, the purified product was run on a 2% EX gel (Thermo Fisher, #G401002) and the 200 bp band was isolated and gel purified by Qiaquick gel extraction kit (Qiagen, #28706). The purified product was bioanalyzed and confirmed for a 204bp peak prior to sequencing with 150bp paired-end reads on an Illumina MiSeq. ECB barcode replicates were established for 5 cultures (D1C1, D2C1, D2C2, D2C3, D3C2).

#### Single-cell RNA sequencing

##### Preparation of cell Hash tags

Cell 'hashing' with barcoded antibodies enables multiplexing and doublet detection for single cell genomics [62]. We modified the original protocol slightly by using (1) EpCAM as the antibody, (2) different oligonucleotide adaptor sequences (**Supplementary Table 1**) and (3) a custom bioinformatics pipeline to link hashtags to cells. Briefly, the EpCAM antibody was conjugated to streptavidin using LYNX Rapid Streptavidin Antibody Conjugation Kit in a 10µg streptavidin to 15µg antibody reaction. The conjugated antibody was then combined with 800 pmol of pre-biotinylated 66-bp-long polyA-tailed index oligonucleotides with phosphorothioate at the 3'end purchased from IDT (**Supplementary Table 1**).

##### 10X scRNA sequencing

Organoids were thawed in a water bath, added to a 15 ml tube containing 10 ml FBS, spun down for 5 min at 600 x g and resuspended into 2 ml TrypLE and incubated at 37C for 10 min. If the organoids were close to single cell suspensions, 1 ml FBS was added and cells were pelleted for 5 min at 600 x g. If they were not yet suspended as single cells, they were incubated for 10 min in TrypLE before being centrifuged. All single-cell RNA-sequencing (scRNA-seq) assays utilized cell hashing for multiplexing [62]. Single cells were suspended in a staining buffer (PBS with 2mM EDTA and 2% heat inactivated FBS) at 1,000,000 per 50µl buffer and relevant antibody-oligo conjugates were added at 1 µg per antibody. The volume was brought to 100 µl using the same staining buffer and cells were incubated with the conjugates for 1 hour on ice. To remove unbound antibodies, cells were washed with staining buffer 3 times and re-suspended in appropriate volume for FACS sorting at the final wash step. Cell suspensions were strained and FACS sorted to obtain single cell suspensions before loading onto the Chromium instrument (10X) for 3' scRNA-seq (v3 chemistry). 10X sequencing was performed at the Stanford Functional Genomics Facility according to the manufacturer's recommendation except that during the cDNA amplification step, a primer was added to amplify the hashtag (HashTag cDNA amplification primer, **Supplementary Table 1**). Additionally, for cultures that had been transduced with the ECB virus, a primer to amplify the ECB tag (ECB cDNA amplification primer, **Supplementary Table 1**) was added. During the bead separation after cDNA amplification, the supernatant that contained the shorter hash and ECB tags were collected and further purified using AmpureXP, as described previously [62].

**Hash tag library preparation**
Hash tags were amplified as a separate library using 2X KAPA Hifi PCR Master Mix (Roche Sequencing Solutions, #KK2601) according to the manufacturer's recommendations, and with 12 PCR cycles. The hash tags were amplified with adaptors containing primers specified in **Supplementary Table 1**. The libraries were purified using AmpureXP (Beckman Coulter, # A63881) at 2X concentration.

**ECB cDNA library preparation**
ECB tags were amplified as a separate library in two reaction steps. First, ECB tags were amplified using 2X KAPA Hifi PCR Master Mix (Roche Sequencing Solutions, #KK2601), per manufacturer's recommendations, with 15 PCR cycles. Primers ECBseq PCR 1 forward and reverse were used (**Supplementary Table 1**). The reaction was purified using MinElute PCR Purification Kit (Qiagen, # 28006). Next, adaptors were added to the amplified ECB tags using 2X KAPA Hifi PCR Master Mix again, utilizing index adaptors specified in **Supplementary Table 1**. This reaction was again purified using MinElute PCR Purification Kit, and then further purified using E-Gel EX Gel 2% (Thermo Fisher, #G402002). Typically, we detected two peaks at 450 and 550 bp, which were extracted and gel purified using QIAquick Gel Extraction Kit (Qiagen, # 28704). Both the Hash and ECB libraries were quantified using Qubit (Thermo Fisher) and size verified via Bioanalyzer (Agilent).

#### Quantification and statistical analysis

**Organoid growth curve derivative estimates**
To estimate growth curve derivatives of ECB-specific subclones (detailed below) we used cell counts and seeding dilution information (obtained for each culture at each passage) to generate a whole-culture growth curve (as a function of time). To reduce the effects of noise in cell counting, we approximated the growth curve using locally estimated scatterplot smoothing (LOESS) with the R function loess (span=2). The relative abundance of each ECB subclone at each time point (derived from the relative fraction of reads corresponding to its barcode in targeted DNA sequencing of the ECBs) was used to estimate its population size at that time point:

$$log_{10}[S_{i,t}] = log_{10}[S_t] + log_{10}[f_{i,t}] \quad (1)$$

Where $\boldsymbol{S_{i,t}}$ is the estimated population size of subclone i at time t; $\boldsymbol{S_t}$ is the whole-culture population at time t; and $\boldsymbol{f_{i,t}}$ is the relative abundance (fraction of the population) of subclone i at time t. These values were then used to plot an estimated growth curve (using loess with span=2) for each ECB subclone, and the derivatives across time were calculated.

To assess changes in whole culture growth rates along a broad time span, we estimated growth curve derivatives in Early-to-Late and Early-to-Mid time point trajectories. Unlike the growth curve estimation for shorter-term cultures, these trajectories are based on the expansion of cultures which were frozen and thawed at different time points (requiring time to initially recover from thawing) and contained gaps in counts or missing passage information (**Supplementary Table 2**). Therefore, growth curves were estimated based on relative/interpolated passage numbers, where missing time points were removed and whole-culture population sizes were calculated using time points for which complete data exists, assuming a fixed (14-day) interval between passages. While this interval is not entirely accurate along the complete time course (due to missing time points), we found the resulting growth curves to be smoother and more robust compared to imputation of these missing points. Furthermore, to reduce local noise/bias effects due to inaccuracies in time at specific points, we again approximated the growth curve with a smooth curve (loess, span=2) prior to calculating the derivative. All cultures exhibited consistent growth increases throughout the time course, as observed by the log-linearity of their growth curves (**Fig. 3b**).

**Evaluation of mycoplasma levels and the association with molecular features**
Through rigorous and quantitative analysis, early passage wild-type (WT) cultures and some derivative samples were found to be infected with mycoplasma at varied levels. In particular, we systematically analyzed mycoplasma levels across the experimental time course by mapping WGS data from all samples to six mycoplasma genomes: *Alaidlawii, Arginini, Fermentans, Hominis, Hyorinis* and *Orale*. *M.* The dominant strain, *M. hyorhinis,* is common in gastrointestinal tissues and thought to infect the stomach via consumption of porcine products [63]. Such naturally occurring microbes can be transmitted to organoid culture, and while patterns are suggestive of donor origin, laboratory contamination cannot be ruled out. When present, mycoplasma typically accounted for less than 1% of reads (with the exception of D1C1 which was 3.7% at an early time point, day 55) (**Supplementary Fig. 7a-f, Supplementary Table 3**). Prior studies have assessed mycoplasma positivity based on RNA-seq data with a cutoff of 100 reads per million (RPM) [64]. Here we chose to use a stringent cutoff of 30 or more mycoplasma RPM to call a sample mycoplasma positive from sWGS data (**Supplementary Fig. 7d**).

Importantly, and in contrast to *H. pylori* [65] and Epstein Barr Virus (EBV) [66], which are established risk factors for gastric cancer, *M. hyorhinis*, is not known to produce potent genotoxins, to have strong transformational potential, nor is it considered a human pathogen [67]. While some mycoplasma strains such as *M. fermentans* have been reported to induce transformation in immunodeficient models, these are not typical strains that infect cells grown in the lab [68]. Since the objective of this study was not to examine clonal evolution in the presence of bacterial infections, when mycoplasma was detected, cultures were treated with normocin to eliminate the infection.

Moreover, to systematically evaluate the potential impact of mycoplasma infection on the biological processes of interest, a major portion of the study was repeated by evolving all original *TP53*$^{-/-}$ cultures sampled at *Early/Mid* (190-290 days) and *Late* (540-730 days) time points under mycoplasma-free conditions through constant antibiotic (normocin) treatment. These cultures, thawed from frozen stocks, are referred to as *Mid* and *Late* trajectories, respectively (**Extended Data 1, Supplementary Fig. 7**). For each culture, multiple cellular phenotypes were assessed, including organoid growth, genomic, and single cell transcriptomic profiles over time, providing high resolution into clonal dynamics and transcriptional profiles in the context of *TP53* deficiency. These experiments enable direct comparison of the evolutionary trajectories under mycoplasma-free conditions with the original cultures harboring intermittent and variable mycoplasma levels, as well as the impact of antibiotic therapy on organoid growth, and their molecular profiles.

Of note, all cultures appeared healthy with consistent growth increases throughout the time course (**Fig. 3b, 5d**), irrespective of mycoplasma, which has been reported to slow cell growth. The presence of mycoplasma did not correlate with copy number aberrations (CNAs) in *TP53* deficient cultures (**Supplementary Fig. 7g**). Moreover, convergent copy number evolution was observed, with nearly identical breakpoints present in mycoplasma free versus infected samples as illustrated for specific samples (**Supplementary Fig. 8**). More generally, despite the distinct evolutionary trajectories of each *TP53*$^{-/-}$ engineered culture, the overall molecular patterns recapitulate those seen *in vivo* and are enriched amongst chromosomally instable (CIN) gastro-esophageal cancers (**Fig. 2C, Extended Data 2b-d, 3e,4a**). These molecular patterns are consistent with those seen in murine gastric organoids upon co-deletion of *TP53* and *CDKN2A* [19,69]. In contrast, wild-type *TP53* proficient HGOs remain genomically stable (**Supplementary Fig. S4-6**) and were transcriptionally similar to normal gastric tissue (**Fig. 3**).

We also investigated the potential effect of mycoplasma on single cell transcriptional profiles and clonal fitness. No significant pathway alterations were observed when comparing the *original trajectories* (which includes samples with variable mycoplasma levels) relative to comparable mycoplasma negative cultures (termed *Mid* and *Late trajectories*), emphasizing that these patterns are not driven by mycoplasma (**Supplementary Fig. S19b**). Moreover, comparisons of the winning subclone (which achieved clonal dominance) and all other subclones was highly concordant between the *original trajectories* versus *Mid* and *Late trajectories* (**Supplementary Fig. 19c,d**).

Thus, these analyses do not identify significant effects for mycoplasma exposure or for antibiotic therapy on organoid growth, genomic and transcriptomic profiles or fitness determinants across winning subclones. Rather, similar patterns were observed across different tiers of replication, across donors in non-barcoded and barcoded experiments and irrespective of mycoplasma levels or antibiotic treatment. The striking degree of reproducibility in this system is even more notable given the potential sources of technical and biological variation and implies that any such effects are evidently *modest* relative to the overwhelmingly dominant effect of *TP53* inactivation. Given the strong concordance observed across these and the multiple tiers of replication in this study (spanning distinct donors, distinct *TP53* CRISPR/Cas9 edited clonally derived lines, barcoded and non-barcoded cultures, as well as parallel experiments with normocin treatment), we focus on the original trajectory for simplicity, and cross-reference these comparisons.

While mycoplasma is evidently not the dominant factor in this system, the contribution of this and other sources of technical variation (media batches, matrigel digestion duration, donor differences, other microbes not routinely tested) cannot be ruled out. Future experiments conducted with and without constant antibiotics will be needed to address these questions. However, prior studies have suggested that antibiotics are not necessarily mycoplasmacidal in long-term culture or always effective [70]. Additionally, given that antibiotics such as fluoroqinolones, which target bacterial type II topoisomerases, can be genotoxic [71,72], evaluation of their long-term effects will be important as will evaluation of the outgrowth of other microbes. Since human tumors develop and evolve in the context of near constant exposure to microbes, understanding the relationship with the host is of biological relevance, and should be the focus of future investigation.

**Analysis of sWGS and WGS data**

Shallow WGS (sWGS) raw fastq files were processed using the Nextflow-based pipeline nf-core/sarek [73] v2.7.1 with BWA v0.7.17 [74] for sequence alignment to the reference genome GRCh38/hg38 and GATK [75] v4.1.7.0 to mark duplicates and calibration. The recalibrated reads were further processed and filtered for mappability and GC content using the R/Bioconductor quantitative DNA-sequencing (QDNAseq) v1.22.0 [76]. Copy number alterations (CNAs) were called using QDNAseq in R v3.6.0 with 50-kb bins obtained from (https://github.com/asntech/QDNAseq.hg38). We analyzed in total 140 sWGS samples and 20 WGS samples sampled along the experimental time course (**Extended Data Fig. 1**, **Supplementary Tables 3, 4**).

**Fraction genome altered**

To reduce noise in the CNA profiles estimated from sWGS, a moving average of 25 windows was applied to the 50kb window output from QDNAseq (see above) to compute the fraction genome altered (FGA). The data was exponentiated back to read counts and centered around 1. A window was called as altered according to **equation 2**:

$$CN = \begin{cases} CN_w \leq \overline{CN} \times 0.75 \mid 0\ (Loss) \\ \overline{CN} \times 0.75 < CN_w < \overline{CN} \times 1.25 \mid 1\ (Diploid) \\ CN_w \geq \overline{CN} \times 1.25 \mid 3\ (Gain) \end{cases} \tag{2}$$

Where CN is the assigned copy number, $CN_w$ the copy number value for the window of interest and $\overline{CN}$ depicts the average copy number. As relevant to this analysis, the y-axis in **Fig. 1d** denotes the percentage of 50kb windows that are altered.

**Timing of copy number alterations**
The time of appearance of CNAs was estimated using a similar approach as for computing the fraction genome altered, although the mean was calculated across whole chromosome arms instead of 50kb windows. For each time point, the chromosome arm was called altered according to **equation 2** above. Only CNAs that were consistently altered after the first identified time point were retained. Thus, if a chromosome arm alteration was present in a subclone and became extinct, it was not taken into consideration.

**Timing of bi-allelic deletions**
Biallelic deletions were called from sWGS when a 50 kb window possessed 25% of normalized reads compared to the average of all windows, as shown in **equation 3**. If such a window occurred within the *FHIT* or *CDKN2A* regions, these genes were considered as having bi-allelic loss. Only alterations that were consistently altered after the first identified time point were retained.

$$CN = \{CN_w \leq \overline{CN} \times 0.25 \mid 0 \ (Bi - allelic\ loss) \qquad (3)$$

Where CN is the assigned copy number, $CN_w$ the copy number value for the window of interest and $\overline{CN}$ represents the average copy number.

**SNV, SV, CNA and ploidy calls from WGS**
A subset of cultures and timepoints were subject to deeper WGS (as summarized in **Extended Data Fig. 1**, **Supplementary Fig. 1, Supplementary Table 4**), facilitating single nucleotide variant (SNV) calling. Short reads produced by WGS on the Illumina platform were aligned to hg38 using BWA (v0.7.17) [74]. Following the GATK (v4.1.4.1) best practice workflow, raw alignment files (bam) were pre-processed by marking duplicated reads and base recalibration. SNV calls were made using MuTect2 and FilterMutectCalls from the GATK package. Both "PASS" mutations and "multiallelic" mutations were retained. To ensure high confidence INDEL calls, two methods were used: 1) MuTect2 and FilterMutectCalls from the GATK package and 2) Strelka (v2.9.10) [77] and, only INDELS called with both filters were retained.

PURPLE (v3.0) was used to estimate the copy number states, purity, LOH and ploidy of organoid cultures, comparing matched wild-type (WT) and *TP53*, *TP53/APC* KO cultures with WGS data. PURPLE implements B-allele frequency (BAF) from AMBER (v3.3), read depth ratios from COBALT (v1.8) [78,79]. For each sample and each of the 22 autosomal chromosomes, the percentage of gained and lost genomic material was calculated relative to the ploidy of the sample. The whole genome instability (wGII) score was calculated as the average of this percentage over the 22 autosomal chromosomes [80]. Additionally, CNAs were called using QDNAseq v1.22.0 [76] as described above.

To further investigate whether *TP53* deficient HGO cultures underwent genome doubling, ploidy was estimated via TitanCNA, FACETS and Hatchet on paired KO and WT organoid BAM files. TitanCNA (v1.24.0) [81] was run via the snakemake workflow. FACETS (v0.6.1) [82] used the following parameters: --purity-cval=1000 and –cval=500. Hatchet (v0.2.10) [83] was run with dbSNP151.GRCh38 as a reference to call heterozygous germline SNPs from the matched-normal sample and Michigan imputation server v1.2.4 [84] against population AMR (1000g-phase-3-v5) to phase the SNPs. The original files in GRCh38/hg38 format were converted to GRCh37/hg19 format for the phasing and then converted back with LiftoverVcf in package Picard v2.18. These analyses revealed limited evidence for tetraploidy and genome doubling.

Somatic structural variants (SVs) were called from WGS data using four tools: Manta (v1.6.0) [85], Delly (v0.8.1) [86], GRIDSS (v2.9.4) [78,79] and SvABA (v.1.1.3) [87]. Consensus calls were made by comparing the output of GRIDSS and the other three tools, with a maximum allowed distance of 100bp as measured pairwise between breakpoints. SV calls from the Paulson et al. BE progressors and non-progressors were based on a similar consensus approach [16]. Distinct rearrangement states were called in *TP53*$^{-/-}$ HGOs based on junction balance analysis using JaBbA (v1.1) [34]. This analysis identified *rigma-like* deletion chasms in *TP53* deficient organoid cultures (D3C1), similar to those seen in GC, as shown for pfg008 from [23] (**Supplementary Fig. 9d**). We used ClusterSV (https://github.com/cancerit/ClusterSV) to define SVs as simple or complex [88]. Briefly, simple SVs were defined as having 2-9 rearrangements while complex SVs contained 10 or more rearrangements and others are considered non-clustered SVs.

**COSMIC analysis**

SNV calls from WGS were annotated using Gencode version 38 and the VariantAnnotation package in R [89]. Specifically, identification of mutations in coding genes was made using the function "predictCoding". For identification of gene locations, the "locateVariants" function was utilized. To identify coding mutations in cancer associated genes, the output from predictCoding was filtered based on whether the gene was included in the Catalogue Of Somatic Mutations In Cancer (COSMIC), version 94 [90].

**Mutational signature analysis**

Mutational signature analysis was performed using the SNV data from WGS. Single-base substitution (SBS) signatures were extracted using the HDP package in R (https://github.com/nicolaroberts/hdp) based on a hierarchical Bayesian Dirichlet process. De novo signatures were extracted and utilized to run an expectation-maximization algorithm in order to deconvolve these signatures into known signatures from the pan cancer analysis of whole genomes [91]. The results were manually inspected and the *sigfit* [92] package in R was used to reconstruct donor-specific mutation spectrums, yielding a cosine similarity ≥ 0.9 in all samples. The signature contribution per sample was visualized using a custom R script.

**Fishplot schematic visualization of subclonal copy number evolution**

To reduce noise in CNAs, a moving average of 25 windows was applied to the 50kb window output from QDNAseq analysis of sWGS data. The data were exponentiated back to read counts and centered around 1. A copy number subclone was defined by having at least 100 consecutively altered 50kb windows, where each window conformed to **equation 4** below:

$$CN = \begin{cases} CN_w \leq \overline{CN} \times 0.875 \mid 0\ (Loss) \\ \overline{CN} \times 0.875 < CN_w < \overline{CN} \times 1.125 \mid 1\ (Diploid) \\ CN_w \geq \overline{CN} \times 1.125 \mid 3\ (Gain) \end{cases} \quad (4)$$

Where CN is the assigned copy number, $CN_w$ the copy number value for the window of interest and $\overline{CN}$ represents the average copy number. The number of consecutive windows and percentage gain or loss required were determined based on visual inspection that these settings retained most copy number based subclones. Subclone copy number was determined as the average of all subclone windows multiplied by 2. The subclone cell frequency was determined as the absolute value of copy number deviation from the diploid copy number state. Thus, a chromosome arm with a copy number of 2.65 was interpreted as a copy number subclone with an allele gained with a prevalence of 65% in the cell population. Regions that co-varied over time were assigned to the same subclone. For fish plots with ECB-based subclones (e.g. **Fig. 6c**), all non-highlighted subclones were grouped into "other", and each subclone fraction was multiplied with 1000, shifted into $\log_2$ scale, and then re-scaled back to "fraction". The frequencies of copy number subclones within an ECB subclone was estimated from inferCNV based on the scRNA sequencing data. The figures were plotted with the Fishplot package in R [93]. The implementation of this required adjustment of cell frequencies, such that if two subclones from a shared parent had a higher frequency than the parent subclone itself, this would not be accepted. Cell frequency adjustments and determination of parent to child subclone relationships, were made manually. To facilitate reproduction of these schematics, cell frequencies and adjustments for each culture are provided (**Supplementary Tables 5, 8**).

**Comparison between organoid cultures and tumor copy number alterations**
Arm-level copy number alterations from the Cancer Genome Atlas (TCGA) data was downloaded from http://gdac.broadinstitute.org/, and the files "broad_values_by_arm.txt" based on Affymetrix Genome-Wide Human *SNP* Array 6.0 were used. These files report chromosome arm amplification levels for each sample in units of absolute copy number (centered at 0). A chromosome arm was called as lost or gained based an increase or decrease of 0.25 in absolute copy number. Copy number alterations (CNAs) for organoid cultures were calculated similarly as in FGA (equation 2), where alteration status was calculated for each 50kb genome window. For a chromosome arm to be called as gained or lost, at least 25% of windows for that arm were required to be called as gained or lost.

#### Analysis of scRNA-seq and linked ECB data

**Quality control and scRNA-seq data pre-processing**
The raw base call files from the 10X Chromium sequencer were processed utilizing the Cell Ranger Single-Cell Software Suite (release v3.0, https://support.10xgenomics.com/single-cell-gene-expression). First, the "cellranger mkfastq" command was used to demultiplex the sequencing samples and to convert barcode and read data to fastq files. Based on the fastq files, "cellranger count" was executed to perform alignment, filtering, as well as barcode and unique molecular identifier (UMI) counting. The reads were aligned to the hg38 reference genome, implementing a pre-built annotation package downloaded from the 10X Genomics website. Several output files including a barcoded binary alignment map (bam) file and a summary csv file are provided. Importantly, a filtered feature-barcode matrix folder, containing a valid barcode file for all QC-passing cells, a feature file with ensembl gene ids and a matrix in the genes x cells format are generated. The filtered genes x cells matrix was further used as input for the data processing workflow described in the following.

**Computing subclone frequencies from ECB sequencing**
ECB barcodes were extracted from the ECB fastq files. Only reads exactly matching the sequence around the 30 base-pair barcode were retained to ensure read quality. Then for all cultures from a single transduction event (e.g. parent and replicate cultures from D2C2), the number of reads per barcode were combined into a single file. This was utilized as input to UMI-tools [94], which merged barcodes that were similar, i.e. where the difference between barcodes was more likely to depend on sequencing and PCR errors as compared to being a separate barcode. Each such group of combined barcodes was called a read group. The file containing all read groups (RGs) was called "ECB_groups_master_list".

For most replicate cultures, scRNA-seq was performed at multiple time points, allowing flagging of cells where two or more barcodes were inserted during transduction. In these cases, one early scRNA-seq time point was used to identify the multiple insertion events, and if such events were identified, RGs in the ECB_groups_master_list were combined. Note that the same updated master list was utilized to convert barcodes to RGs for all replicate cultures from the same transduction experiment. In addition, this master list was used for the cDNA-based ECB barcode analysis. The following section describes how multiple transduction events in a single cell were handled.

The data were assembled into a read group count matrix (RGC matrix), with RGs as columns and time points as rows. The RGC matrix was average normalized across rows, to obtain read group frequencies per time point. Next, RGs that had a frequency of less than 0.1% at all time points were removed, and the RGC matrix was average normalized across rows again so that the frequencies per time point sum to 1. The barcode frequency data was visualized as Mueller plots using custom R scripts.

**Computational association of hashtags from 10X scRNA-sequencing fastq files**
Although all 10X scRNA-seq was performed using the previously published Hash-seq protocol, results were analyzed by a custom bioinformatics pipeline, enabling the link of hash tags to cells. First, read pairs were filtered according to several criteria: (1) Read 1 started with 28 bases of A, C, G or T followed by fives bases of T, (2) Read 2 contained the adapter sequence "CCTTGGCACCCGAGAATTCCA" conforming to a hamming distance of 2. To link paired reads to a cell, each read included unique information, namely the 10X cell identifier and UMI found in positions 1-16 and 17-26 on Read 1 and the antibody-derived tag (ADT) for cell hashing found in position 21-33 on Read 2. 10X cell identifier reads were kept if they were within 1 hamming distance of the 10X cell identifiers in the 10X Whitelist and only matched with one unique cell identifier. The ADTs were kept if the hamming distance from any designed hash sequence was 2 or lower. Similar UMI reads were clustered and combined using UMItool *clusterer*. For UMIs with multiple ADTs after clustering, the ADT with the most reads was assigned to the UMI. Finally, each 10X cell identifier was associated with the valid ADTs, if it fulfilled the following criteria: (1) at least three unique UMIs mapping to the ADT (UMI counts) and at least three reads per UMI for the respective cell, 2) for 10X cell identifier with multiple ADTs, only ADTs with UMI counts larger than a threshold: number of UMI counts / number of different ADTs * 0.3, were considered valid ADTs, (3) the cell had less than four supporting valid ADTs, and (4) if there were two or three valid ADTs associated with the cell identifier, the ADT with the highest percentage of associated UMI counts was assigned to the cell, unless this ADT has less than 70% of associated UMI counts. In such cases the cell was labelled as ambiguous and therefore discarded from further analysis. This procedure was implemented in a custom python script.

**Computational association of ECBs from 10X scRNA-sequencing fastq files**
Equivalent to the described association of scRNA-sequenced hashtags to fastq files from scRNA-seq, the ECB tags were associated with the 10X cell identifiers. First, the read pairs were filtered according to two criteria: (1) Read 1 started with 28 bases of A,C,G or T followed by fives bases of T and (2) Read 2 contained the adapter sequence "CTGCAGTCTGAGTCTGACAG" in the first 20bp, conforming to a hamming distance of maximum 2. The cell identifier and UMI could be found on positions 1-16 and 17-26 on Read 1 respectively, while the ECB tag are found on position 21-50 on Read 2. A cell UMI was associated with an ECB if a single ECB tag had at least two reads and more than 50% of all reads were associated with that cell UMI. From the longitudinal experiments a set of read groups was known (see "Subclone frequency derived from ECB sequencing"), and only ECB reads with the exact sequence of the predefined ECB read groups were kept. 10X cell identifier reads were kept, if they were within 1 hamming distance of the 10X cell identifiers in the 10X Whitelist and only matched with one unique cell identifiers were kept. Similar UMI reads were clustered and combined using UMItool clusterer (UMItools version 1.01). For UMIs with multiple ECBs after clustering, the ECBs with the most reads were assigned to the UMI.

Each 10X cell identifier was associated with an ECB tag, if it fulfilled the following criteria: (1) at least three unique UMIs mapping to the ECB (UMI counts) and at least three reads per UMI for the respective cell, (2) for 10X cell identifier with multiple ECB, only ECBs with UMI counts larger than a threshold: number of UMI counts / number of different ECBs * 0.3 (when total UMI counts <100), or number of UMI counts / number of different ECBs * 0.2 (when total UMI counts >100), were considered valid ECB, (4) when there were multiple valid ECBs associated with the cell identifiers, the multi-ECB was kept if the number of cell identifiers associated with that multi-ECB was at least 10 and was at least 15% of the total number of cell identifiers associated with each individual ECB. For instance, if the number of cell identifiers associated with ECB1/ECB2 was at least 10 and at least 15% of the total number of cell identifiers were associated with ECB1 alone and with ECB2 alone, ECB1/ECB2 were considered as a valid multi-ECB that integrated at an early lineage rather than a duplet. All read groups with overlapping ECBs were collapsed into one read group. Cells with multi-ECBs that failed criteria 4 were considered duplets and discarded from further analysis. Finally, ECB and ADT were associated by exact matching of 10X cell identifiers. The procedure described was implemented in a custom python script.

**Seurat QC analysis**

The output from the Cell Ranger analysis framework was used as input to a scRNA-seq data analysis workflow, structured around the Seurat software toolkit in R (Seurat v.3.2, https://satijalab.org/seurat/). The filtered barcode matrix which is provided by the Cell Ranger framework was read into R using the "read10X" function from Seurat. Cells were removed based on commonly used scRNA-seq quality metrics, including the number of genes detected per cell, the percentage of reads mapped to mitochondrial or ribosomal genes as well as the number of housekeeping genes assessed from a gene list provided by Tirosh et al. [95]. Hence, cells were flagged as poor quality upon meeting one of the following thresholds: (1) the number of expressed genes was lower than 500 genes, (2) 20% or more reads were mapped to mitochondrial genes, (3) 40% or more reads were mapped to ribosomal genes and (4) there were less than 55 housekeeping genes expressed (**Supplementary Table 6**). In addition, the output from the computational association of hashtags and ECB tags was leveraged to remove cells with ambiguous assignments for the respective sequencing runs. Thus, for sequencing runs with only hashtags, cells flagged as "multiplets" and without information ("NA") were removed. For sequencing runs with ECB tags, cells flagged as "multiplets", as well as cells with no associated ECB tag were excluded from further analysis.

To account for confounding factors resulting in different sequencing depths across cells, SCtransform, a regularised, generalized linear model (GLM) was used with additional covariates for cell cycle difference, mitochondrial and ribosomal percentage as well as sequencing depth [96]. Following the normalization, the "RunPCA" function from Seurat was used to perform principal component analysis (PCA) on each individual single cell expression matrix restricted to the highly variable genes (HVGs) determined by SCtransform. Accordingly, the optimal number of PCs was determined through an elbow plot, and graph-based clustering implemented in the "FindNeighbors" function was executed. To cluster the cells, the Louvain algorithm (https://neo4j.com/docs/graph-algorithms/current/algorithms/louvain/) as implemented in Seurat's "FindClusters" function was used to iteratively group cells together. Next, uniform manifold approximation and projection (UMAP) was performed using the "RunUMAP" function with default settings. To minimize the effect of doublets in our dataset, we utilized DoubletFinder, a method to determine the abundance of doublets (https://github.com/chris-mcginnis-ucsf/DoubletFinder) [97]. Doublets were removed from the Seurat object stored for downstream analysis.

**Analysis of scRNA-seq from paired tumor-normal gastric tissue**

The Sathe et al. dataset was accessed as filtered barcode matrices, corresponding to the output from the Cell Ranger Single-Cell Software Suite provided by the Ji Lab at Stanford University (https://dna-discovery.stanford.edu/research/datasets/) [41]. Peripheral blood mononuclear cells (PBMCs) in the Sathe et al. dataset were excluded in these analyses. The Seurat pipeline described above was similarly run on these data with the same criteria and after running DoubletFinder, the dataset was stored as a Seurat object for downstream analysis. However, in contrast to the analysis of the organoid datasets, this dataset was derived from heterogeneous tumor populations and hence comprised endothelial, immune as well as epithelial cells. Since the organoids investigated in this project solely comprised epithelial cells, endothelial and immune cells were removed. Accordingly, the expression of *EPCAM*, *KRT18*, *MUC1*, *KRT19*, *CDH1* and *CLDN4* was assessed as representative for epithelial cells, while *CD4*, *VIM*, *ACTA2* and *PTPRC* were utilized as markers for non-epithelial cells. After removal of non-epithelial cells, Seurat's "FindNeighbors" and "FindCluster", as well as "RunTSNE" and "RunUMAP" functions were run again. In a final step, the resulting Seurat objects were saved and utilized in the latent semantic indexing (LSI) projection.

**Cell type marker assessment and annotation**

To evaluate the cellular identity of distinct cultures and timepoints, the expression of literature-derived cell marker genes was assessed [40–42]. This list of known marker genes included, but was not limited to, *MUC5AC, TFF1, TFF2* (Pit mucosal cells, PMCs), *MUC6* (Gland mucosal cells, GMCs); *OLFM4* and *REG1A* (Mucosal stem cells, MSCs); *FABP1* and *FABP2* (Enterocytes); *PGA3*, *PGA4* and *LIPF* (Chief Cells) and *ATP4A*, *APT4B* and *VEGFB* (Parietal cells). In addition, gastric cancer (GC) genes were assessed and derived from GEPIA [98]. Markers were visualized utilizing Seurat's "DotPlot" function. The same marker genes were used to annotate cell types in the Sathe et al. dataset. It should be noted that PMCs and GMCs are transcriptionally highly similar and only discriminated by the expression of *MUC5AC* and *MUC6* respectively. Hence, both the unsupervised and supervised transcriptional analyses point to the general mucosal-ness of cells prevalent across donors.

**Removal of batch effects using COMBAT**

To facilitate comparison of the gene expression profiles of HGOs relative to gastric tumor and normal tissue, batch correction was performed using the "COMBAT" function from the surrogate variable analysis package (sva, https://bioconductor.org/packages/release/bioc/html/sva.html) was used. The count matrix stored in the Seurat object from the *Early*, *Mid* and *Late* organoid data and from the Sathe et al. dataset were used as input [41]. Cells were labelled as either belonging to organoid or tissue, which was treated by COMBAT as two individual batches. Upon COMBAT normalization, the adjusted count matrix was utilized as input for the LSI projection.

**Latent semantic indexing projection for Early, Mid and Late scRNA-seq organoid data**

The scRNA-seq data from the *Early*, *Mid* and *Late* organoids were next analyzed to assess progression towards a more malignant, gastric cancer-like phenotype. We deployed a recently published approach based on natural language processing, termed latent semantic indexing (LSI) [99], to compare the organoid scRNA-seq data with the Sathe et al. gastric tumor and normal reference scRNA-seq dataset. The use of a reference enables comparisons across experimental conditions without requiring the definition of discrete cell types in the organoid dataset [100]. First iterative LSI was performed to reduce the dimensionality of the reference dataset. To compare the transcriptional state of the *Early*, *Mid* and *Late* organoids to the gastric tissue dataset, the count matrix of the Sathe et al. reference dataset was subset to the genes in common with the COMBAT corrected organoid matrix. Next, transcript counts were normalized with $\log_2$(counts per 10,000 transcripts +1) and used as input to the LSI projection workflow as detailed below.

In brief, for each dataset, the term frequency and inverse document frequency (TF-IDF) was calculated using the 3000 HVGs generating an essentially normalized count matrix as described by Granja et al. [99]. Singular value decomposition (SVD) was performed, keeping the first 25 dimensions as input for Seurat's shared nearest neighbor (SNN) approach [101]. Cells belonging to the individual clusters were summed up and log CPM-transformed using edgeR's "cpm" function. Across these clusters, the top 3,000 HVGs were identified again. Next, these HVGs were used to perform the TF-IDF, resulting in a new TF-IDF matrix which was used as input for the SVD using the first 25 dimensions. The re-calculated SVD matrix was in turn used as input for Seurat's SNN function with an increased resolution of 0.6. Again, the individual clusters were summed up and the logCPM transformation was calculated using edgeR. This procedure was repeated for one more iteration using 3,000 HVGs and a resolution of 1. The final features and clusters were saved and used as input to the uwot implementation of UMAP, equivalent to the parameters used by Granja et al. [99]. The UMAP embedding was plotted using the ggplot2 package in R and saved for the projection using uwot's "save_uwot" function. Following this, the TF-IDF normalized matrix resulting from the last iteration was added to the original Seurat object using "CreateAssayObject " where it was stored as "LSI" assay. In addition, the SVD matrix as well as the UMAP coordinates were added by executing the "CreateDimReducObject" function. The adapted Seurat object was saved for further processing and used to evaluate the transcriptional similarity of HGO cells.

**Classification of the K-Nearest Neighbor (KNN) environment**

To evaluate whether and which cell populations shift over time, the iterative LSI approach was implemented [99]. For this purpose, the gastric tissue data processed as detailed above was clustered individually with each organoid culture according to the respective time point of the scRNA-seq experiment in *Early*, *Mid* and *Late* cells. Similar to the approach described by Granja et al., [99] the clustering analysis was implemented using 25 dimensions, 1,000 HVGs and Seurat's SNN resolution of 0.2, 0.8 and 0.8 relying on the R package irlba (https://cran.r-project.org/web/packages/irlba/index.html). UMAP visualizations were plotted using ggplot2, where organoid cells were colored according to the time point of interest. In order to illustrate the local neighborhood, a 2D density distribution was added to the UMAP embedding generated before. Next, the location of the individual organoid cells in the UMAP manifold, the local neighborhood of the cells of interest, was assessed. Therefore, the "get.knnx" function from the FNN package in R was used to identify the 25 nearest neighbors (NN) of each organoid cell on the basis of Euclidean distance between the projected organoid cell and the gastric tissue cells in SVD feature space. Through the evaluation of the respective cell identity of the 25 NN and based on the annotations described above, the frequency of each cell type within each organoid culture's neighborhood was determined. After implementation of the described approach, cell type frequencies in the local neighborhood for each individual organoid culture and time point were compared. Note that cell types with minor contributions to the results are not shown in **Fig. 4e**. For all samples, the final frequencies were calculated per culture and plotted using a custom R script.

**Corroboration of inferred single cell copy number states via scDNA-seq**

scDNA sequencing was performed on a representative sample for comparison with CNA states inferred from the scRNA-seq data. The raw base call files from the 10X Chromium sequencer were processed using the Cell Ranger DNA Software Suite (release v1.0, https://support.10xgenomics.com/single-cell-dna). The "cellranger-dna mkfastq" command was used to demultiplex the sequencing samples and to convert barcode and read data to fastq files. This command provides a wrapper around Illumina's "bcl2fastq". Building up on the fastq files, the "cellranger-dna cnv" command was utilized to perform reference alignment, cell calling as well as copy number estimation and hierarchical clustering. Reads were aligned to the hg38 reference genome using a pre-built annotation package downloaded from the 10X Genomics website. Several output files including a barcoded bam file, a summary csv file, containing sample metrics and per-cell summary metrics, as well as a browser extensible data (bed) file were provided. Then, the "node_cnv_calls.bed" file, containing genome-wide coordinates of copy number segments for every single cell, was used as input to a custom R script. This file contained individually sized segments per cell and chromosome. The "bedtools intersect" (v.2.27.1) command was used to construct a bin x cell matrix for all cells which passed the Cellranger DNA QC, facilitating comparisons between cells. The bedfile was then used subset to an individual barcode of interest, sorted according to the bin coordinates. Using "bedtools intersect -wb", the first cell was compared pairwise against a second cell, splitting the genomic coordinates of cell A by their intersection with all other cells, resulting in a bed file with the smallest possible number of overlapping regions across all cells. The bedfile was utilized to create a matrix with bins x cells, which was used as input for a custom R script to create a heatmap representing the copy number values per bin across the genome.

**Inference of copy number from single-cell RNA-seq with inferCNV**

CNAs were inferred from scRNA-seq data using *inferCNV* [102] as previously described (https://github.com/broadinstitute/inferCNV/wiki) [103]. The count matrix from the final quality-controlled Seurat objects generated for each single sample was used as input, with the WT cells from the matching donor used as a reference cell population. *InferCNV* filters low quality genes, normalizes for sequencing depth and applies a log-transformation. Data from each cell was smoothed using a weighted running mean of 151 genes with a pyramidal weighting scheme, favoring genes according to their proximity and centered according to the cell's median expression. CNAs were estimated based on the average normalized gene expression of the cells of interest relative to the normalized gene expression of the reference cells, referring to a subtraction of the mean of the normal from the tumor cells. Dynamic noise filtering was also applied to enhance the signal-to-noise ratio between potential CN altered regions and diploid regions. A six-state hidden Markov model (HMM) was utilized to segment the expression profiles into genomic regions and to predict underlying CN states for each region. Lastly, based on the predicted CN state, a Bayesian latent mixture (BayesNet) model was used to assign posterior probabilities for a segment conforming to a diploid CN state 3. Using the default threshold of $p_{normal} = 0.5$, segments that had an assigned probability of $p_{normal} > 0.5$ were set to CN state 3, while all others remained in a different CN state instead. For each of the above mentioned steps, including the modified expression, HMM and BayesNet, heatmap visualizations were generated by default. In addition to the dynamic de-noising, median filtering was performed to facilitate visual representation of the modified expression. These data were used as input to a custom plot inspired by HoneyBADGER [104].

**Inferring copy number clones from inferCNV output**

In order to assess the copy number landscape of subclones present in the scRNA-seq data, the posterior probability filtered HMM output from *inferCNV* (genes x cells matrix with CN states represented by CN State 1 = 0, CN State 2 = 0.5, CN State 3 = 1 (diploid), CN State 4 = 1.5, CN State 5 = 2, CN State 6 = 2.5) was utilized to define distinct copy number clones. Chromosome arm-level coordinates for hg38 were used and transformed into a *GRange* object. Using the gene names provided in the HMM output matrix, gene coordinates were retrieved with the *biomaRt* package in R and the genomic coordinates were subsequently converted to a *GRange* object. Next, the "mergeByOverlaps" function of the GenomicRanges package was applied to map the respective gene to the matching chromosome arm. To transform the CN state data from gene- to arm-level summaries, the percentage of each chromosome arm covered by genes present in the matrix was calculated (corresponding to informative regions within the chromosome arm).

Next, a gene-specific weighting factor was computed based on gene lengths relative to the length of the coding region per chromosome arm. By multiplying the weighting factor by the respective CN state for each gene, a weighted CN state per gene, per cell was generated. The standard deviation of this metric across all cells was used to assign a single copy number state per arm, per cell. The maximum copy number in this scenario was set to 4, representing gain of more than one allele. Hence, this approach is not well suited to assess the copy number landscape in samples where high CN states or whole genome doubling (WGD) is expected.
A chromosome arm per cell was assigned a specific copy number according to the following equation:

$$CN_a = \begin{cases} CN_w \leq 1 - \sigma \,|\, 0 \; (Bi-allelic\ Loss) \\ 1 - \sigma < CN_w \leq 1 + \sigma \,|\, 1 \; (Loss) \\ 1 + \sigma < CN_w \leq 2 + \sigma \,|\, 2 \; (Diploid) \\ 2 + \sigma < CN_w \,|\, 3 \; (Gain) \end{cases} \tag{5}$$

Where $CN_a$ is the assigned copy number, $CN_w$ represents the weighted copy number state and $\sigma$ corresponds to the standard deviation. The resulting chromosome arm by cell matrix was transposed, and 1 was added to make the copy number states (single copy loss, diploid, single copy gain or gain of more than one copy) more intuitive. Based on this matrix, copy number clones were assigned by investigating the abundance of distinct CNAs per cell. Following this procedure, calls were subject to stringent quality control, whereby CNAs that were not present in specific subclones or were not evident in bulk WGS were considered noise and ignored. Clones were removed if less than 10 cells were assigned to a specific CNA profile. Output files including a data frame of cell barcodes, CNA profiles and assigned clone identifiers were used to perform differential expression analysis between copy number subclones.

**Identification of ECB copy number subclones**
Frequently, ECB defined subclones consisted of two or more copy number subclones. To identify cells belonging to such copy number subclones, the copy number profiles derived from *inferCNV* were assessed for each barcode. In cases where the winning subclone or a subclone comprising most of the population contained more than one copy number clone, it was split into multiple subclones. This was a frequent scenario for winning subclones which accrued additional CNAs. For example, the winning red subclone for D2C2 replicate 2 was split into two subpopulations, (0a and 0b) where 0a harbored a copy number clone with loss of chr4q and gain on chr20q, whereas 0b did not.

**Analysis of differential gene expression and Gene Set Enrichment Analysis (GSEA)**
Differential gene expression was performed using the *FindMarkers* function from Seurat with no log fold threshold. Statistical analysis was based on the Wilcoxon rank sum test, and Bonferroni adjusted p-values are reported. For both volcano plots and GSEA, the full gene list comparing differentially expressed genes between two conditions was used. *Late* and *Mid* cultures were compared to their *Early* counterparts. For ECB defined subclones (or ECB copy number subclones), each subclone was compared to all other cells at a specific time point. The generated gene list was combined with the log2 fold change values and utilized as input for gene set enrichment analysis (GSEA) as implemented in the R package *HTSanalyzeR2* (https://github.com/CityUHK-CompBio/HTSanalyzeR2). GSEA was performed using the MSigDB hallmark gene sets provided by Liberzon and colleagues [105].

#### Summary of software tools

| Name/Version | Source | Link/Identifier |
|---|---|---|
| R (v4.0.1) | - | https://www.r-project.org/ |
| RStudio Server (v1.3.1056) | - | https://login.scg.stanford.edu/ |
| BWA (v0.7.8 & v0.7.17) | Li and Durbin, 2009, Bioinformatics | https://academic.oup.com/bioinformatics/article/25/14/1754/225615 |
| Samtools (v1.8) | Li et al., 2009, Bioinformatics | https://academic.oup.com/bioinformatics/article/25/16/2078/204688 |
| QDNAseq (v1.22.0) | Scheinin et al., 2014, Genome Research | https://bioconductor.org/packages/release/bioc/html/QDNAseq.html |
| GATK (v4.1.4.1 & v4.1.7.0) | | https://gatk.broadinstitute.org/hc/en-us |
| Strelka (v2.9.10) | Kim et al., 2018, Nature Methods | https://www.nature.com/articles/s41592-018-0051-x |
| PURPLE (v3.0), AMBER (v3.3), COBALT (v1.8) | Cameron et al., 2019, bioRxiv | https://www.biorxiv.org/content/10.1101/781013v1 |
| Manta (v1.6.0) | Chen et al., 2015, Bioinformatics | http://dx.doi.org/10.1093/bioinformatics/btv710 |
| Delly (v0.8.1) | Rausch et al., 2012, Bioinformatics | https://academic.oup.com/bioinformatics/article/28/18/i333/245403 |

| | | |
|---|---|---|
| GRIDSS (v2.9.4) | Cameron et al., 2019, bioRxiv | https://www.biorxiv.org/content/10.1101/781013v1 |
| SvABA (v.1.1.3) | Wala et al., 2018, Genome Research | https://genome.cshlp.org/content/28/4/581 |
| JaBbA | Hadi et al. 2020, Cell | https://github.com/mskilab/JaBbA |
| Cell Ranger DNA Software Suite (v1.0) | - | https://support.10xgenomics.com/single-cell-dna |
| bedtools (v2.27.1) | Quinlan and Hall, 2010, Bioinformatics | https://academic.oup.com/bioinformatics/article/26/6/841/244688 |
| Cell Ranger Single-Cell Software Suite (v3.0) | - | https://support.10xgenomics.com/single-cell-gene-expression |
| Seurat (v.3.2) | Butler et al., 2018, Cell | https://satijalab.org/seurat/ |
| SCtransform (v0.2.1) | Hafemeister and Satija, 2019, Genome Biology | https://github.com/ChristophH/sctransform |
| DoubletFinder (v2.0.3) | McGinnis et al., 2019, Cell Systems | https://github.com/chris-mcginnis-ucsf/DoubletFinder |
| inferCNV (v1.2.1) | Patel et al., 2014, Science | https://github.com/broadinstitute/inferCNV/wiki |
| HTSanalyzeR2 (v0.99.19) | Subramanian et al., 2005, PNAS | https://github.com/CityUHK-CompBio/HTSanalyzeR2 |
| sva (v3.34.0) | Leek et al., 2020, | http://bioconductor.org/packages/release/bioc/html/sva.html |
| iterativeLSI | Granja et al., 2019, Nature Biotechnology | https://github.com/GreenleafLab/MPAL-Single-Cell-2019 |
| irlba (v2.3.3) | - | https://cran.r-project.org/web/packages/irlba/index.html |
| ClusterSV | Li et al., 2020, Nature | https://github.com/cancerit/ClusterSV |

Supplementary Figures 1-19

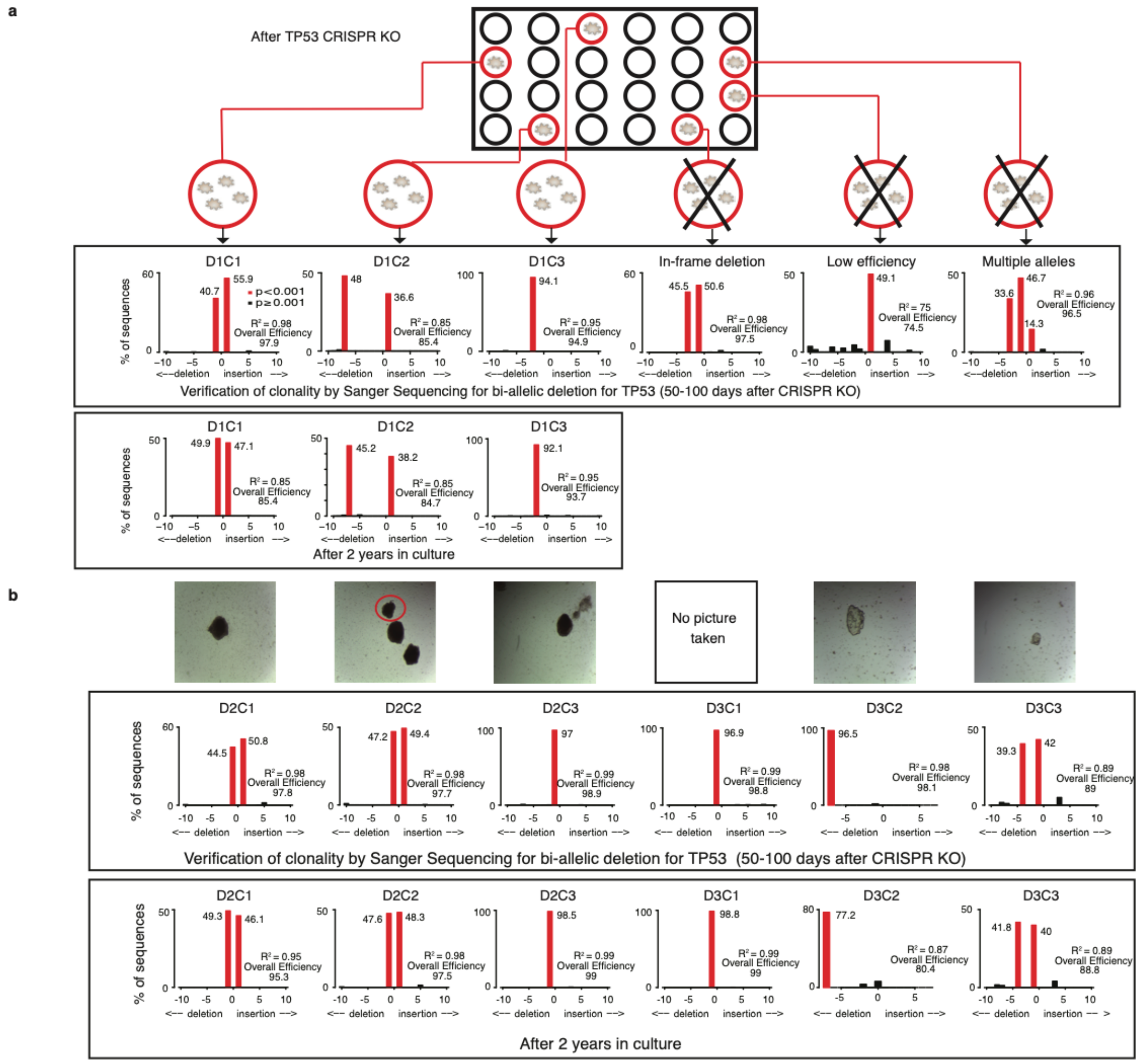

**Supplementary Figure 1. Assessment of single cell derived organoid cultures via Sanger sequencing. a,** Illustration of how single organoids were picked following nutlin-3 selection to establish clonal *TP53*$^{-/-}$ and *APC*$^{-/-}$ human gastric organoids. Individual organoids were Sanger sequenced for the *TP53* deletion site. Only organoid cultures with bi-allelic deletions were retained, while those with in-frame deletions, low efficiency of deletion or more than two alleles (due to multiple knock-out cultures being present) were discarded. The same procedure was utilized to verify clonal status over time, including after two years of culture. Sanger sequencing of the deletion site over time was analyzed using *Tracking of Indels by Decomposition* (TIDE). The $R^2$ value is an assessment of goodness of fit calculated by TIDE. **b,** As in panel a, but for organoids derived from donors 2 and 3.

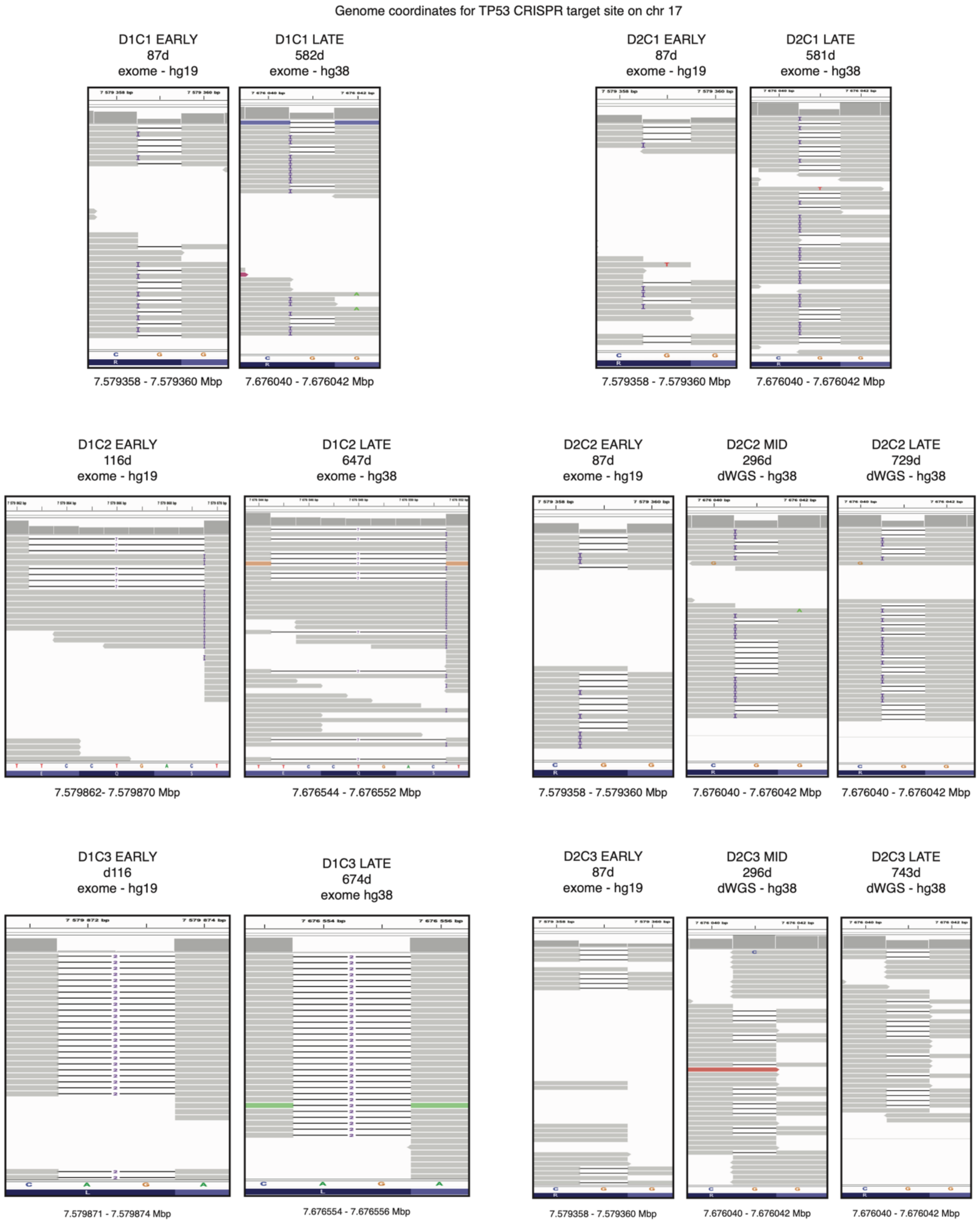

**Supplementary Figure 2**. **Verification of *TP53* edits based on genome sequencing.** Integrated Genome Viewer (IGV) plots of the CRISPR/Cas9 *TP53* edit sites for cultures from donors 1 and 2 over time based on WGS or WES.

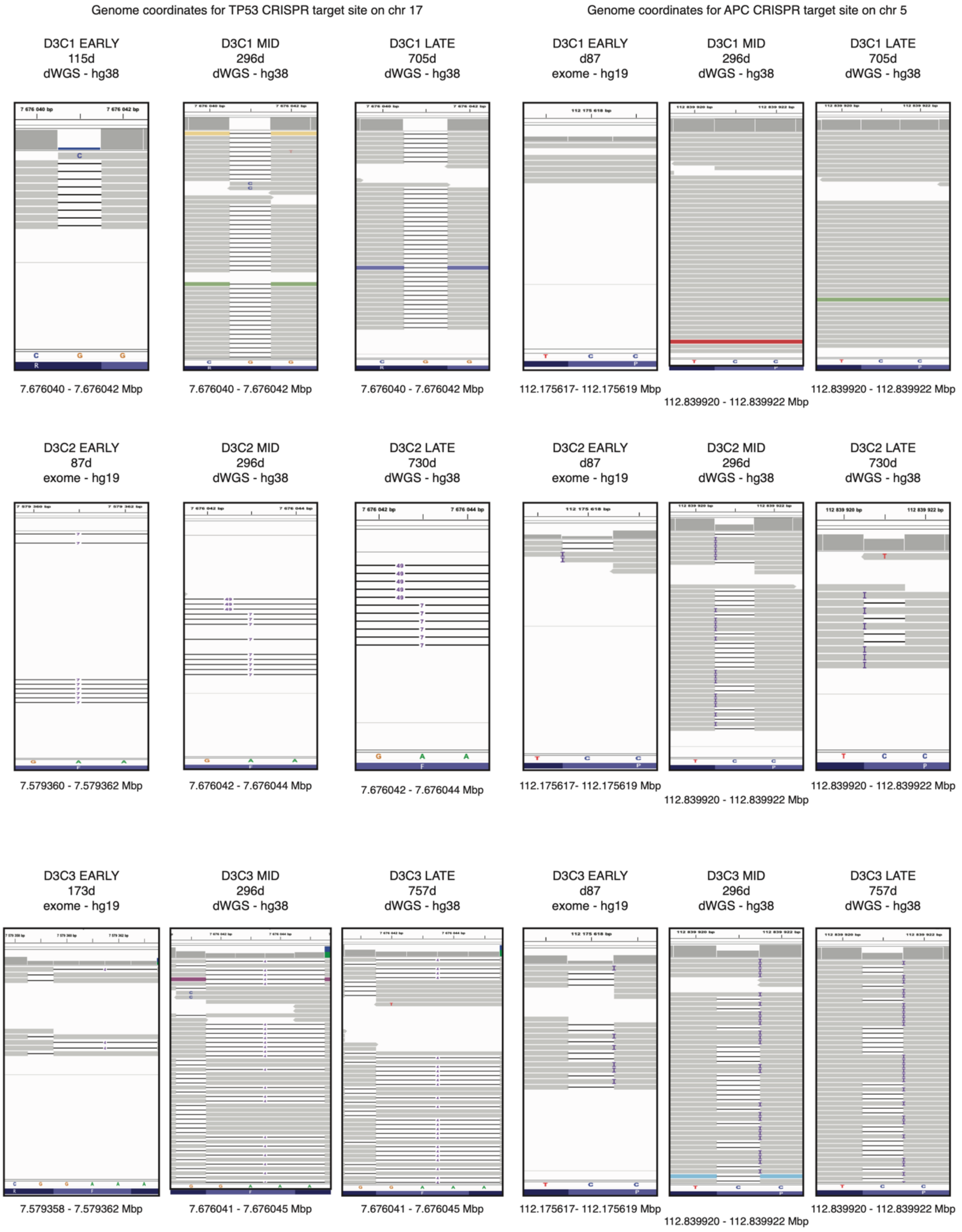

**Supplementary Figure 3**. **Verification of *TP53* and *APC* CRISPR/Cas9 edits based on genome sequencing.** Integrated Genome Viewer (IGV) plots of the CRISPR/Cas9 *TP53* and *APC* edit sites for donor 3 cultures over time based on WGS or WES.

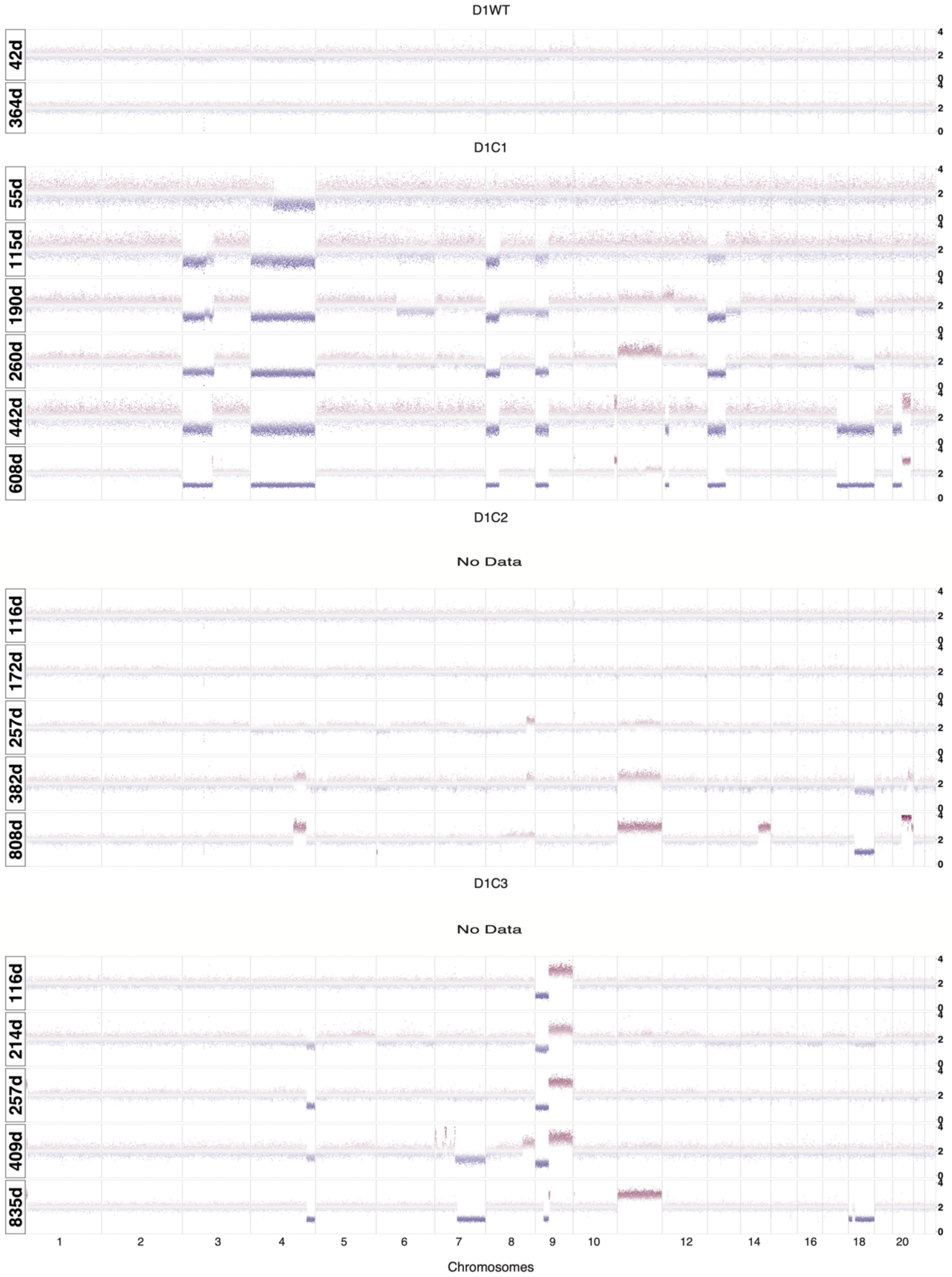

**Supplementary Figure 4. Copy number aberration profiles in Donor 1.** Copy number alteration (CNA) profiles based on shallow whole genome sequencing (sWGS) of WT human gastric organoid cultures (passage 2) and *TP53* deficient cultures. The Y-axis denotes integer copy number based on normalized read counts across 50kb windows of the genome.

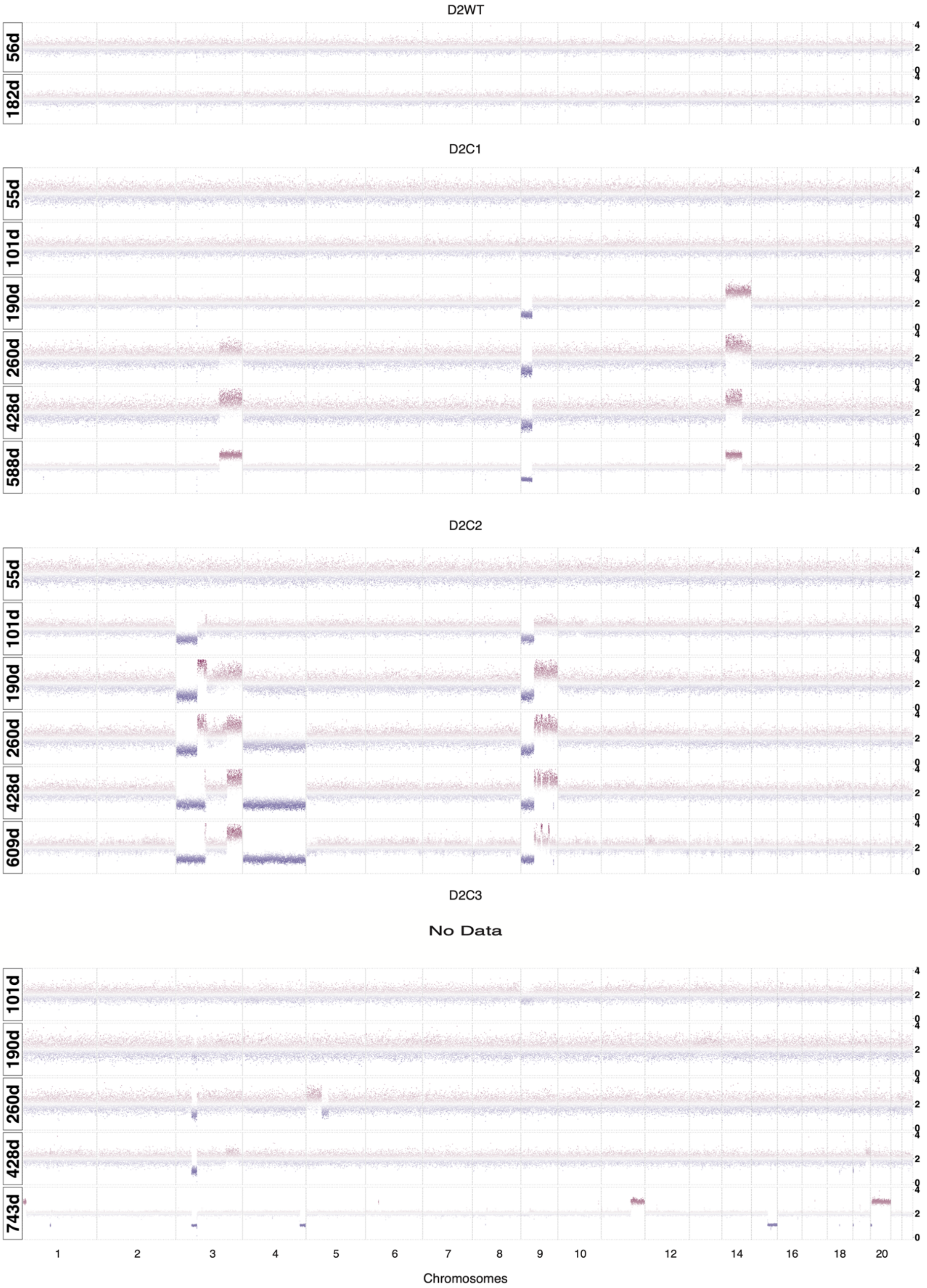

**Supplementary Figure 5. Copy number aberration profiles over time in Donor 2.** Copy number alteration (CNA) profiles based on shallow whole genome sequencing (sWGS) of WT human gastric organoid cultures (passage 2) and *TP53* deficient cultures. The Y-axis denotes integer copy number based on normalized read counts across 50kb windows of the genome.

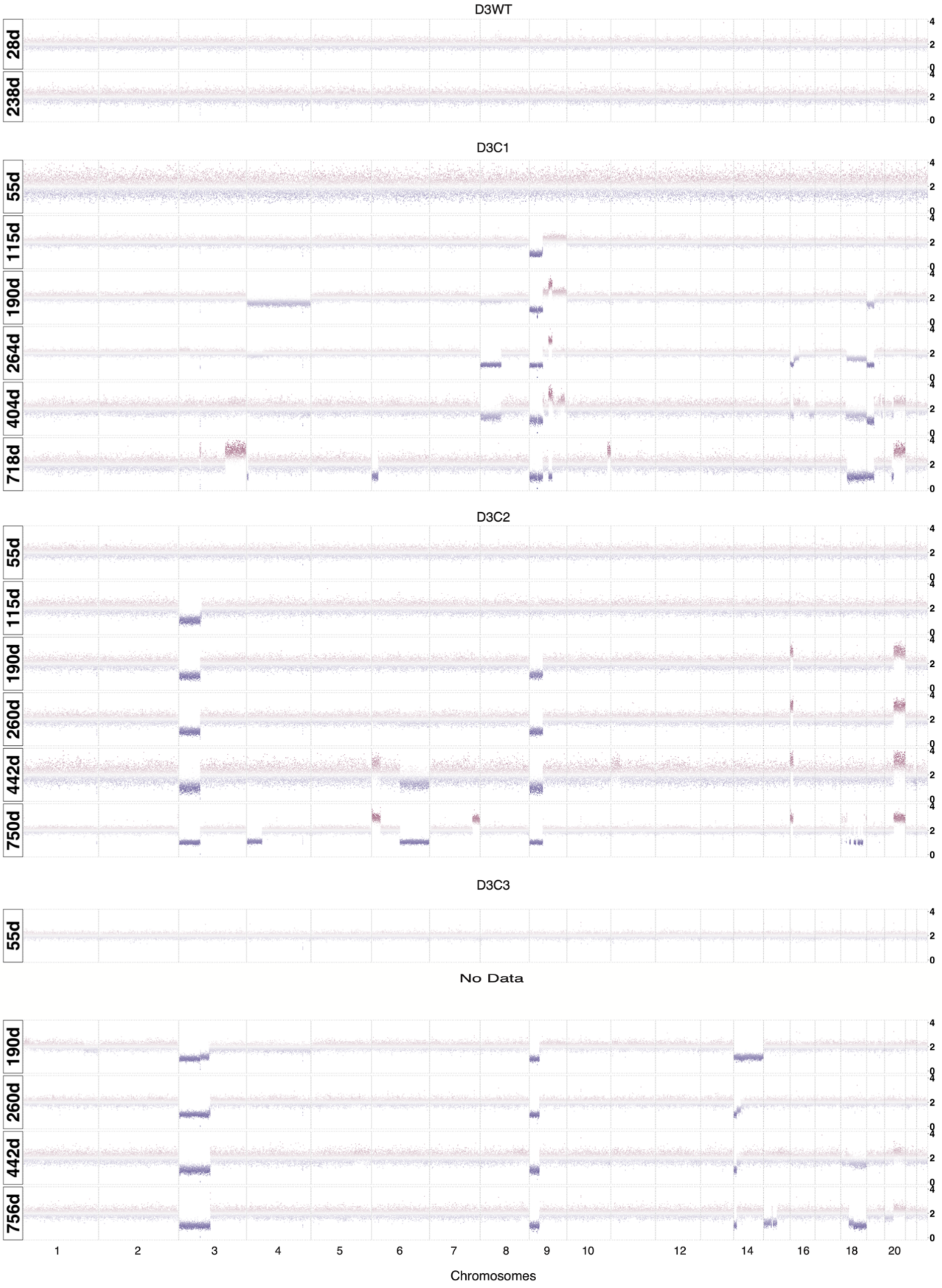

**Supplementary Figure 6. Copy number aberration profiles over time in Donor 3.** Copy number alteration (CNA) profiles based on shallow whole genome sequencing (sWGS) of WT human gastric organoid cultures (passage 2) and *TP53* deficient cultures. The Y-axis denotes integer copy number based on normalized read counts across 50kb windows of the genome.

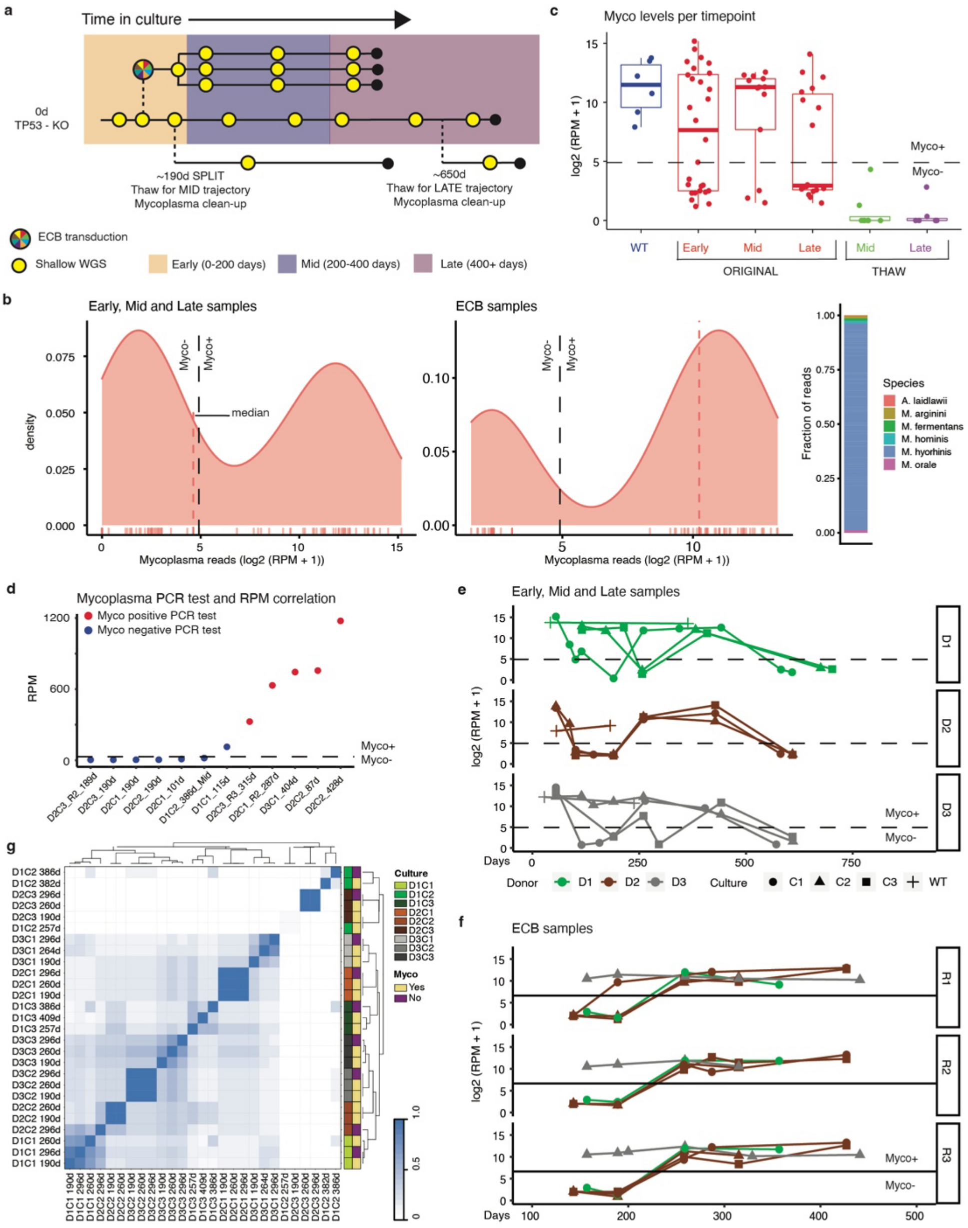

**Supplementary Figure 7. Detection of mycoplasma contamination and evaluation with molecular features. a,** Timeline for a typical culture, where culture time is shown (not real time; Mid and Late trajectories where thawed and cultured in parallel). Note that the Mid and Late thaw cultures were cleaned from mycoplasma at thawing. **b**, Overview of mycoplasma contamination for all samples based on sWGS reads mapped to Mycoplasma genomes. The black dotted line denotes 30 mycoplasma reads per million (RPM) and red dotted line shows the median sample mycoplasma level. The bar-plot on the right shows the fraction of mycoplasma reads per genome. **c**, Mycoplasma contamination stratified on time periods and trajectories. Each dot corresponds to a sequenced sample. The black dotted line denotes 30 mycoplasma RPM. Boxes show inter-quartile range (IQR), center lines represent the median, whiskers extend by 1.5 × IQR. **d**, Correlation between mycoplasma PCR status and mycoplasma RPM from whole genome sequencing. Sample were chosen based on closeness in mycoplasma RPM to the valley shown in (b). The black dotted line denotes 30 RPM. **e,** Mycoplasma levels over time from the longitudinal evolution experiment (i.e., Mid and Late thaw trajectories not included). The black dotted line denotes 30 mycoplasma RPM. **f,** Same figure as in (e), but for the ECB replicates cultures. **g,** To facilitate comparisons, for each of the 9 cultures, 3 samples were chosen for the pairwise Jaccard index analysis based on their proximity in time to the split point (denoted in panel a). The plot shows that samples from the same culture cluster together, regardless of mycoplasma levels. For example, D2C1 at day 190 (the time of the split), day 260 original trajectory and day 296 of the mid thaw trajectory all have chromosome 9p deletion and 14q amplification and no other alterations, resulting in a Jaccard index of 1. The only sample that did not cluster immediately adjacent to the other samples of the same culture was D1C2 at day 190, because it lacked CNAs at that time point. Rather, it clustered with D2C3 at day 190 which similarly lacked CNAs. Samples were clustered using divisive analysis clustering.

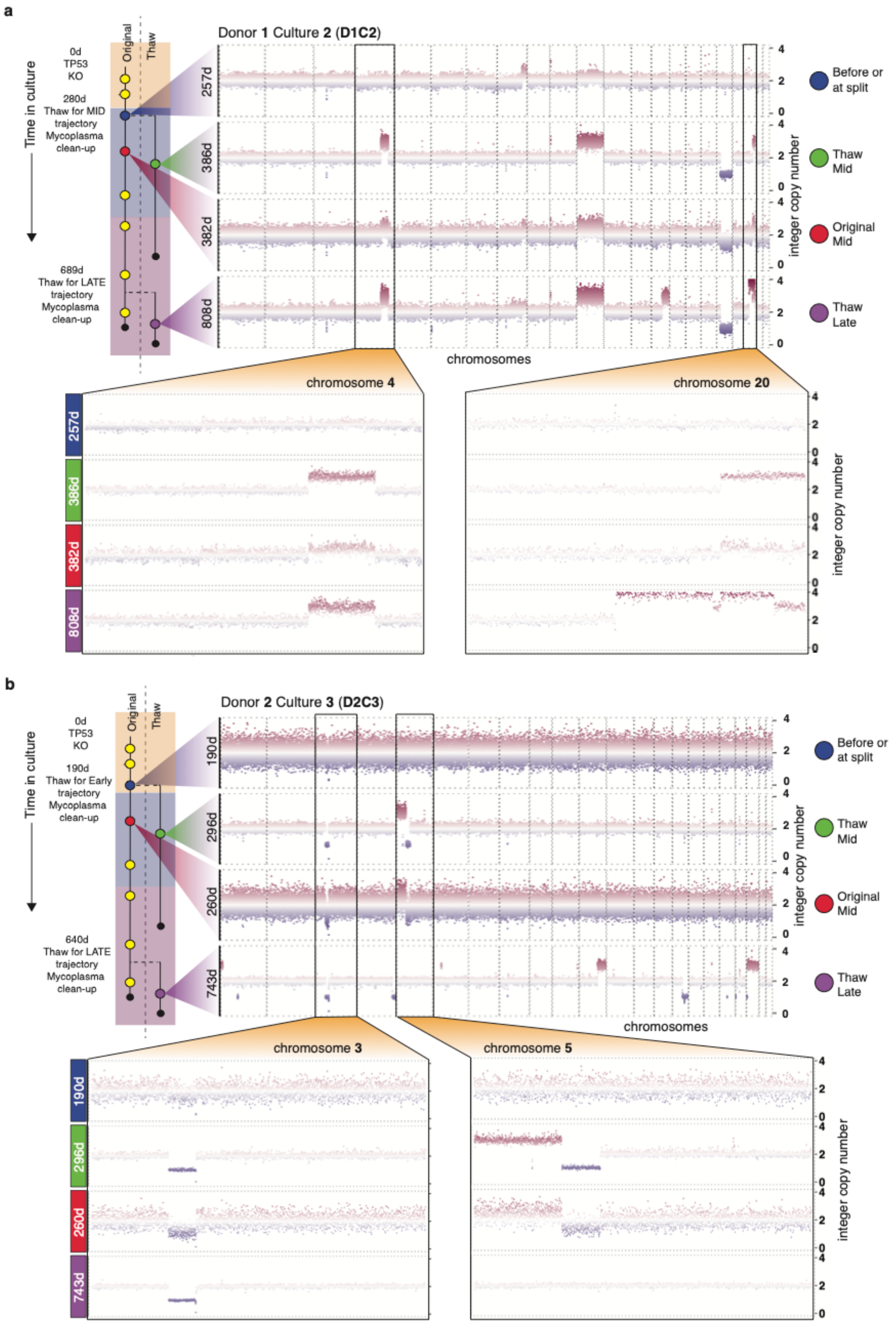

**Supplementary Figure 8. Convergent copy number evolution across cultures, irrespective of mycoplasma levels. a,** Copy number alterations (CNAs) based on shallow whole genome sequencing (sWGS) were assessed at multiple time points along the original trajectory and the 'thawed' myco-free trajectories for D1C2. Culture times for 'Thaw Mid' and 'Original Mid' are similar. The thaw Mid' culture was revived more than a year after the original experiment and was grown separately in culture time for more than 100 days. The boxes zoom in on chromosomes 4 and 20, both of which contain *within chromosome arm* breaks showing *identical breakpoints between* the Original Mid and Thaw Mid cultures, where the latter was myco-negative. **b,** Same as for panel a, but for D2C3.

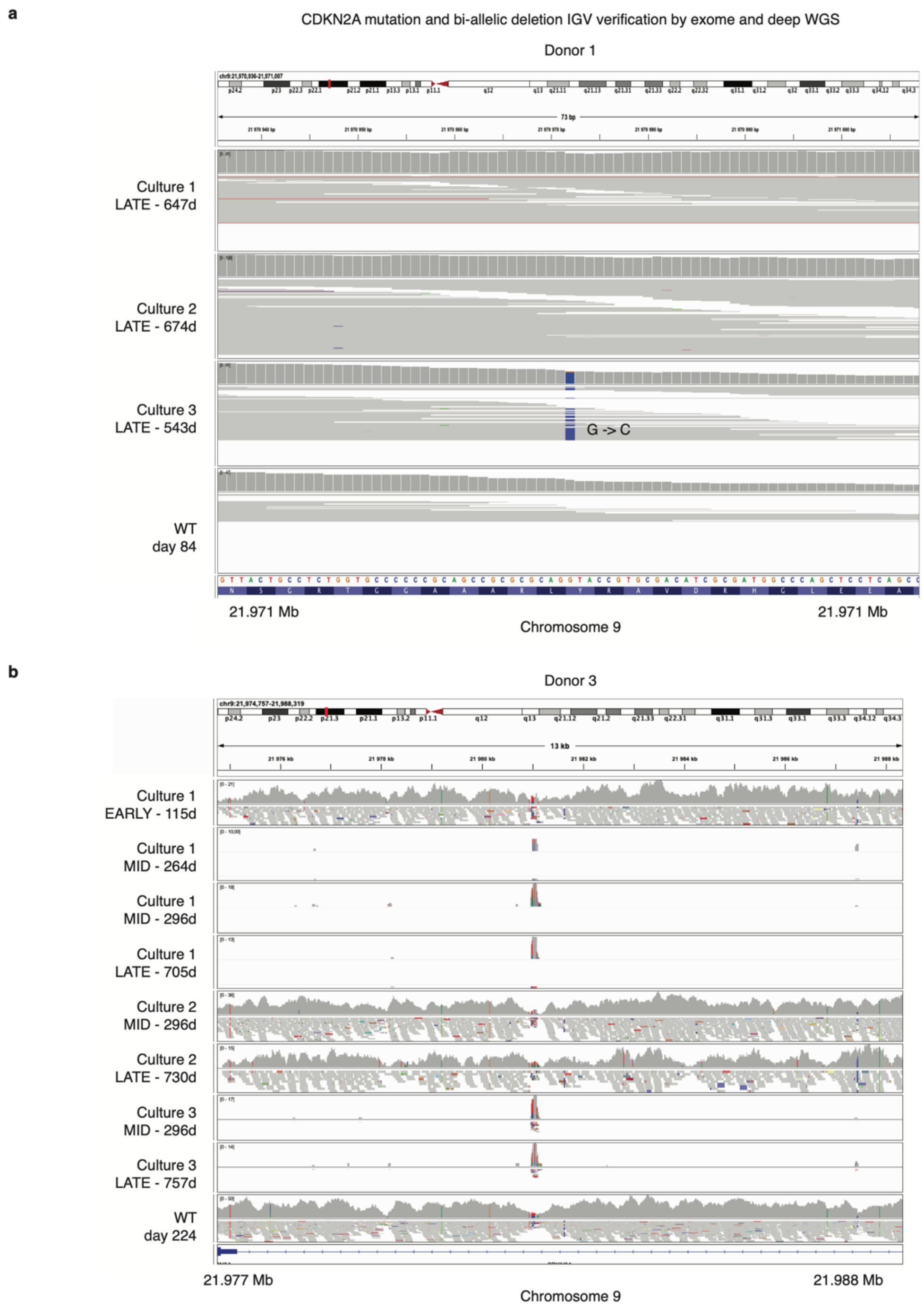

**Supplementary Figure 9. Summary of *CDKN2A* genomic alterations and bi-allelic inactivation.** Integrated Genomic Viewer (IGV) plots illustrating mutations or loss of genomic segments in the *CDKNA2A* gene. **a,** mutation in D1C1 assessed by WES. **b,** loss of genomic segment in D1C1 and D1C3, but not D1C2, assessed by WGS.

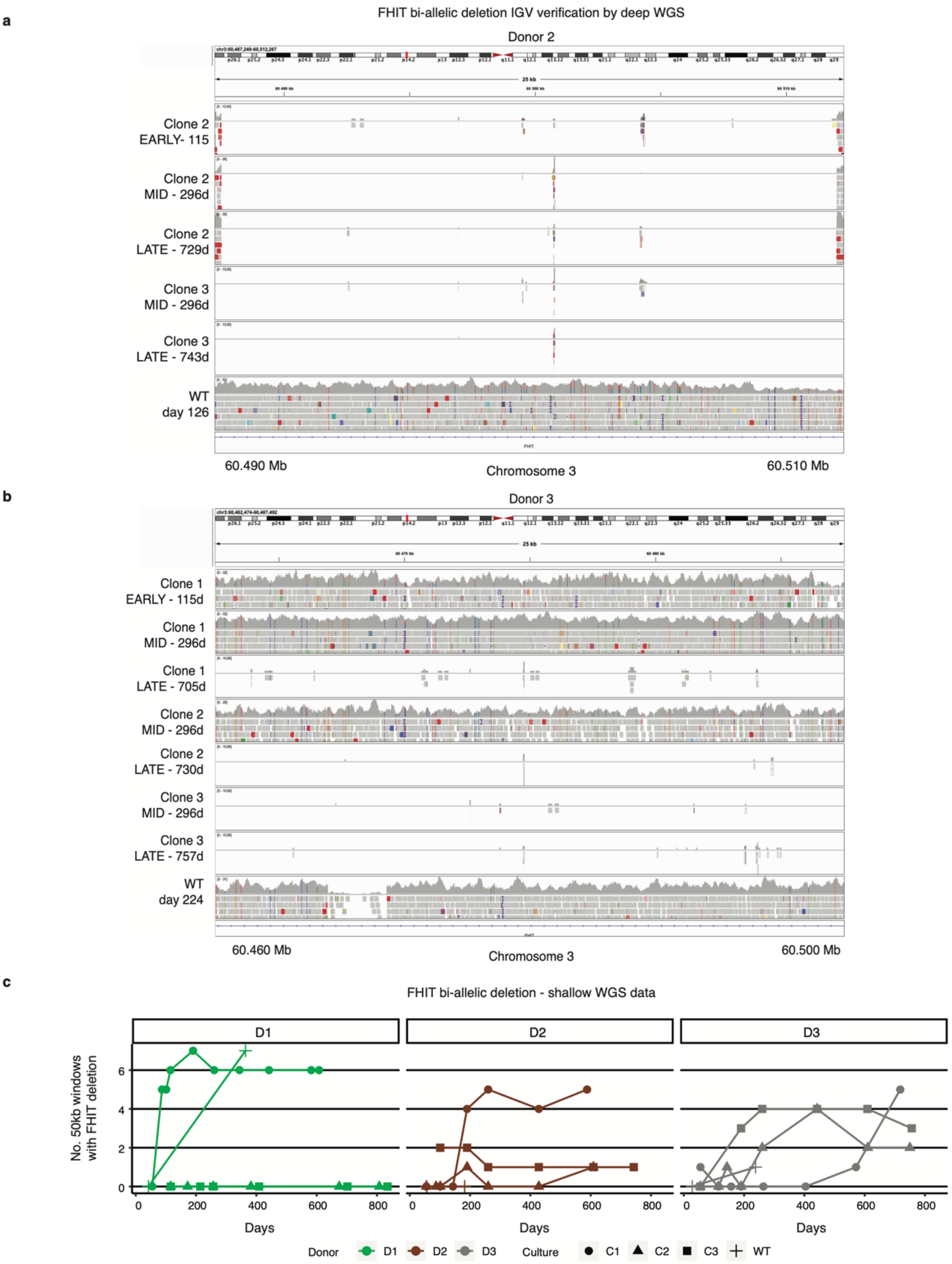

**Supplementary Figure 10. Summary of *FHIT* genomic alterations and bi-allelic inactivation.** **a,** IGV plots illustrating loss of genomic segments in the *FHIT* gene as assessed by whole genome sequencing for donor 2. **b,** IGV plots illustrating loss of genomic segments in the *FHIT* gene as assessed by whole genome sequencing for donor 3. **c,** Loss of genomic segments in the *FHIT* gene over time across clones based on shallow whole genome sequencing (sWGS). Each point represents the number of 50kb windows that have less than 25% of normalized reads compared to the median of all windows, suggestive of a subclone with bi-allelic loss. Note that both D1 and D3 WT cultures exhibit *FHIT* loss at later time points.

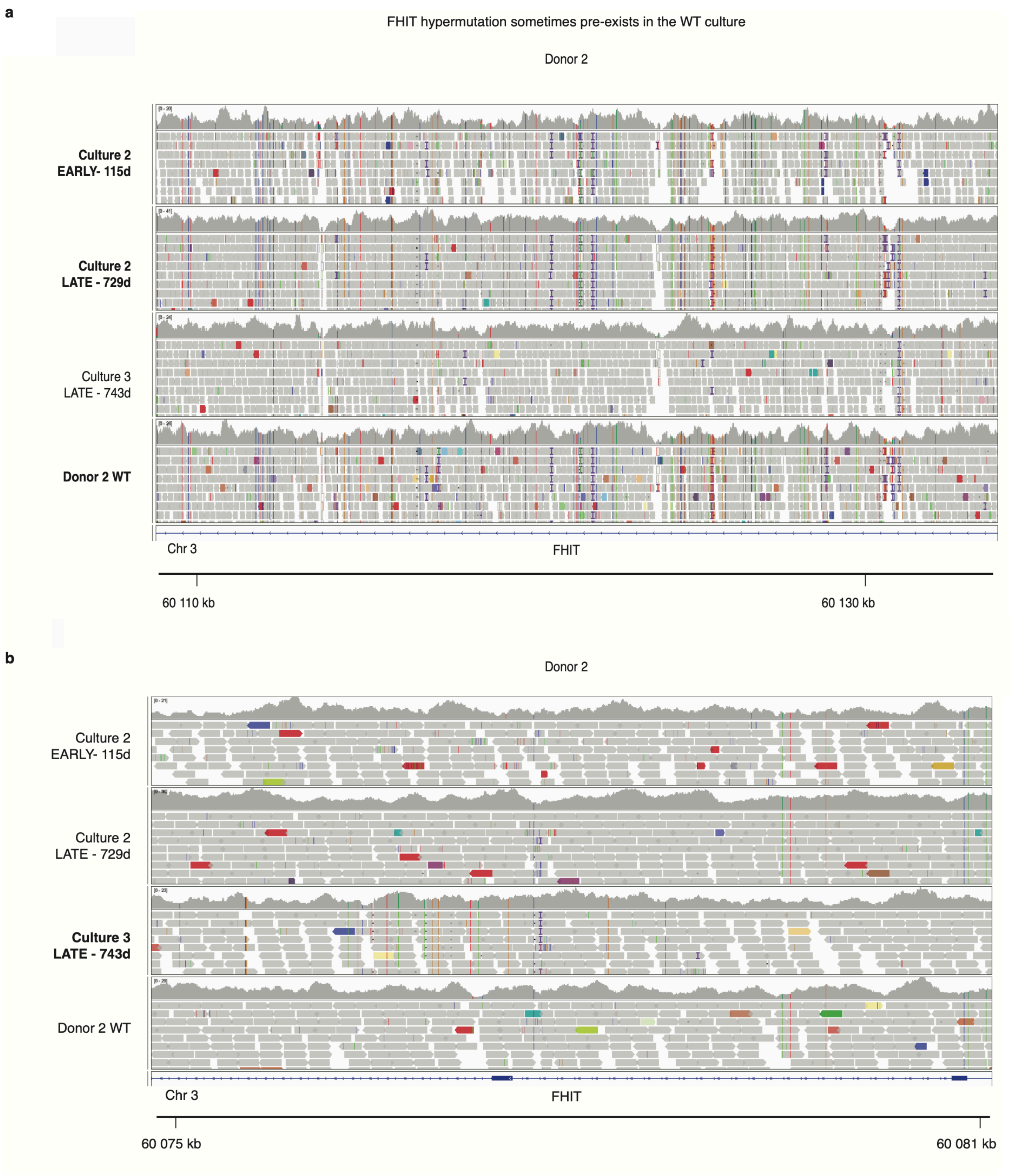

**Supplementary Figure 11. Hypermutation at the *FHIT* locus in gastric organoids from donor 2.** IGV plots illustrating hypermutation in several separate regions of the *FHIT* gene for donors 2 and 3 across all cultures with available WGS data. Sample name in bold indicates that the sample carries a hypermutated region. **a,** shows similar hypermutation patterns in D2C2 and D2WT but not D2C3, potentially due to somatic mosaicism in the WT. **b,** shows another region of *FHIT* with a small hypermutation pattern in D2C3, but not in D2C2 and D2WT.

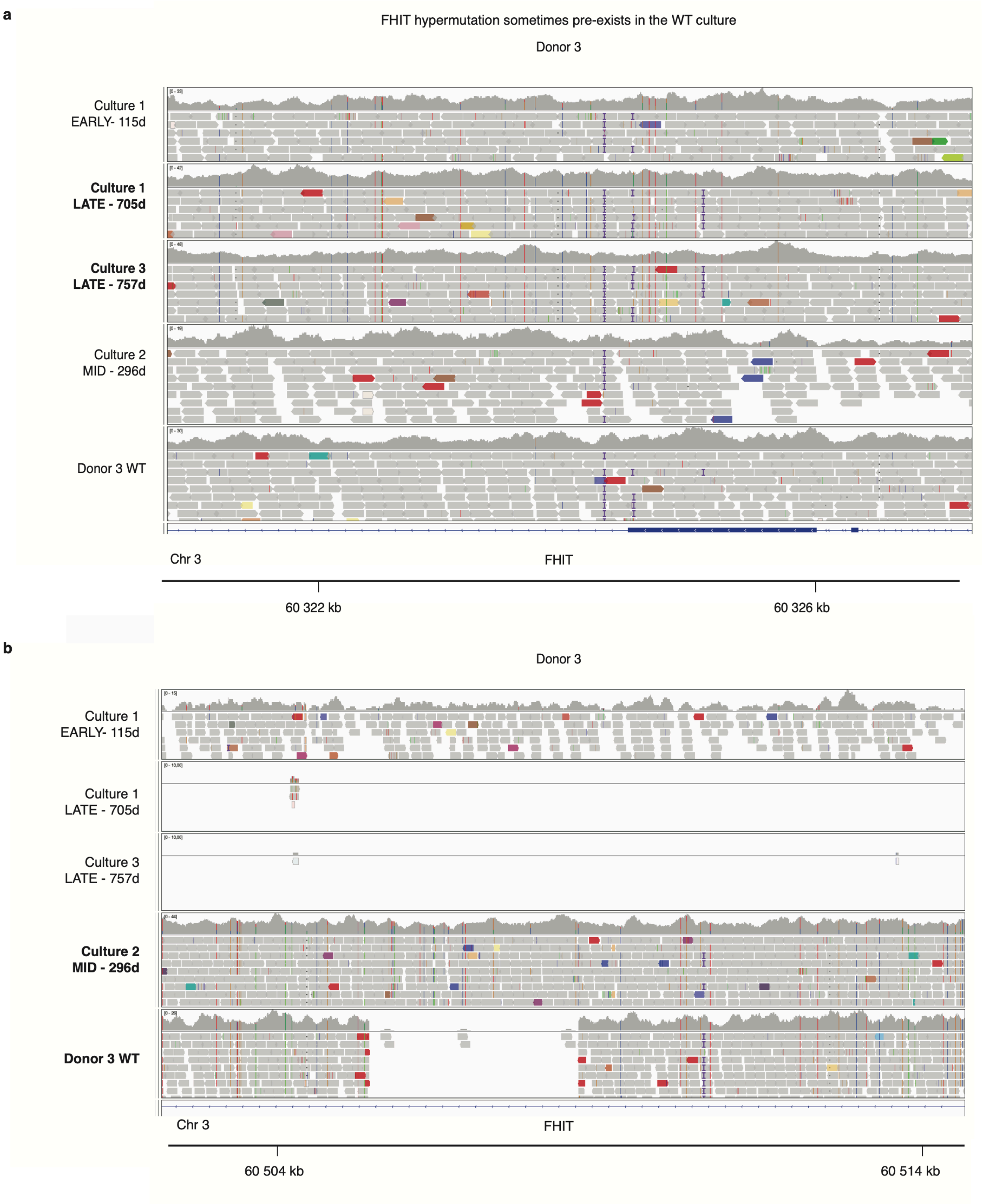

**Supplementary Figure 12. Hypermutation at the *FHIT* locus in gastric organoids from donor 3.** IGV plots illustrating hypermutation in several separate regions of the *FHIT* gene for donor 3 across all cultures with available WGS data. Sample name in bold indicates that the sample carries a hypermutated region. **a,** shows hypermutation in D3C1 and D3C3, but not in D3C2 and D3WT, suggesting mosaic hypermutation patterns in D3WT. **b,** shows similar patterns of hypermutation at *FHIT* in D3C2 and D3WT, but not D3C1. In addition, a small deletion in an intronic region of *FHIT* is evident in D3WT.

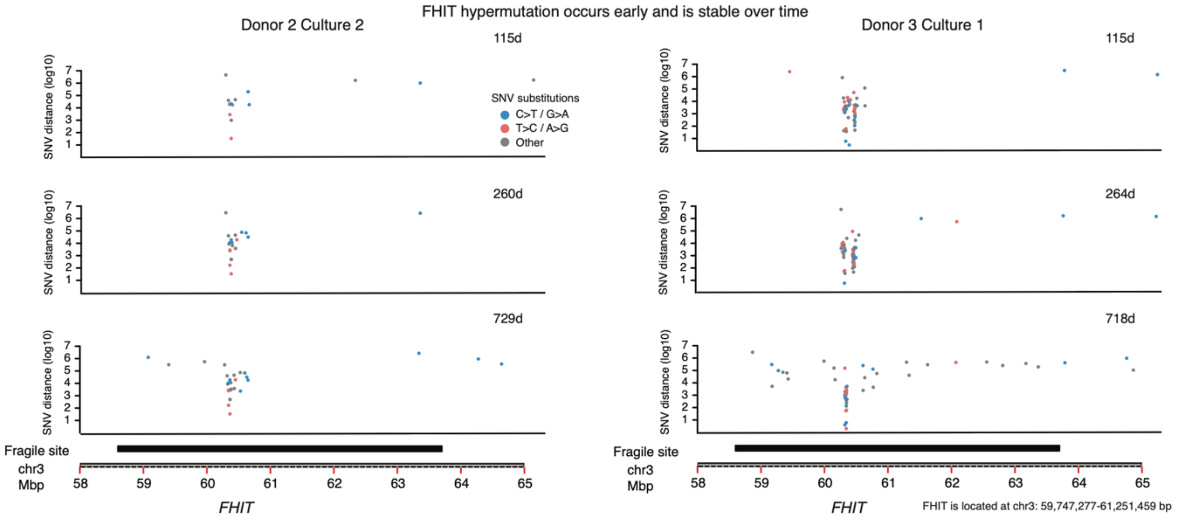

**Supplementary Figure 13. Spatial location of hypermutations at the *FHIT* locus.** Patterns of SNVs at the *FHIT* locus as indicated in the legend. Black horizontal lines denote known fragile sites in the genome.

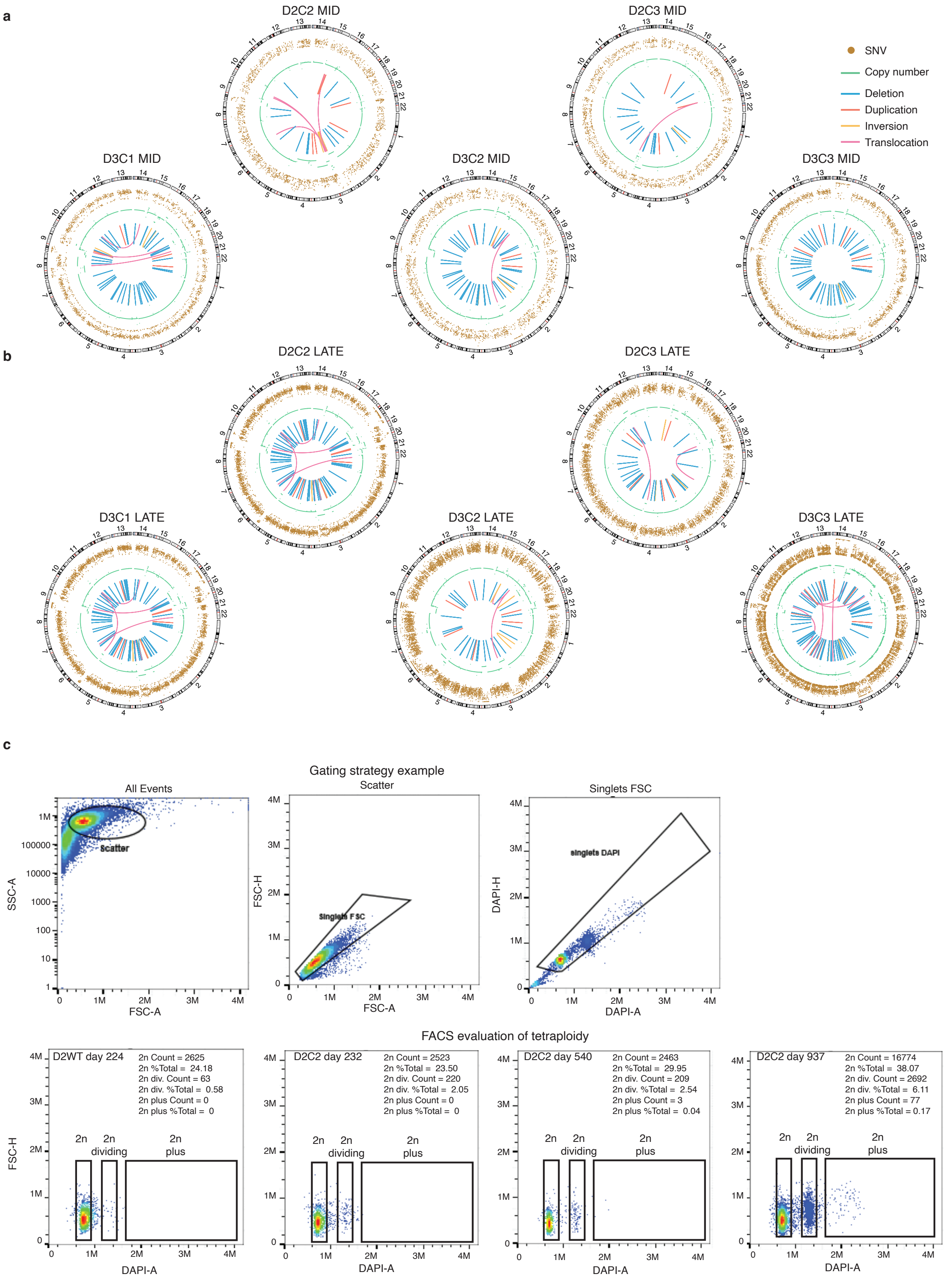

**Supplementary Figure 14. Summary of genomic alterations over time based on WGS of donors 2 and 3 and evaluation of ploidy. a,** Circos plots depicting SNVs (corrected variant allele frequencies, outer track), copy number alterations (CNAs, logR values, middle track) and structural variants (SVs, inner track) based on WGS analysis of D2C2, D2C3, D3C1, D3C2 and D3C3 sampled at Mid. **b,** same as (a) but sampled at Late time point. **c,** FACS sorting on cell cycle synchronized and DAPI stained cells from D2 WT and corresponding *TP53*$^{-/-}$ culture (D2C2). Top panel gating strategy. Bottom panel HGO data. By day 937 a small number of cells in the *TP53*$^{-/-}$ culture were 2n plus indicative of greater genome content than an actively dividing diploid cell.

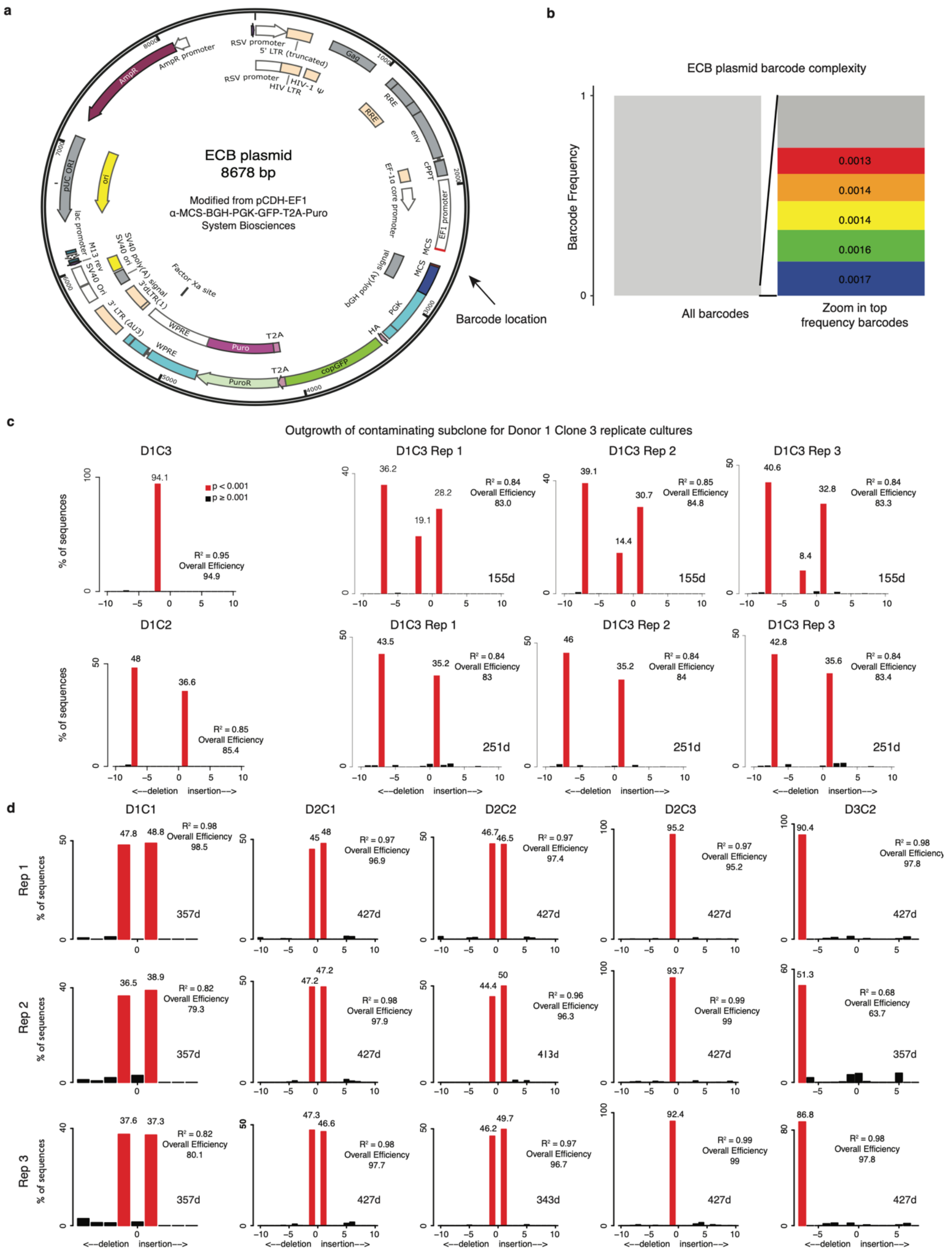

**Supplementary Figure 15. Expressed cellular barcode construct design and quality control assessment. a**, ECB plasmid vector design. **b**, Verification of ECB plasmid complexity. The complexity of the plasmid after insertion of the ECB, but before production of lentiviral particles, was examined by barcode PCR amplification and sequencing. The top barcode constituted 0.0017% of all reads. **c**, Outgrowth of a contaminating subclone. Replicates were monitored for the correct *TP53* knockout site via Sanger sequencing and TIDE analysis. This analysis indicates that the ECB replicates of donor 1 culture 3 (D1C3) were contaminated by a subclone from donor 1 culture 2 (D1C2) resulting in exclusion of replicate cultures from D1C3. **d**, Sanger sequencing of the deletion site over time was analyzed using *Tracking of Indels by Decomposition* (TIDE) and indicates that the allele frequency of the *TP53* edit site is stable over time across replicates. The $R^2$ value is an assessment of goodness of fit calculated by TIDE.

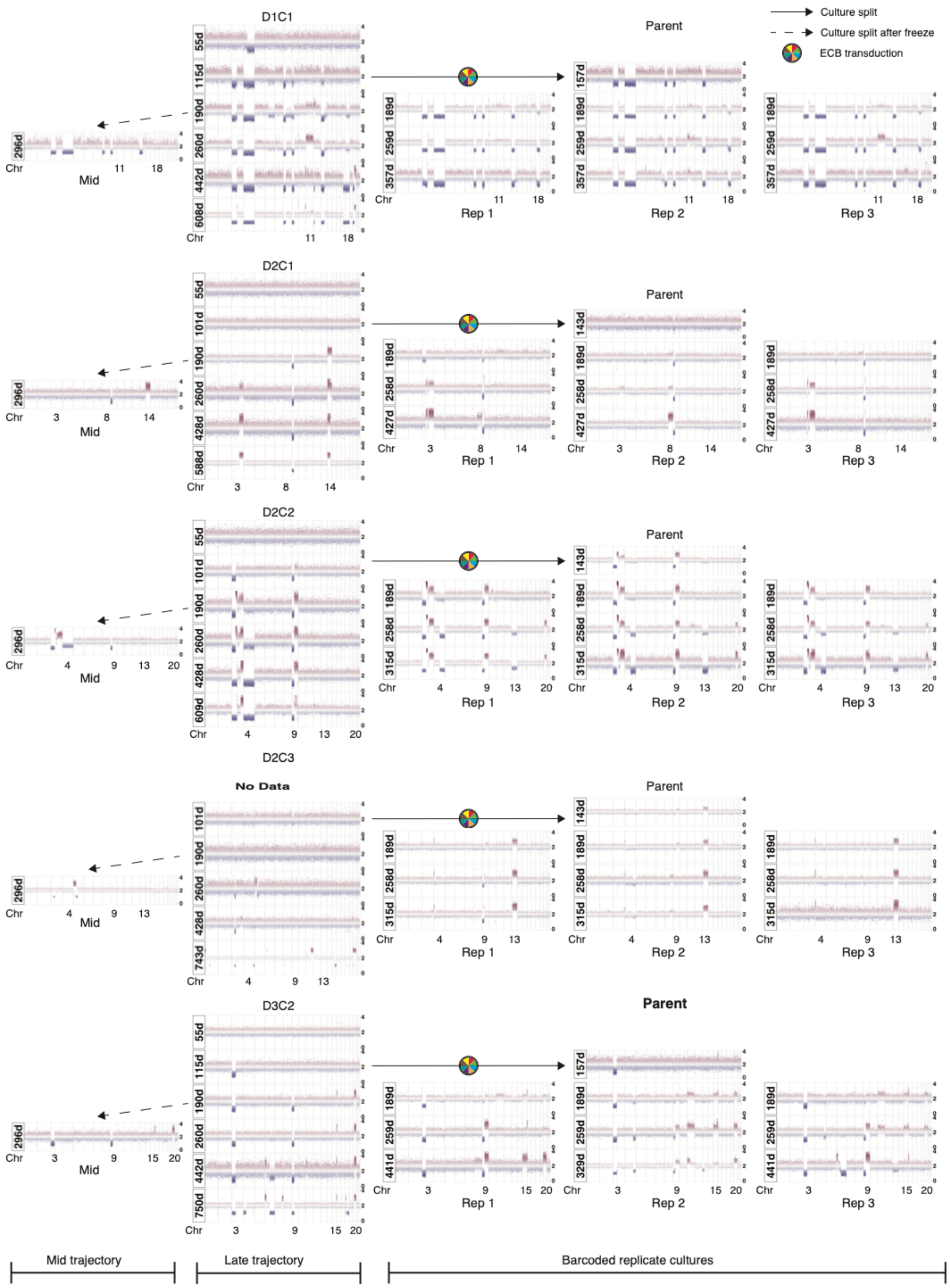

**Supplementary Figure 16. Comparison of the landscape of copy number alteration (CNAs) between non-barcoded and barcoded replicate cultures over time.** The column with donor and culture identifiers shows sWGS data over time for the non-barcoded samples. Solid arrows and multi-colored circles indicate where the population was split to generate the ECB replicate cultures. Similar patterns of CNAs were often observed across non-barcoded versus barcoded cultures, although there were exceptions most notably for D2C3. The greater divergence in CNA profiles for D2C3 may be attributable to the fact that the parental population harbored few detectable CNAs at the time of the split, and as such CNAs develop independently across replicates with fewer constraints.

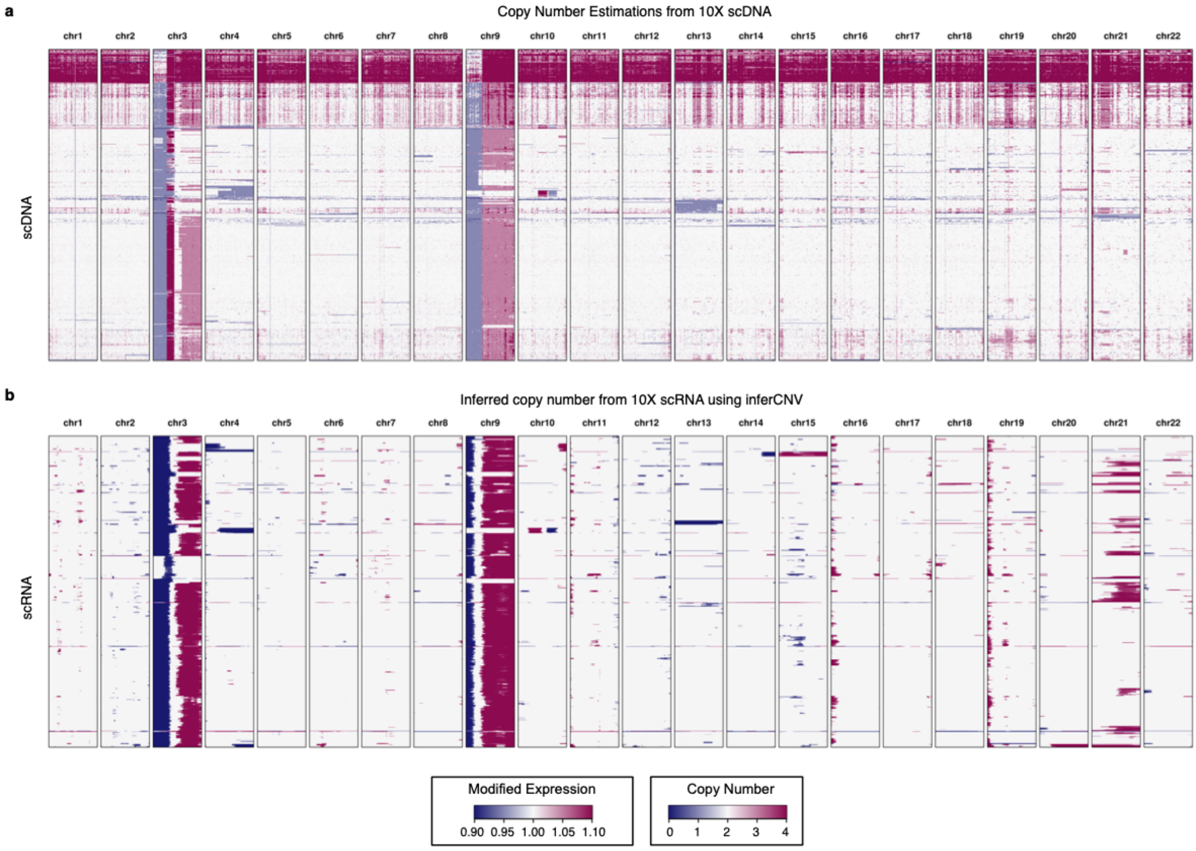

**Supplementary Figure 17. Comparison of single cell DNA sequencing and inferred copy number alteration from single cell RNA sequencing**. a, Copy number estimates based on single cell DNA sequencing on the 10x Chromium platform for D2C2, replicate 2 at day 173. Cells at the top with high copy number (red) are attributable to doublets. b, Inferred copy number alterations represented as modified expression based on inferCNV analysis of single cell RNA-sequencing on the 10X Chromium platform for the same sample corroborates the single cell DNA-sequencing data.

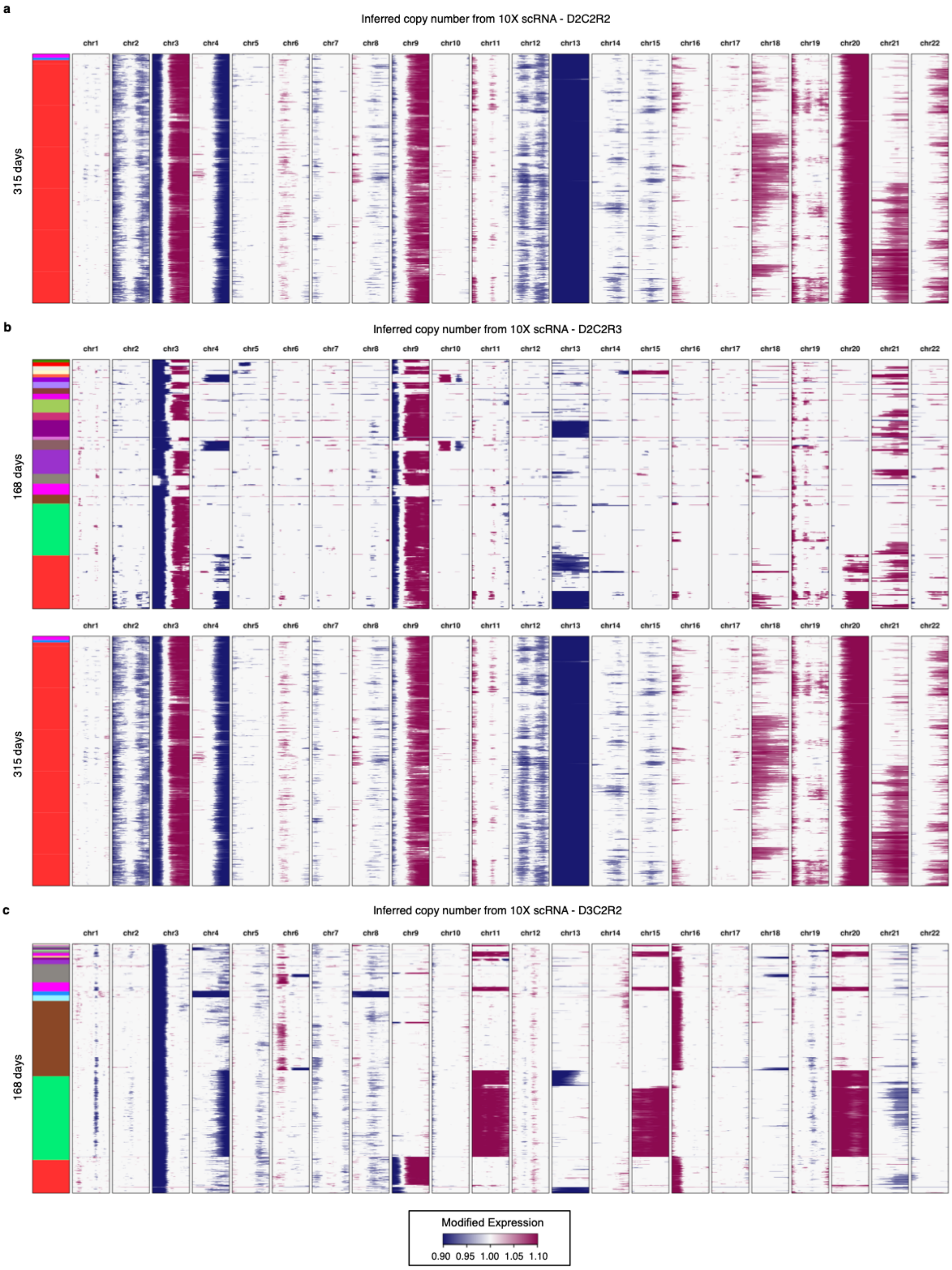

**Supplementary Figure 18. Inference of copy number alterations at single cell resolution from ECB experiments over time. a,** Inference of copy number from 10X single cell RNA sequencing data using inferCNV for Donor 2, Culture 2, Replicate 2 (D2C2R2) at day 315. **b,** As in (A) but for Donor 2, Culture 2, Replicate 3 (D2C2R3) at day 173 and at day 315. **c,** as in (A) but for Donor 3, Culture 2, Replicate 2 (D3C2) at day 173.

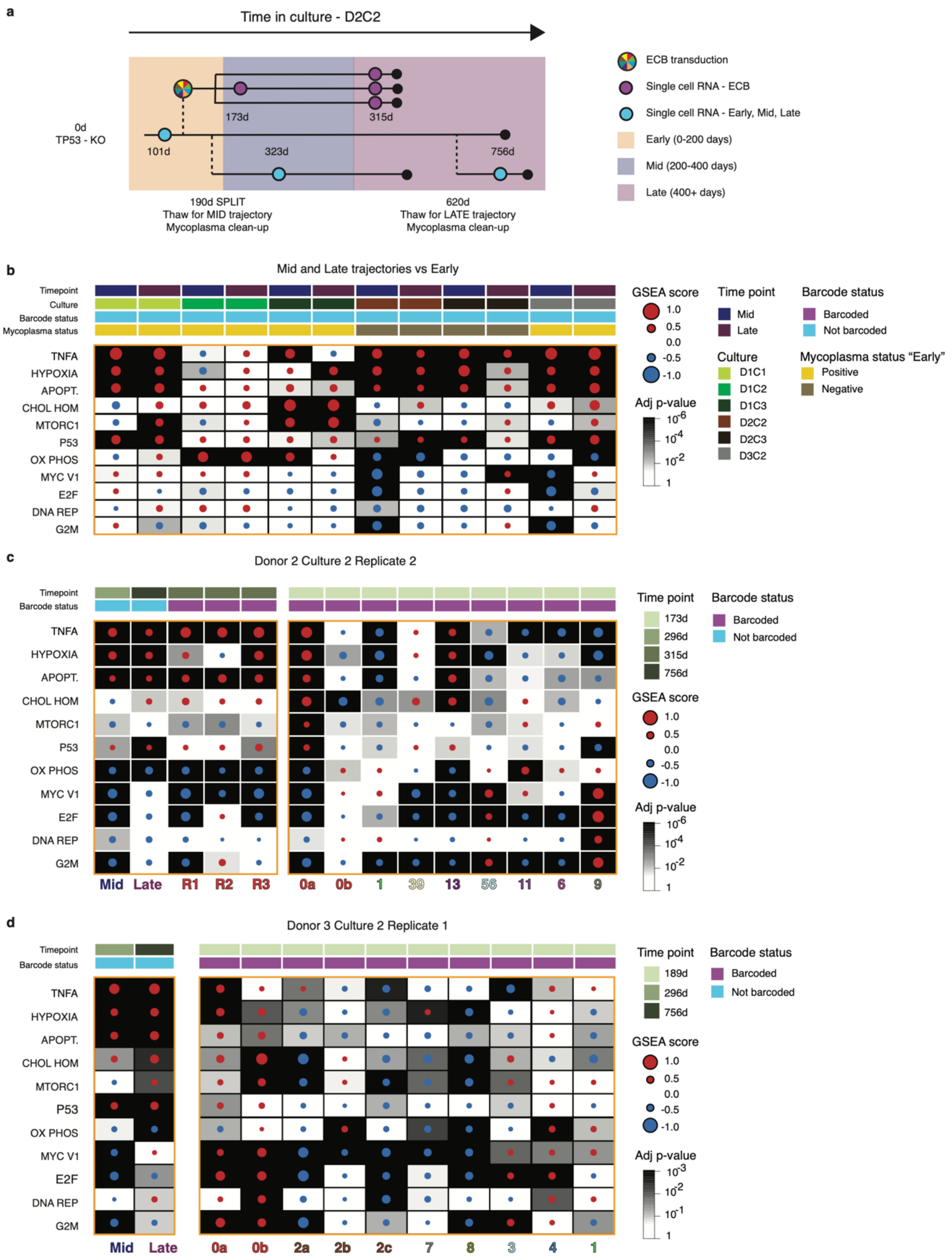

**Supplementary Figure 19. Convergent pathway alterations in the dominant clones across replicates within culture and across donors, irrespective of mycoplasma levels. a,** Schematic overview of the time course for a typical culture where single-cell RNA sequencing timepoints are highlighted. **b,** Gene Set Enrichment Analysis (GSEA) heatmap for MSigDB Hallmark gene sets showing the most significantly altered pathways for each culture comparing for each culture Mid and Late timepoints with Early time point (Kolmogorov-Smirnov statistic, Benjamini-Hochberg adjusted). GSEA score is indicated by dot size and colored according to the directionality of expression profiles (up, red; down, blue). **c,** GSEA heatmap from MSigDB Hallmark gene sets showing the most significantly altered pathways for specific subclones for Donor 2 Culture 2 Replicate 2 (D2C2R2) at day 173 (right; Kolmogorov-Smirnov statistic, Benjamini-Hochberg adjusted). GSEA score is colored according to the type of differential expression (up = red; down = blue). The heatmap on the left shows a comparison between subclone 0a and all other cells at a later time point (day 315) for all three replicates (R1-R3) as well as the comparison of D2C2 *Mid* (day 323) and *Late* (day 756) time points relative to *Early* (day 101), as in b. **d,** Same as c but for Donor 3 Culture 2 Replicate 1 (D3C2R1).

#### Supplementary Tables 1-9

**Supplementary Table 1.** List of oligonucleotides utilized in this study.

**Supplementary Table 2.** Cell passaging information for each donor and culture.

**Supplementary Table 3.** Summary of sWGS data, including time points, coverage and QC metrics, and genomic alterations.

**Supplementary Table 4.** Summary of WGS data, including time points, coverage and QC, as well as ploidy estimates.

**Supplementary Table 5.** Copy number alterations accompanying fishplot schematics.

**Supplementary Table 6.** Single-cell RNA sequencing results, including QC metrics, top differentially expressed genes and GSEA results.

**Supplementary Table 7**. Timing of clonal sweeps based on ECB sequencing data.

**Supplementary Table 8**. Subclone frequencies in fishplot schematics.

**Supplementary Table 9**. Single-cell RNA sequencing information for ECB subclones, top differentially expressed genes and GSEA results.
